# Proximity-Informed Graph Learning Defines Spatial Protein Communities for Tumor-Associated Proximity Antigen Discovery

**DOI:** 10.64898/2026.02.13.705629

**Authors:** Cody Scandore, Clare F. Malone, Christopher K. May, Anna K. de Regt, Jeff Guernsey, Hayley Ma, Noah Dephoure, Ben Setter, Rebecca A. Howell, Kendall R. Johnson, Carol L. Farr, Sophia Romero, Lydia Vignale, Tali Vittum, Emma Dawson, Tsadik Habtetsion, Francesca Nardi, Brian Woodruff, Martin Mathay, Julia Swanson, Quynh Ton, Payam E. Farahani, Robert W. Gene, Jason Misurelli, Zach Caldwell, Hengyu Xu, Michael Hornsby, Marc A. Gavin, Heath E. Klock, Ertan Eryilmaz, Pamela M. Holland, Scott A. Lesley, Rob C. Oslund, Olugbeminiyi O. Fadeyi

## Abstract

The spatial organization of membrane proteins is an underexplored dimension of cell-surface biology. Spatial proximity shapes cellular function and therapeutic targetability, yet efforts to identify tumor-associated antigens (TAAs) have largely focused on expression alone. Here, we developed an industrialized surface-protein proximity mapping workflow to interrogate TAAs within their membrane microenvironments. In the process, we generated 248 proximity maps across 12 receptor tyrosine kinases (RTKs) and 28 tumor cell systems. This proximity atlas enabled two advances: first, MetaMap, a correlation-based analytical framework that defines spatial protein communities and infers non-targeted proximal proteins from reproducible proximity signatures; and second, tumor-associated proximity antigens (TAPAs), a conceptual class of co-targets defined by disease-specific spatial proximity to TAAs rather than expression alone. Applying these proximity-derived relationships within a multimodal prioritization framework, we identified and validated an EGFR×CDCP1 TAA-TAPA pair that enhanced tumor cell killing across therapeutic modalities. By integrating spatial organization with multimodal data, this work expands the design space for precision-guided therapeutic strategies.

## Introduction

The cell surface is a densely packed and dynamic landscape in which the spatial proximity of membrane proteins shapes the activity and specificity of diverse biological processes, including immune synapse formation, epithelial junctional organization, and receptor-mediated signaling events^1-3^. Such proximity-driven arrangements are a defining feature of multiple protein classes that include immune receptors^4^, cytokine receptors^5^, multi-pass membrane proteins^6^, and receptor tyrosine kinases (RTKs)^7^, many of which assemble into localized signaling hubs^8^.

In cancer, the cell surface proteome undergoes extensive remodeling, including changes in protein abundance, trafficking, post-translational modifications, mutational status, and membrane organization^9-12^. Recent studies further demonstrate that tumor tissues are not merely heterogeneous but self-organize into spatially confined “neighborhoods” or sub-tumor microenvironments with conserved cellular, matrix, and signaling architectures^13^. These organizational features reflect alterations in surface protein spatial networks that can disrupt normal physiology and promote disease progression^10-12,14-16^. Importantly, because this spatial organization emerges at the protein level, it is not fully captured by gene or protein expression–based analyses, which quantify abundance but lack information on protein proximity. An approach that resolves surface protein organization is therefore required to contextualize disease-relevant surface proteins, such as tumor antigens, within their functional membrane environments and to more precisely inform the rational design of multi-target therapeutic strategies.

Despite the importance of spatial organization, scalable and industrialized approaches to map surface protein microenvironments with high resolution remain limited. We previously introduced a photocatalytic proximity-labeling technology that enables covalent labeling of proteins within a nanoscale radius to define surface protein neighborhoods^17-20^. Here, we extend this technology into an integrated, high-throughput workflow that enables proximity profiling at scale. This approach employs orthogonal labeling chemistries, including diazirine probes activated by iridium-based photocatalysts to generate short-lived carbenes^17^ and tyramide probes activated by riboflavin-based photocatalysts^20^ to generate tyrosyl radicals, enabling covalent labeling of proteins in close spatial proximity to capture local surface microenvironments. The workflow integrates standardized experimental procedures and cross-experiment normalization strategies to control technical variability and enable quantitative comparison of protein proximity across targets, cell systems, and tumor contexts.

Using this integrated membrane interactomics approach, we generated spatial maps of surface protein microenvironments across diverse cancer models. To extend these measurements and infer reproducible proximal relationships beyond directly targeted proteins, we developed an analytical framework that defines spatial protein communities and identifies conserved co-localization partners of non-targeted proteins, which we term MetaMap. Furthermore, to translate spatial organization into therapeutic prioritization, we also implemented a deep learning framework that leverages experimentally measured protein proximity and protein abundance to identify candidate proximity-associated pairs. To ensure biological relevance and therapeutic selectivity, predicted pairs were further prioritized using normal tissue expression, tumor-versus-normal expression differences, clinical proteomics data, and disease-relevant signaling features (**Figure 1**). These advances provide a general framework for contextualizing tumor-associated antigens (TAAs) within their spatial membrane neighborhoods and for prioritizing proximity-defined co-target pairs.

**Figure 1.**
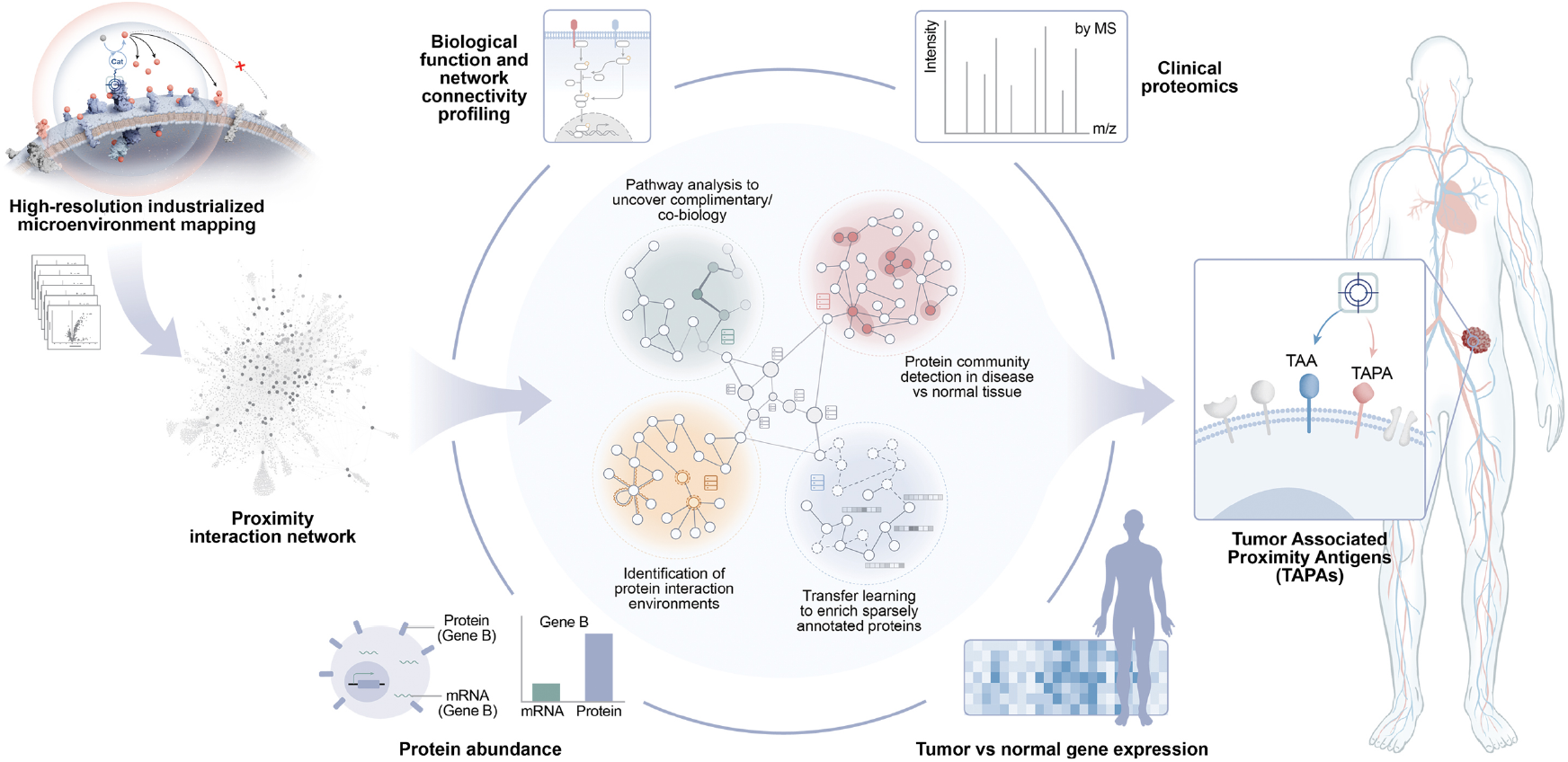
A data-driven framework for identifying Tumor-Associated Proximity Antigens (TAPAs). Industrialized microenvironment mapping (left) generates a global proximity network capturing local protein neighborhoods across diverse cell systems. Integrating these proximity signatures with protein abundance enables detection of disease-associated protein communities and nomination of candidate co-targets (center); pathway annotations, clinical proteomics, and tumor versus normal expression further contextualize these candidates for biological interpretation and prioritization. Together, these analyses nominate TAPAs: surface antigens defined not only by abundance but by disease-relevant proximity to tumor anchors (right), expanding therapeutic design toward spatial co-biology and actionable protein neighborhoods.

TAAs remain a foundational entry point for targeted cancer therapeutics, including antibody–drug conjugates (ADCs), T cell engagers (TCEs), and CAR-T cell therapies^21,22^. However, many clinically attractive TAAs exhibit incomplete tumor selectivity, with measurable expression in healthy tissues, such that engagement can produce dose-limiting toxicities and constrain therapeutic index. These limitations motivate strategies that move beyond single-antigen selection toward context-aware co-targeting, in which a second antigen is chosen not simply by expression, but by its tumor-specific spatial co-organization with the primary target on the cancer cell surface.

As an initial application of this approach, we focused on RTKs, a clinically important class of cell surface TAAs frequently dysregulated in cancer and amenable to extracellular engagement by multiple biologic therapeutic modalities, including ADC^23,24^, TCE^25^, and CAR-T cell^26^ therapies. Because RTKs form diverse homo- and heteromeric interaction networks that shape receptor neighborhood organization and signaling^7,16,27^, spatially informed co-target discovery offers a principled approach to enhance selectivity and efficacy across emerging RTK-directed ADC and TCE clinical efforts and supports exploration beyond the current landscape of tractable RTK targets (**Supplementary Figure 1**).

RTK interaction networks have been previously observed using affinity purification and intracellular enzyme-based proximity labeling approaches within a HEK293-based cellular background^28^. While this provided important insight into intracellular RTK signaling architecture, extracellular surface organization has remained less explored. To our knowledge, this study is the first to systematically resolve surface protein microenvironments of endogenously expressed RTKs across diverse RTK families and tumor contexts using extracellularly anchored proximity labeling, enabling quantitative comparison of RTK-associated proximity features across receptors and tumor cell systems.

Here, we introduce the concept of Tumor-Associated Proximity Antigens (TAPAs), defined as proteins that are not only co-expressed, but are also spatially co-enriched with canonical TAAs in the tumor microenvironment. By anchoring co-target discovery in protein proximity rather than expression alone, TAPAs provide an antigen-agnostic framework for identifying biologically coupled and therapeutically selective target pairs across the tumor surface proteome. Applying this framework to RTKs as an initial TAA family, we identify EGFR×CDCP1 as a TAA-TAPA pair that is spatially co-enriched across multiple tumor types and drives enhanced tumor cell killing when co-engaged using ADC and TCE modalities. Our integrated Membrane Interactomics (MInt) platform, which combines high-throughput surface microenvironment mapping (micromapping), the MetaMap analytical framework, and the TAPA concept establishes protein proximity as a generalizable and translatable design principle for precision multispecific therapeutics.

## Results

### Mapping TAA microenvironments using RTKs

To establish a scalable workflow for micromapping, we selected 12 representative RTKs spanning 10 structural families and profiled them across 28 cancer cell systems validated for tissue origin and surface antigen expression (**Supplementary Figures 2–4**). We generated a total of 248 micromaps using two orthogonal, visible-light-activated photocatalytic proximity-labeling chemistries: a mid-range riboflavin/biotin-tyramide (RFT/BT) and a short-range iridium/diazirine (Ir/Dz) system (**Figure 2a and Supplementary Figure 5**). These complementary chemistries differ in effective labeling radius and residue preference, providing robust coverage across distinct spatial scales and enhancing neighborhood resolution^29,30^. To enable mapping at this scale, we standardized cell input and implemented automated processing for key steps, including protein pulldown, digestion, and TMT labeling, with a KingFisher Apex system. Across the full mapping dataset, each intended RTK anchor target was consistently enriched with strong concordance observed between the two chemistries (**Supplementary Figure 6**), supporting a reproducible foundation for systematic microenvironment characterization.

**Figure 2.**
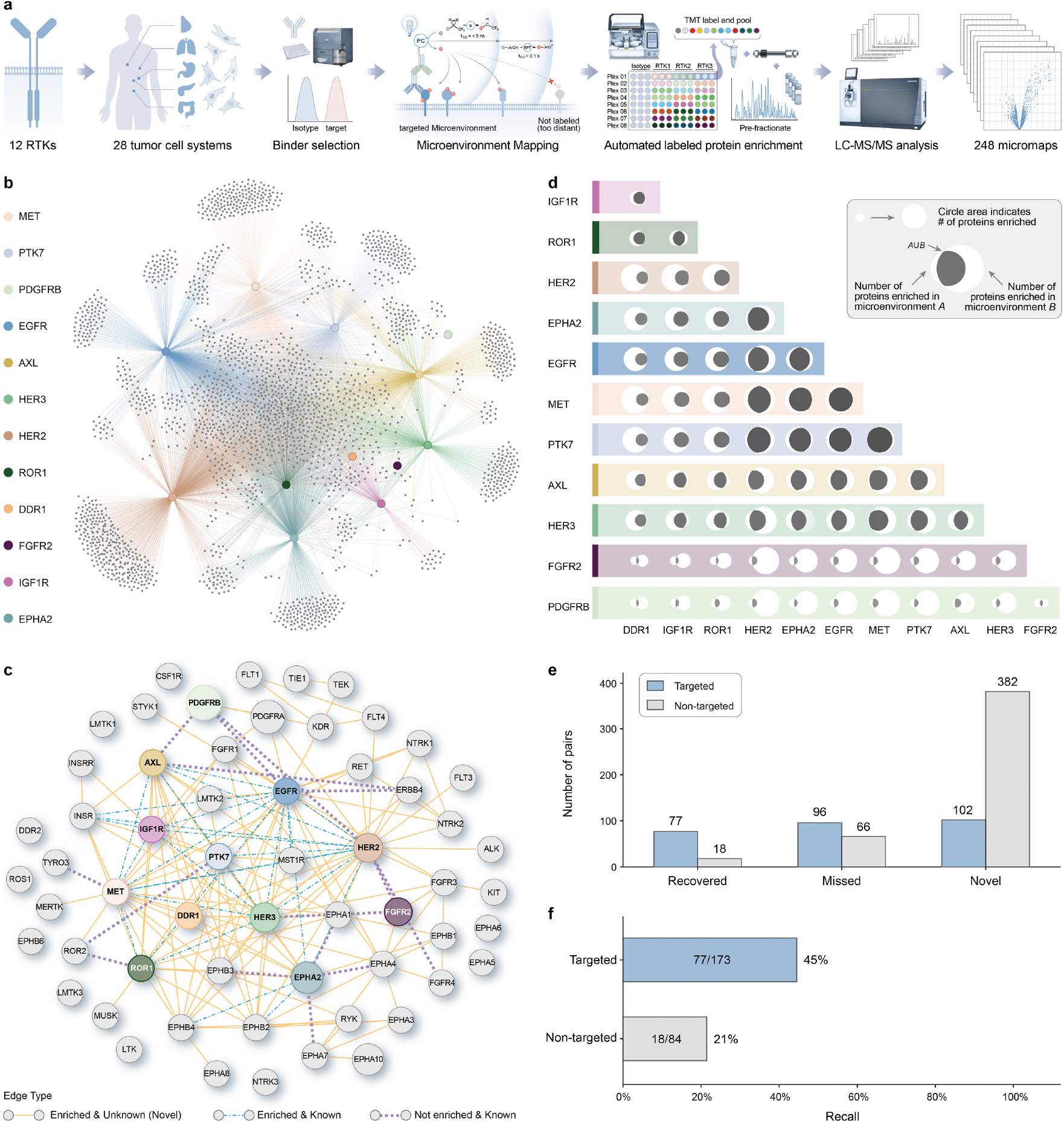
Scalable micromapping of RTK surface neighborhoods and benchmarking against the annotated protein interactome. **a)** Overview of the industrialized micromapping workflow used to generate 248 RTK-anchored proximity maps across 12 RTKs and diverse cancer cell systems, combining antibody-guided localization with two orthogonal visible-light proximity-labeling chemistries (mid-range RFT/BT and short-range Ir/Dz), automated enrichment and sample processing, TMT multiplexing, and LC-MS/MS quantification. **b)** Global proximity network summarizing enriched protein neighborhoods across the full dataset, revealing a densely connected landscape with receptor-specific structure. **c)** RTK interaction network highlighting recovered known^7^ RTK-RTK relationships alongside additional putative proximal partners. Targeted RTKs shown in color, non-targeted RTKs shown in gray. **d)** Pairwise neighborhood overlap across RTKs (e.g., Jaccard similarity) showing shared and distinct microenvironment composition, including high-overlap receptor pairs. **e)** Benchmarking of RTK-RTK proximity-derived associations against external protein-interaction resources (e.g., STRING, CORUM, BioGRID, IntAct), summarizing recovery of annotated interactions and the additional interaction space uniquely captured by proximity mapping. **f)** Recovery of annotated RTK-RTK interactions by proximity mapping. Bars indicate the fraction of database-curated interactions (from STRING, CORUM, BioGRID, and IntAct) detected in our dataset, stratified by whether RTKs were directly targeted for proximity labeling (77 of 173 known interactions recovered, 45%) or identified as interactors within non-targeted RTK microenvironments (18 of 84, 21%).

Resolving the complex architecture of the RTK microenvironment across hundreds of maps requires a rigorous computational approach to harmonize data and extract reproducible proximity structure. To address the inherent variability in large-scale mass spectrometry data, we developed a statistical processing pipeline that normalizes results across the full proximity dataset. We applied robust z-score normalization using the median absolute deviation (MAD) to t-statistics within each of the 248 datasets (**Supplementary Figure 7**). This approach equalizes score distributions across experiments with varying dynamic ranges, enabling fair cross-experiment comparisons. High-confidence protein associations were defined using a stringent threshold (MAD-normalized t >= 2.0) (**Supplementary Figure 8**), ensuring that the resulting interaction networks are driven by consistent, cross-experiment protein co-enrichment patterns, reflecting reproducible biological associations rather than experiment-specific variation.

Global connectivity analysis of our proximity datasets (**Figure 2b**) revealed both densely interconnected protein proximity landscapes between the mapped RTKs as well as receptor-specific neighborhoods. Our platform achieved high recall of the known RTK interactome, successfully identifying the majority of previously reported heterotypic interactors^7^ that includes EGFR-MET, EGFR-HER2, and HER2-HER3, and detected novel putative interactors such as EGFR-PTK7, MET-EPHB2, and HER3-MST1R (**Figure 2c**). Cross-referencing our proximity-derived RTK-RTK interaction network with publicly disclosed industry therapeutic pipelines reveals that clinical-stage exploration of heterotypic RTK-RTK co-targeting is highly limited. Only three RTK×RTK pairs (EGFR×MET, EGFR×HER3, and HER2×HER3) are currently in clinical-stage evaluation for dual targeting antibody-based modalities (**Supplementary Figure 9**) highlighting that the vast majority of RTK-RTK interactions identified by systematic proximity mapping remain clinically unexplored.

Analysis of RTK microenvironment similarity revealed distinct patterns of shared neighborhood composition; notably, pairs including HER2, EPHA2, EGFR, MET, and PTK7 exhibited the highest overlap, with Jaccard indices ranging from 0.4 to 0.5 (**Figure 2d and Supplementary Figure 10**). Importantly, elevated neighborhood similarity was confined to a defined subset of RTK pairs rather than broadly distributed across the receptor set, indicating that proximity overlap reflects unique environment organization rather than nonspecific membrane colocalization or surface abundance (**Supplementary Figure 11**). To further contextualize the RTK-RTK interaction landscape captured by proximity mapping, we compared our data more broadly with protein interaction databases that include STRING^31^, CORUM^32^, BioGRID^33^, and IntAct^34^ revealing substantial overlap with database captured RTK-RTK interactions as well as suggesting putative RTK-RTK proximity relationships not represented in existing annotations (**Figure 2e**). Notably, this overlap captured nearly half of the annotated interactions among directly targeted RTKs and extended to a substantial fraction (18 of 84) of annotated interactions involving RTKs that were not experimentally targeted, demonstrating recovery of RTK-RTK relationships beyond the mapped receptors (**Figure 2f**). Collectively, these analyses demonstrate that systematic proximity mapping captures structured and biologically meaningful interaction landscapes, providing confidence in established relationships while enabling exploration of previously unannotated protein associations.

### MetaMap resolves non-targeted proximity structure at scale

We next asked whether the statistical power of our large-scale proximity dataset could be leveraged to infer spatial relationships among non-targeted surface proteins. To address this, we computed pairwise Spearman rank correlations across the full set of proximity maps, capturing how protein proximity signatures co-vary across RTK anchors, cellular contexts, and experimental conditions. This correlation-based framework, which we term MetaMap, identifies high-confidence protein communities and non-targeted proximity associations based on consistent co-variation signatures, capturing shared spatial organization while reducing sensitivity to outliers and proteomic sampling noise (**Figure 3a**).

**Figure 3.**
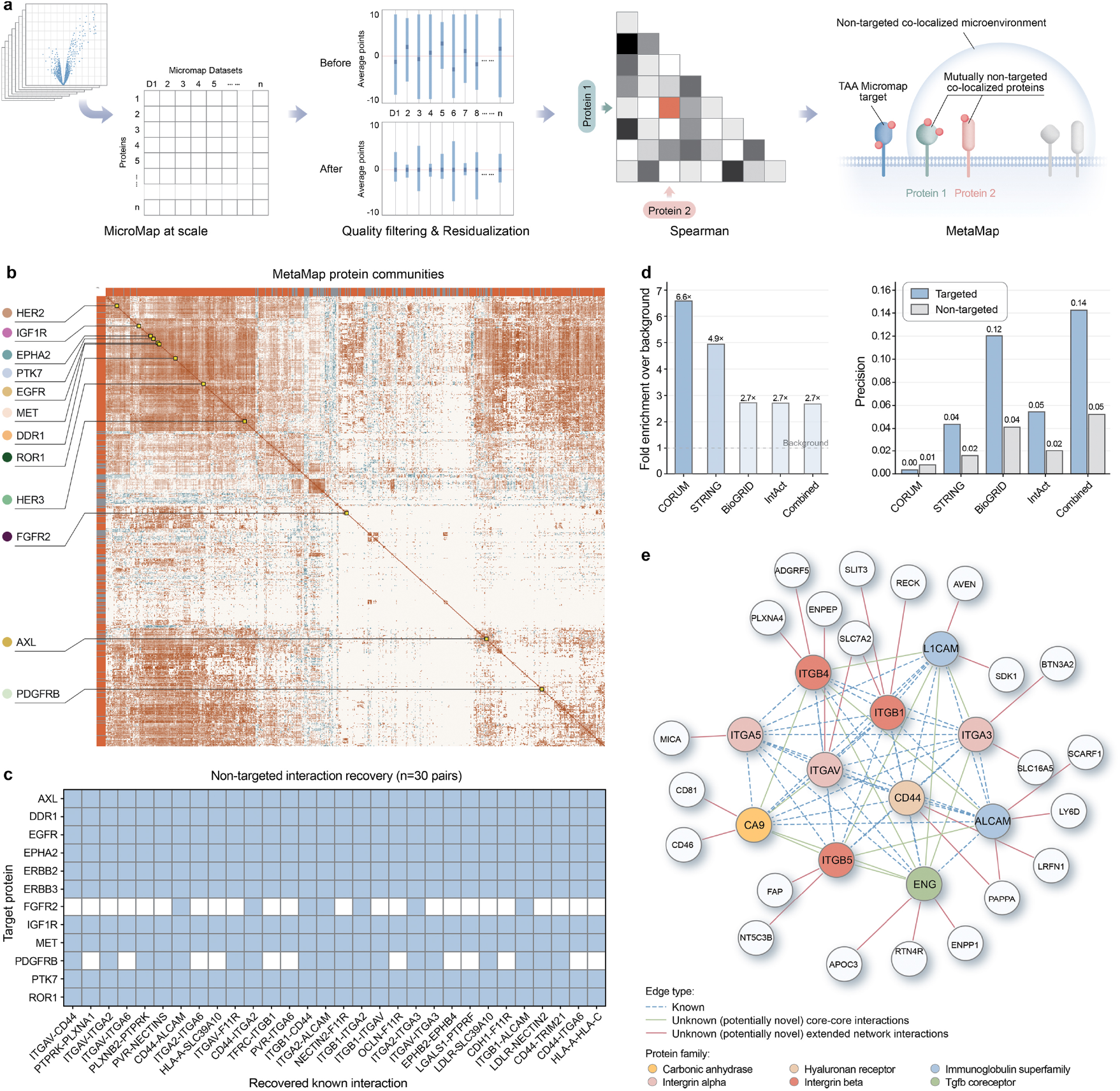
MetaMap infers spatial protein communities from large-scale micromaps. **a)** MetaMap workflow: micromaps are quality-filtered and normalized to reduce batch/efficiency effects, converted to protein-by-protein association matrices (Spearman rank correlation), and integrated to infer mutually proximal surface-protein relationships beyond directly targeted antigens. **b)** Global MetaMap correlation heat map across proteins. The matrix positions of the 12 mapped RTK targets are indicated at left and highlighted (yellow box). **c)** Recovery of known protein-protein interactions across targeted RTK microenvironments. Each column represents a curated interaction pair, and rows indicate the RTK-anchored experiment in which the interaction was detected. **d)** Benchmarking MetaMap associations against curated interaction databases. Left: fold enrichment over background for non-targeted interactions: CORUM 6.61×, STRING 4.92×, BioGRID 2.71×, IntAct 2.68×, Combined 2.67× (all p < 0.01, permutation test, n = 500). Right: precision across reference databases, with targeted interactions shown for comparison. **e)** Representative MetaMap community comprising integrin family members and cell adhesion molecules. Core proteins (colored) include integrin alpha/beta subunits; extended neighborhood shown as uncolored nodes. Edges indicate a high confidence correlation metric (Spearman rho > 0.4) and distinguish known, database annotated interactions (dashed) from potentially novel core-core (solid green) and extended network associations (solid red).

The resulting correlation structure reveals a highly organized surface interactome in which individual RTKs resolve into distinct and reproducible patterns within the correlation heatmap (**Figure 3b)**. Systematic benchmarking confirmed that many known direct and indirect interactions were reliably recovered across nearly all targeted microenvironments (**Figure 3c**). An exception was the FGFR-family anchors (e.g., FGFR2), which showed comparatively sparse recovery, likely reflecting more limited context sampling together with distinct membrane organization across FGFR members (**Supplementary Figure 12**). Crucially, when restricting the analysis to non-targeted interactions, MetaMap-derived associations were more likely to match curated interactions compared to size matched random protein pairs (**Figure 3d**).

We next applied community detection to the MetaMap association graph to identify a consistently enriched protein community (**Figure 3e**). One such community resolves into a coherent adhesion protein-enriched assembly, with integrin α/β subunits (ITGA/ITGB family members)^35^ forming a densely connected core, together with canonical adhesion molecules such as CD44, ALCAM, and L1CAM that engage in known interactions^36,37^. The recovery of these expected adhesion features supports that MetaMap captures reproducible membrane neighborhood structure from correlated proximity signatures.

Importantly, that same protein community also contains recurrent proximity-defined co-associations that are not annotated as established interactions. For example, MetaMap highlights extended links connecting FAP, CD81, RECK, and PAPPA to the core integrin network (**Figure 3e**). While these proteins have been implicated in adhesion-related biology ^38-41^, their inclusion here reflects reproducible spatial co-association, suggesting additional organization within integrin microenvironments beyond curated interaction databases. Together, these results establish MetaMap as a correlation-based framework that defines spatial protein communities and enables non-targeted inference of reproducible surface co-associations.

### Graph learning for prioritizing microenvironment co-pairs with therapeutic relevance

Having established that our large-scale proximity-derived micromapping approach captures reproducible and structured protein co-associations across the tumor cell surface, we next focused on translating these spatial patterns into therapeutic prioritization. We sought to identify co-target pairs defined by spatial organization and biological context rather than surface abundance alone. EGFR serves as a stringent reference antigen for this analysis, as its broad surface distribution and extensive therapeutic history make it a suitable test case for evaluating proximity-informed co-target prioritization.

To operationalize this prioritization, we applied machine learning-based graph models to directly labeled proximity data derived from all our RTK-anchored micromaps. We constructed a homogenous protein graph per experiment that integrates experimentally measured proximity co-enrichment with protein abundance (**Figure 4a**). Using these experimental representations, we trained three model architectures, a naïve structure only model (Node2Vec)^42^, a variational graph autoencoder (VGAE)^43^, and a graph attention network (GAT)^44^, to learn latent structure and generate ranked EGFR-associated co-target predictions. Comparison of model outputs across high-confidence EGFR-associated candidates revealed both shared and architecture-specific prioritization patterns (**Figure 4b**). GAT and VGAE consistently recovered key EGFR signaling components, including HER2, HER3, and MET (**Figure 4b**), supporting the biological validity of their learned representations. Notably, CDCP1 was also highly ranked in both models, despite not being a canonical EGFR network member, highlighting its identification as a proximity-defined association. By comparison, Node2Vec produced noisier predictions and was less efficient at recovering known protein interactions, reflecting its dependence on positional graph embeddings without access to the proximity and abundance features that inform GAT and VGAE.

**Figure 4.**
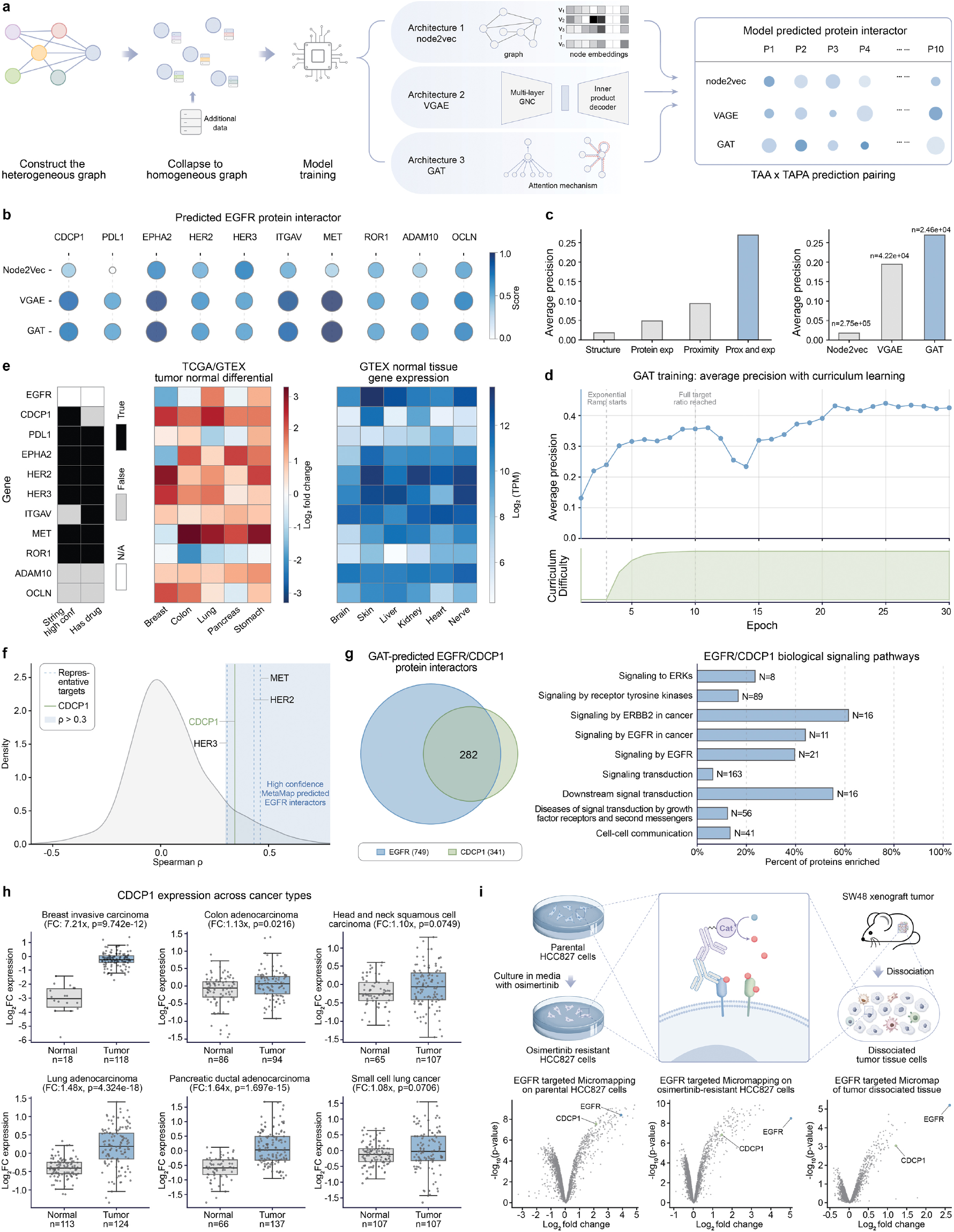
Graph-based learning prioritizes CDCP1 as an EGFR-associated, tumor-selective partner. **a)** Overview of graph-based machine learning workflow. Heterogeneous graphs integrating protein proximity and protein expression data are collapsed into homogeneous protein graphs, then used to train three architectures (Node2Vec, VGAE, GAT) that output ranked protein interactor predictions for TAA–TAPA pair prioritization. **b)** Predicted EGFR interactor rankings across all three models. Dot size and color indicate normalized prediction score. **c)** Left: Feature ablation analysis of the GAT model showing average precision when trained using graph structure alone (Structure), graph structure with protein expression (Protein exp), graph structure with proximity features (Proximity), or the full feature set combining proximity and protein expression (Prox and exp). Right: Cross-model performance comparison with all features enabled; n indicates number of predictions per model. **d)** GAT training dynamics under curriculum learning. Average precision (top) across training epochs, with dashed lines indicating curriculum transitions. Curriculum difficulty (bottom) shows progressive introduction of harder negative samples. **e)** Multimodal annotation of ten high confidence EGFR interactors predicted by graph models. Left: STRING high-confidence EGFR interaction status and clinical-stage therapeutic existence. Center: Log_2_ tumor-versus-normal expression differential (TCGA/GTEx) across five indications. Right: GTEx normal tissue expression. **f)** MetaMap-derived Spearman correlation coefficients for EGFR across all profiled proteins. **g)** Left: Overlap of GAT-predicted protein interactors between EGFR and CDCP1. Right: Reactome pathway enrichment of shared interactors. **h)** CDCP1 protein expression (log_2_ fold change) in tumor versus matched normal tissue across six cancer types from published clinical proteomic datasets^62-67^. **i)** Volcano plots of EGFR-anchored microenvironment maps in parental HCC827 cells, osimertinib-resistant HCC827 cells, and dissociated SW48 xenograft tumor tissue (n = 3 experiments per condition).

To understand the contributions of individual data modalities to model performance, we performed systematic feature ablation analyses (**Figure 4c** and **Supplementary Figure 13**). Removal of either protein proximity or protein expression reduced model precision, whereas integrating both features consistently yielded the highest performance. Of note, proximity-derived information provided independent, non-redundant signal beyond protein expression or graph structure (i.e., annotated protein interactions) alone. Training dynamics revealed rapid early learning, with GAT models achieving the majority of performance gains within the first several epochs under a curriculum schedule that progressively introduced more challenging prediction tasks (**Figure 4d**). This early convergence likely reflects two complementary factors: the attention mechanism efficiently identifies informative neighborhood structure by learning to weight edges according to their predictive relevance, and because input graphs are grounded in experimentally measured proximity, the signal-to-noise ratio is high from initialization.

Building on these results, we integrated multiple independent annotation layers to contextualize high-confidence EGFR-associated candidates (**Figure 4e**), including evidence of EGFR association, clinical stage therapeutic targeting, and expression context. In this analysis, CDCP1 is distinguished as a proximity-defined EGFR-associated non-RTK, in contrast to canonical EGFR network components such as MET, HER2, and HER3 that dominate existing co-targeting strategies.

We next examined the underlying CDCP1 proximity and biological data in greater detail to contextualize its relationship with EGFR. First, analysis of MetaMap correlation networks revealed strong and reproducible spatial co-enrichment between EGFR and CDCP1 across independent proximity-derived datasets (**Figure 4f**), supporting CDCP1 as a proximity-defined EGFR-associated protein. To further deepen this proximity analysis, we generated 38 CDCP1-anchored micromaps spanning 19 tumor cell systems using both proximity-labeling chemistries (**Supplementary Figure 14**). Across these diverse cellular contexts, CDCP1 exhibited robust co-enrichment of EGFR, mirroring reciprocal EGFR-anchored maps (**Supplementary Figure 15**) and remaining detectable across independent micromap targeted binders (**Supplementary Figure 16**). Notably, this spatial proximity was attenuated in normal cells in which both EGFR and CDCP1 are expressed, indicating context-dependent spatial organization rather than simple co-expression (**Supplementary Figure 17**).

In addition, the GAT model trained on the full RTK micromapping dataset predicts EGFR and CDCP1 interactors with overlapping protein neighborhoods and convergent signaling features, supporting their participation in shared biological signaling environments (**Figure 4g**). Consistent with this shared organization, EGFR- and CDCP1-anchored micromaps independently enriched a conserved EGFR-signaling module comprising GRB2, GAB1, SHC1, and SOS1^45^ (**Supplementary Figure 18**). CDCP1 microenvironment maps further highlighted proximity to proteins associated with known CDCP1 roles in tumor progression and TKI resistance, including SRC-family kinases and PRKCδ/θ^46-48^ (**Supplementary Figure 18**). Beyond these previously characterized interactions, CDCP1 mapping identified additional proximal partners including PTPN12^49^, INPPL1^50^, and ARHGEF5/ASAP-family^51-54^ GTPase regulators with roles in cell migration, invasion, and metastasis (**Supplementary Figure 18**). Together, these data independently corroborate prior mechanistic observations of EGFR–CDCP1 signaling relationships^48,55^ by defining their association across diverse tumor contexts and spatial microenvironments.

Analysis of publicly available clinical proteomic datasets revealed elevated CDCP1 protein abundance in tumors relative to matched normal tissues across multiple cancer types (**Figure 4h**), a pattern independently supported by internal proteomic analyses (**Supplementary Figure 19**). These observations align with prior reports describing elevated CDCP1 expression across diverse tumor types, including settings associated with more aggressive disease^55-58^.

Building on these spatial and network-level observations, prior mechanistic studies have shown that higher CDCP1 expression is associated with poorer progression-free survival in EGFR-mutant patients and increased CDCP1 expression in tyrosine kinase inhibitor (TKI)-resistant contexts, including following osimertinib treatment^47,55,59^. To determine whether EGFR-CDCP1 spatial association is preserved under therapeutic pressure, we performed EGFR proximity mapping in parental and osimertinib-resistant HCC827 cellular states and observed that CDCP1 remains consistently enriched **(Figure 4i**). In a tumor tissue context, EGFR proximity mapping in dissociated tumor cells derived from xenograft tissue demonstrated continued CDCP1 enrichment (**Figure 4i**), indicating preservation of this spatial relationship within native tumor material. Finally, CDCP1 proximity mapping performed in the presence of EGF revealed increased EGFR-CDCP1 spatial association, consistent with prior studies reporting that EGF stimulation promotes CDCP1 upregulation and clustering^60,61^ (**Supplementary Figure 20**).

Collectively, these findings support a coherent, context-dependent spatial and biological relationship between EGFR and CDCP1 that extends beyond expression alone and provides a foundation for subsequent functional evaluation of CDCP1 as an EGFR-associated TAPA.

### Functional evaluation of EGFR×CDCP1 as a therapeutic co-targeting pair

Following prioritization of CDCP1 as a TAPA partner for EGFR, we investigated whether co-targeting this pair could enhance tumor-directed cytotoxicity using bispecific antibody–drug conjugates (ADCs). Because CDCP1 exhibits more restricted normal-tissue expression than EGFR (**Figure 4e**), bispecific ADCs were designed with lower-affinity engagement of EGFR and higher-affinity engagement of CDCP1 to favor productive cis-engagement, internalization, and payload delivery in cells where both receptors are co-expressed in close spatial proximity. This configuration implements a logic-gated co-targeting mechanism that biases activity toward co-expressing tumor cells while limiting uptake and cytotoxicity in normal cell systems.

Using these design principles, we assembled a panel of EGFR×CDCP1 bispecific antibody tool molecules for ADC evaluation and selected a representative construct to assess proximity-based co-targeting (**Supplementary Figure 21**). Because ADC efficacy depends on internalization, we first assessed cellular uptake of the unconjugated EGFR×CDCP1 bispecific antibody across cancer cell lines spanning a range of EGFR and CDCP1 expression levels, alongside matched monovalent EGFR-only and CDCP1-only controls (**Figure 5a and Supplementary Figure 21**). The unconjugated bispecific antibody exhibited enhanced internalization relative to either monospecific control (**Figure 5b**), consistent with cooperative cis-engagement of the two receptors.

**Figure 5.**
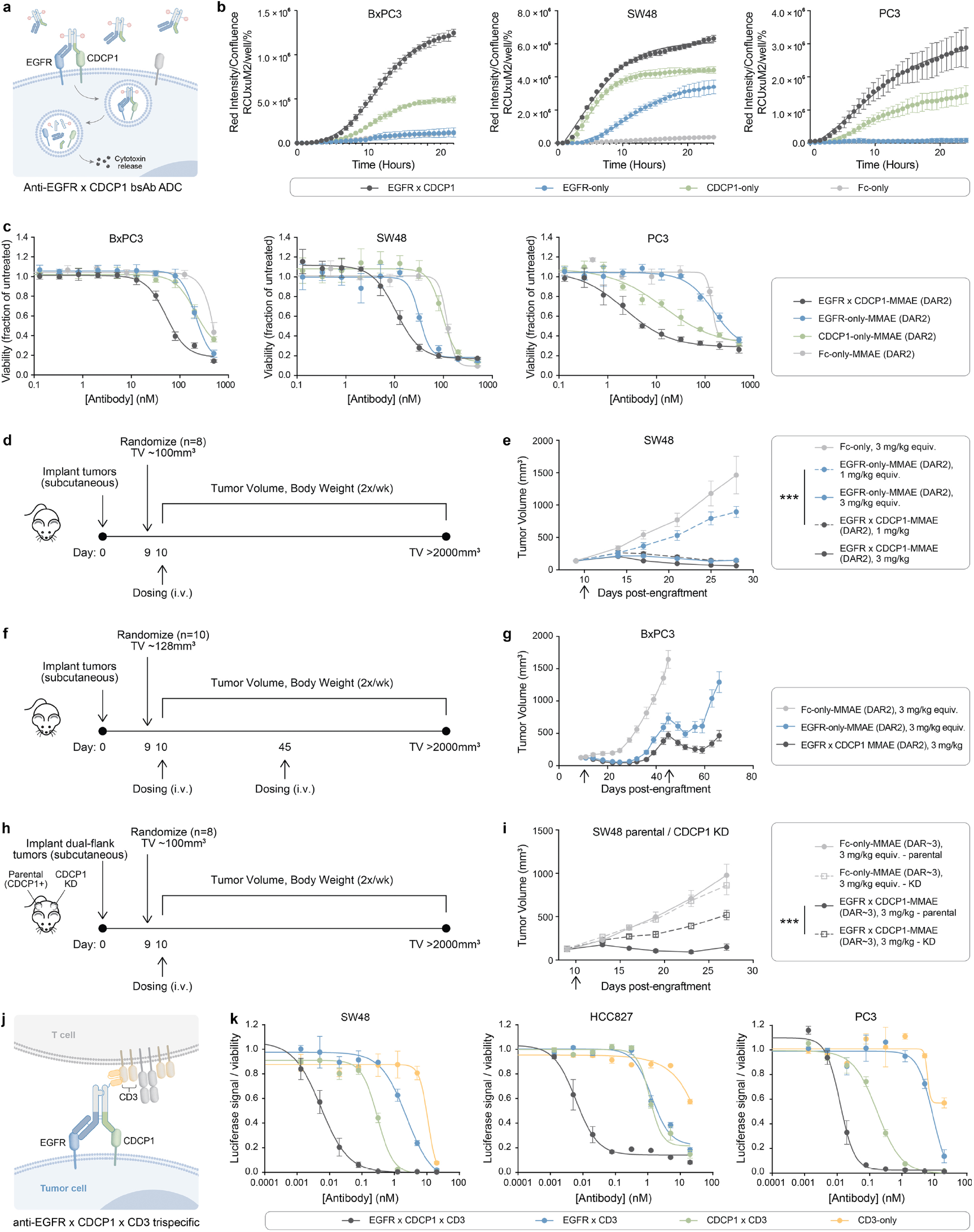
Functional evaluation of EGFR×CDCP1 co-targeting through ADCs and TCEs. **a)** Mechanism of action of a bispecific EGFR×CDCP1 ADC. Target binding induces internalization, lysosomal degradation, and cytotoxic payload release **b)** Live-cell imaging of internalization of unconjugated EGFR×CDCP1 bispecific and controls in EGFR/CDCP1expressing cancer cell lines. Total integrated intensity relative to phase confluence (indicative of internalization) is plotted over time. Data are mean ±SD (n=3). **c)** Relative viability of cancer cell lines treated with EGFR×CDCP1 ADC or matched DAR controls (**Supplementary Figure 21**) measured by CellTiter-Glo and normalized to untreated cells. Data are mean ±SD (n=4). **d)** Study design for SW48 xenograft. **e)** Tumor growth of SW48 xenografts following treatment with indicated test articles. Data are mean tumor volume ±SEM (n=8); arrows denote dosing day. Two-way ANOVA with Tukey’s multiple comparison; ***, p<0.001. **f)** Study design for BxPC3 xenograft. **g)** Tumor growth of BxPC3 xenograft tumors following treatment, with an additional dose on Day 45 (arrow). Data are mean ±SEM (n=10). **h)** Study design for SW48 dual-flank xenograft. **i)** Tumor growth of SW48 parental and CDCP1-knockdown (KD) xenograft tumors treated with indicated agents. Data are mean ±SEM (n=8). Two-way ANOVA with Tukey’s all groups comparison; ***, p<0.001. **j)** Mechanism of action of a trispecific EGFR×CDCP1×CD3 TCE. Bridging antigen-expressing tumor cells and CD3+ T-cells induces cytotoxicity. **k)** T cell dependent cellular cytotoxicity of trispecific EGFR×CDCP1×CD3 TCEs and controls. Indicated cancer cell lines were luciferase labeled and incubated with PBMCs and antibodies. Viability was measured by luciferase signal. Data are mean ±SD (n=3).

We next evaluated internalization of the EGFR×CDCP1 bispecific in primary cervical and bronchial/tracheal epithelial cells expressing EGFR and CDCP1 at levels comparable to tumor cells (**Supplementary Figure 22a**). In contrast to cancer cells, minimal internalization was observed in primary cells (**Supplementary Figure 22b**), supporting tumor-biased uptake driven by spatial co-localization rather than expression alone.

To assess functional cytotoxicity, antibodies were conjugated with a valine-citrulline-linked monomethyl auristatin E (vcMMAE) linker-payload using two ADC formats that included a site-specific engineered cysteine conjugate at a drug-to-antibody ratio (DAR) of 2 or an interchain disulfide conjugate at a higher DAR (DAR∼3) (unless otherwise indicated, in vitro and standard xenograft studies were performed using the site-specific DAR2 format).

The EGFR×CDCP1-MMAE (DAR2) ADC induced dose-dependent killing across multiple cancer cell lines and demonstrated superior potency relative to EGFR-only-MMAE (DAR2) or CDCP1-only-MMAE (DAR2) monovalent ADC controls (**Figure 5c**). Consistent with reduced internalization, minimal cytotoxicity was observed in primary epithelial cells treated with either the bispecific or monospecific ADCs (**Supplementary Figure 22c**). Across these comparisons, the bispecific ADC exhibited up to two orders of magnitude greater cytotoxic potency in cancer cell lines compared with primary epithelial cells, and the greater potency of the bispecific over the monovalent controls was abolished in the primary cells.

We next investigated the in vivo activity of the EGFR×CDCP1 bispecific ADC in SW48 colorectal and BxPC3 pancreatic cancer xenograft models, both of which express EGFR and CDCP1 (**Figures 5d and 5f and Supplementary Figures 22d and 22g**). In the SW48 model, a single 3 mg/kg dose of the bispecific ADC produced >100% tumor growth inhibition (TGI) relative to the Fc-only control, comparable to a molar-matched EGFR-only control ADC (**Figure 5e**). At 1 mg/kg, the bispecific ADC maintained robust antitumor activity (100% TGI), whereas the EGFR-only ADC showed substantially reduced efficacy (43% TGI, **Figure 5e**). Overall, the EGFR×CDCP1 ADC achieved improved overall survival relative to the EGFR-only ADC without observable body-weight loss (**Supplementary Figures 22e and 22f**). In the BxPC3 model, the bispecific ADC again matched EGFR-only ADC activity at 3 mg/kg and, upon re-dosing after tumor regrowth, induced additional tumor growth inhibition and improved overall survival relative to the EGFR-only ADC (**Figure 5g, Supplementary Figures 22h and 22i**).

To directly test whether in vivo activity depends on dual-antigen engagement, we next implemented a dual-flank SW48 xenograft model in which mice carried parental tumors on one flank and matched CDCP1-knockdown (KD) tumors on the contralateral flank (**Figure 5h and Supplementary Figure 23**). For this side-by-side comparison within the same animals, we used an interchain disulfide EGFR×CDCP1-MMAE ADC (DAR ∼3) as a functional tool reagent, with the same binding arms as the site-specific DAR2 construct. Animals were treated with a single i.v. dose of EGFR×CDCP1-MMAE (DAR∼3) or Fc-only-MMAE (DAR∼3) control (3 mg/kg equivalent) using the dosing schedule shown (**Figure 5h**). The bispecific ADC produced marked growth suppression of parental tumors (97% TGI), whereas efficacy was substantially attenuated in the CDCP1-KD flank (46% TGI, **Figure 5i**). In contrast, the Fc-only control ADC showed comparable tumor progression in both parental and CDCP1-KD tumors. These results provide in vivo evidence that the antitumor activity of EGFR×CDCP1 co-targeting is conditioned on CDCP1 presence and supports a dual-antigen-biased delivery mechanism rather than EGFR targeting alone.

To assess whether EGFR×CDCP1 could be extended across therapeutic modalities, we evaluated this pair in a trispecific T cell engager (TCE) format using different EGFR and CDCP1 binding arms in a tool molecule designed for simultaneous engagement of EGFR, CDCP1, and CD3 (**Figure 5j and Supplementary Figure 21**). We reasoned that tumor-specific spatial proximity between EGFR and CDCP1 would promote more efficient immune synapse formation when both antigens are co-engaged on the same cell, while limiting redirection toward normal cells expressing only one antigen^68^. In T-cell-dependent cellular cytotoxicity assays, the EGFR×CDCP1×CD3 trispecific TCE elicited enhanced tumor cell killing relative to EGFR×CD3 or CDCP1×CD3 controls across all three tumor cell lines tested, achieving >100-fold increased potency in one model (**Figure 5k**).

Collectively, these results demonstrate that TAPA-guided, proximity-based co-targeting of EGFR and CDCP1 can be functionally translated across therapeutic modalities, enabling logic-gated tumor selectivity and enhanced antitumor activity. These insights are now informing advancement of an EGFR×CDCP1 bispecific ADC toward the clinic.

## Discussion

This study demonstrates that scalable surface microenvironment mapping, implemented through our MInt platform, can resolve disease-relevant spatial organization on the tumor cell surface. By integrating a standardized micromapping workflow with correlation-based analytics (MetaMap), we inferred reproducible protein co-residence patterns across tumor cell surface environments. Building on these spatial relationships, we integrated proximity information with complementary molecular features to define TAPAs, a class of co-targets that translate spatial context into actionable co-targeting strategies.

As a proof of concept, we identified CDCP1 as a TAPA spatially co-enriched with EGFR across multiple cancer cell systems. Although EGFR is broadly expressed in normal tissues, its consistent spatial coupling with CDCP1, an antigen with more restricted normal tissue distribution (**Figure 4e**), revealed a rational co-targeting axis not apparent from expression-based analyses alone. Dual engagement using bispecific ADC or trispecific TCE modalities enhanced tumor cell cytotoxicity and in vivo tumor control relative to single-antigen targeting, supporting spatially informed co-targeting as a strategy to improve therapeutic index.

Several limitations warrant emphasis. While this initial effort was intentionally broad, sampling multiple RTKs across diverse tumor cell systems, it does not fully resolve which spatial relationships are uniquely enriched in specific disease contexts versus those reflecting conserved aspects of membrane organization. Rather than implying absolute tumor specificity, TAPAs are best viewed as spatial relationships whose enrichment and functional relevance vary across tumor lineage, genetic state, and treatment context. By design, spatial co-enrichment captures both direct molecular interactions and shared compartmental residence, and its interpretation may be influenced by antibody-dependent factors such as epitope accessibility or antigen density. Furthermore, proximity labeling reflects interactions within a finite labeling radius and temporal window, limiting resolution of fine-grained spatial organization as well as dynamic or transient proximity changes associated with receptor activation, trafficking, or acute drug exposure.

An additional consideration is that much of the spatial mapping was performed in tumor cell systems that lack full microenvironmental complexity, and interactions involving stromal or immune components are not comprehensively captured in the current framework. Extending this approach to matched normal tissues, treatment-altered states, and primary tumor material will enable more direct comparisons of tumor-restricted versus conserved spatial architectures than are possible using expression- or genetics-based approaches alone and could serve as potential avenues for future research. Such comparisons will require careful benchmarking given the lack of consensus across existing protein interaction databases and the absence of a unified ground truth.

Despite these limitations, graph-based integration allows us to extract knowledge from protein proximity that exceeds the sum of its parts. This approach transforms static interaction lists into a dynamic, scalable scaffold that becomes increasingly predictive as it expands across new targets and cell lines. We present this dataset as a starting point, offering a layer of spatial resolution that complements and enhances ongoing work in orthogonal fields like spatial transcriptomics and functional genomics.

Together, these results position spatial co-enrichment as a biologically grounded organizing principle for next-generation multispecific therapeutic design. Although applied here to RTKs, the principles underlying proximity-defined co-targeting are broadly applicable to surface protein targets across disease biology, including inflammatory, metabolic, and neurological contexts. As proximity maps expand across tissues, cell states, and disease settings, we anticipate that this integrated experimental and computational framework will enable co-target prioritization and advancement of proximity-guided therapeutic interventions.

## Author contributions

O.O.F. and R.C.O. conceived of the work. C.K.M., N.D., R.A.H., K.R.J., L.V., T.V., and E.D. designed and performed micromapping and proteomic experiments. C.S. and J.G. designed and executed bioinformatic and machine learning analysis. C.S., C.F.M., J.G., F.N., R.C.O., and O.O.F. conceived and developed the multimodal target prioritization workflow. B.S., C.L.F., B.W., M.M., J.S., Q.T., P.E.F., R.W.G., J.M., Z.C., H.X., M.H., and H.E.K. designed, engineered, characterized, and generated ADCs and TCEs. C.F.M., A.K.D., H.M., S.R., T.H., and M.A.G. designed, executed and performed in vitro and in vivo ADC and/or TCE evaluation. E.E., P.M.H., S.A.L. provided insight and direction for experimental design. O.O.F., R.C.O., C.S. wrote the manuscript with input from all authors.

## Competing Interest Statement

All authors were/are employed by InduPro during the experimental planning, execution and/or preparation of this manuscript.

