## Supplementary Information for "Proximity-Informed Graph Learning Defines Spatial Protein Communities for Tumor-Associated Proximity Antigen Discovery"

### Supplementary Figures

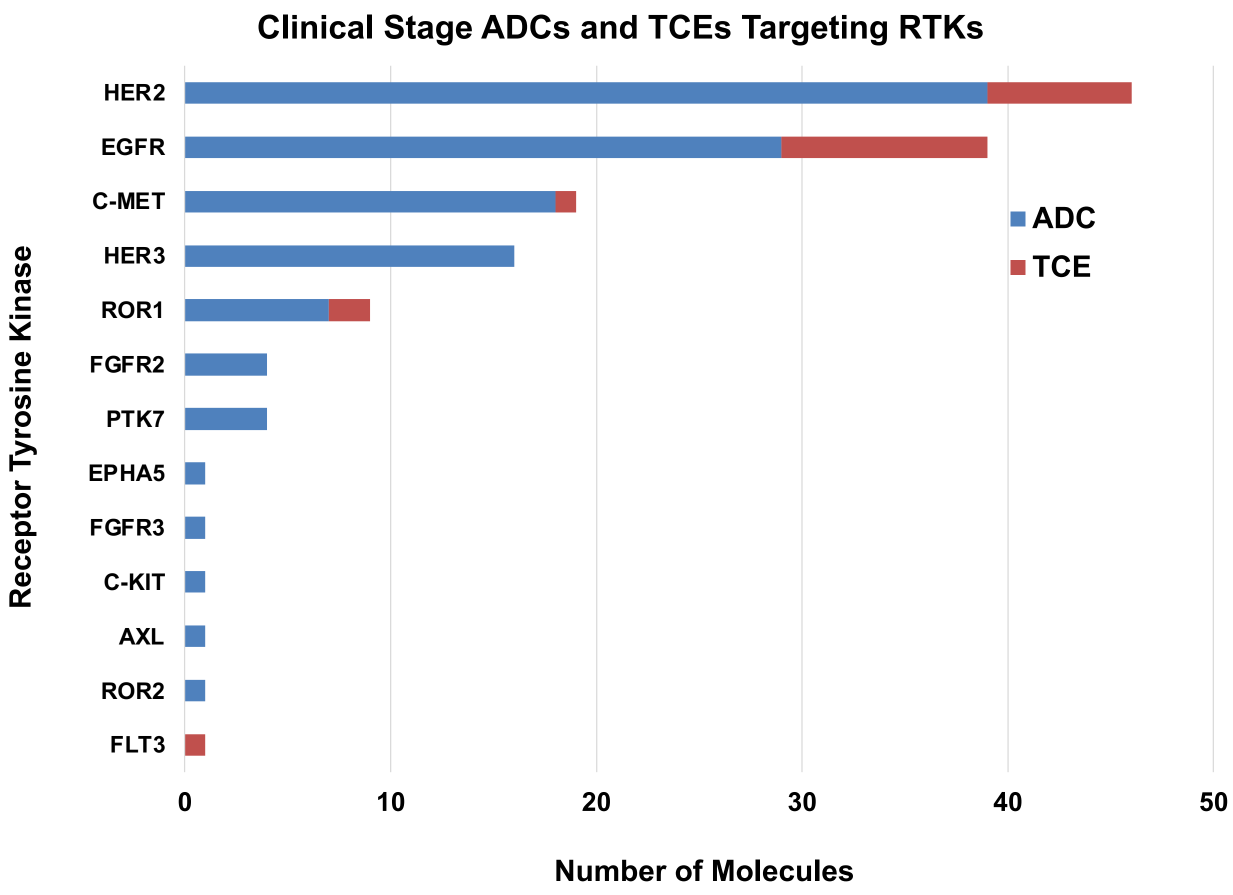

**Supplementary Figure 1. Clinical-stage antibody drug conjugates (ADCs) and T cell engagers (TCEs) targeting receptor tyrosine kinases (RTKs).** Bar chart highlighting the number of ADCs (blue) and TCEs (red) currently in clinical development for each receptor tyrosine kinase (RTK) target. Data were compiled and analyzed using Synapse by Patsnap.

**a**

| | | Photocatalyst | | | | | | | | | | |
|---|---|---|---|---|---|---|---|---|---|---|---|---|
| | | Ir | RFT | Ir | RFT | Ir | RFT | Ir | RFT | Ir | RFT | |
| RTK family | RTK target | Bowel | Bowel | Breast | Breast | Gastric | Gastric | Lung | Lung | Pancreas | Pancreas | Total |
| ErbB | EGFR | 4 | 4 | 4 | 3 | 2 | 2 | 9 | 9 | 2 | 2 | **41** |
| ErbB | HER2 | 2 | 2 | 2 | 2 | 2 | 2 | 7 | 7 | 2 | 2 | **30** |
| ErbB | HER3 | 2 | 2 | 2 | 3 | 3 | 3 | 6 | 5 | 2 | 2 | **30** |
| MET | MET | 4 | 4 | 2 | 2 | 2 | 2 | 10 | 10 | 2 | 2 | **40** |
| TAM | AXL | 2 | 2 | 2 | 2 | 1 | 1 | 7 | 7 | 2 | 2 | **28** |
| PTK7 | PTK7 | 2 | 2 | 2 | 2 | 2 | 2 | 6 | 6 | 2 | 2 | **28** |
| ROR | ROR1 | 2 | 1 | 2 | 2 | 2 | 2 | 5 | 5 | 2 | 2 | **25** |
| Ins | IGF1R | | | | | | | 4 | 4 | | | **8** |
| FGF | FGFR2A | | | | | | | 1 | 1 | | | **2** |
| Eph | EPHA2 | | | | | | | 5 | 5 | | | **10** |
| PDGF | PDGFRB | | | | | | | 1 | 1 | | | **2** |
| DDR | DDR1 | | | | | | | 2 | 2 | | | **4** |
| | | | | | | | | | | | | **Total: 248** |

**b**

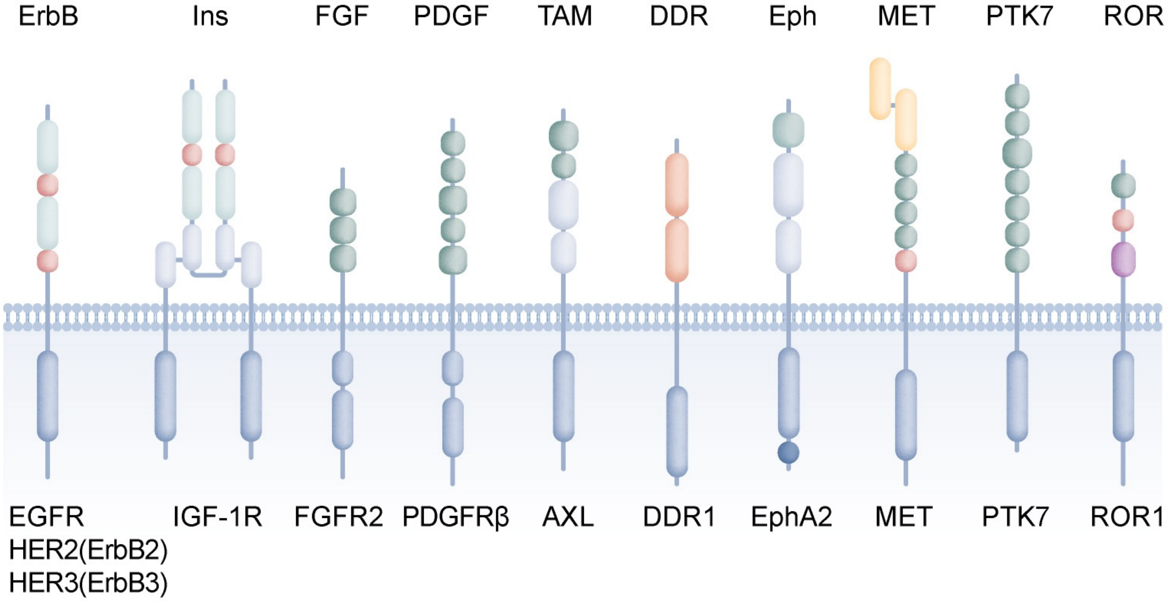

**Supplementary Figure 2. RTKs profiled by micromapping. a)** Table summarizing the number of micromapping experiments performed for each RTK target and the photocatalyst chemistry used. RFT denotes riboflavin tetraacetate–based photocatalysis, and Ir denotes iridium-based photocatalysis. Each micromap corresponds to an independent mapping experiment capturing proximal protein environments. **b)** Schematic representation of the RTKs, grouped by superfamily, that are profiled in this study. Each receptor's extracellular and intracellular domains are depicted based on canonical domain architecture.

**a**

| RTK(s) | Cell line | Lineage | Subtype info |
|---|---|---|---|
| AXL, DDR1, EGFR, EPHA2, HER2, HER3, IGF1R, MET, PDGFRA, PTK7, ROR1 | A549 | Lung | Lung (NSCLC) |
| EGFR, HER2, HER3, MET, PTK7, ROR1 | AGS | Gastric | Gastric |
| AXL, EGFR | BT549 | Breast | Breast (TNBC) |
| AXL, EGFR, HER2, HER3, MET, PTK7, ROR1 | BXPC3 | Pancreas | Pancreas |
| EGFR, HER2, HER3, MET, PTK7, ROR1 | CACO2 | Bowel | Bowel (MSS) |
| AXL, EGFR, HER2, HER3, MET, PTK7, ROR1 | CALU1 | Lung | Lung (NSCLC) |
| AXL, EGFR, HER2, HER3, MET, PTK7, ROR1 | CAPAN2 | Pancreas | Pancreas |
| AXL, EGFR, HER2, HER3, MET, PTK7, ROR1 | EBC1 | Lung | Lung (NSCLC) |
| HER2, HER3 | HCC1954 | Breast | Breast (Her2+) |
| AXL, IGF1R | HCC4006 | Lung | Lung (NSCLC) |
| EGFR, EPHA2, HER2, HER3, IGF1R, MET, PTK7, ROR1 | HCC827 | Lung | Lung (NSCLC) |
| AXL, EGFR, MET | HCT116 | Bowel | Bowel (MSI) |
| AXL, EGFR, HER3, MET, PTK7, ROR1 | HS578T | Breast | Breast (TNBC) |
| EGFR, HER2, HER3, MET, PTK7, ROR1 | HT29 | Bowel | Bowel (MSS) |
| HER3 | KatoIII | Gastric | Gastric |
| AXL | LS123 | Bowel | Bowel (MSS) |
| EGFR, HER3, MET, PTK7, ROR1 | MDAMB468 | Breast | Breast (TNBC) |
| AXL | MKN 7 | Gastric | Gastric |
| AXL | NCIH1563 | Lung | Lung (NSCLC) |
| AXL, DDR1, EGFR, EPHA2, HER2, HER3, FGFR2, IGF1R, MET, PTK7, ROR1 | NCIH1650 | Lung | Lung (NSCLC) |
| AXL, EGFR, EPHA2, HER2, HER3, MET, PDGFRA | NCIH1975 | Lung | Lung (NSCLC) |
| EGFR, EPHA2, HER2, KDR, MET | NCIH358 | Lung | Lung (NSCLC) |
| MET, PTK7 | NCIH441 | Lung | Lung (NSCLC) |
| EGFR, HER2, HER3, MET, PTK7, ROR1 | NCIN87 | Gastric | Gastric |
| EGFR, HER2, HER3, | SKBR3 | Breast | Breast (Her2+) |
| EGFR, MET | SKMES1 | Lung | Lung (NSCLC) |
| EGFR, MET | SW48 | Bowel | Bowel (MSI) |
| EGFR, MET | SW900 | Lung | Lung (NSCLC) |

**b**

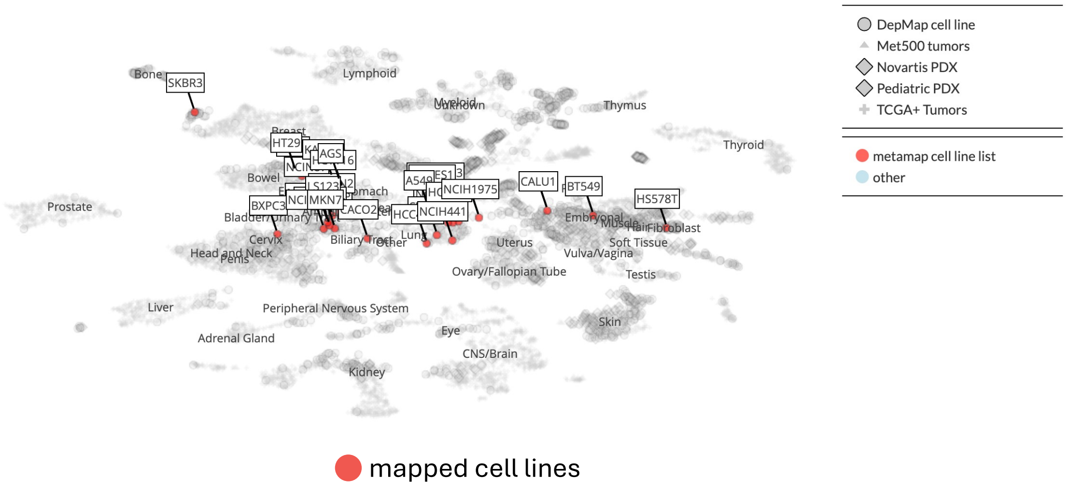

**Supplementary Figure 3. Cell systems used for RTK micromapping. a)** Cell systems used for mapping and the corresponding receptor tyrosine kinase (RTK) targets profiled in each system. Cell models were selected to capture a diverse set of tumor lineages and microenvironmental contexts. **b)** DepMap Celligner representation of cell lines used for mapping, showing their transcriptional similarity to tumor tissues based on RNA-seq data. Cell lines are positioned according to inferred relationships to primary tumor types, with red points indicating those used in this study. Cell line selection considered the purported tissue of origin and its alignment with the target tissue.

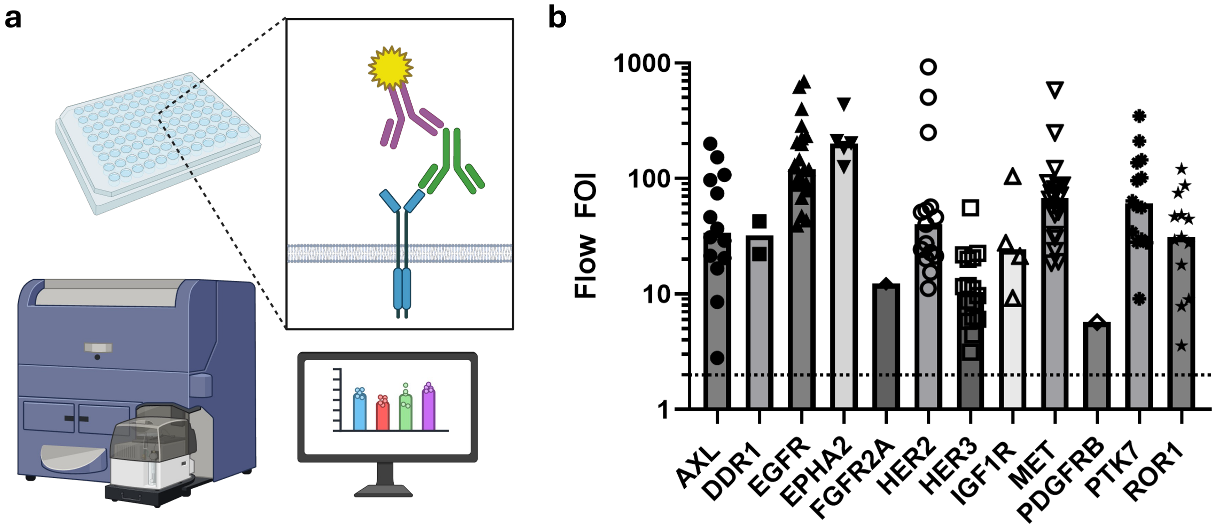

**Supplementary Figure 4. Flow cytometry analysis for cell surface expression of micromapped RTK targets. a)** Schematic of flow cytometry profiling of RTKs. Created in BioRender. Created in BioRender. May, C. (2026) https://BioRender.com/5clk4vw. **b)** Fold-over-isotype signal for the cell line–target pairs used in the large-scale micromapping study, categorized by target. Each symbol represents an individual binder–cell line combination; bars indicate the geometric mean fold change. The dashed line indicates a threshold fold-over-isotype value of 2. These measurements confirm target-specific binding and expression levels across the profiled cell systems.

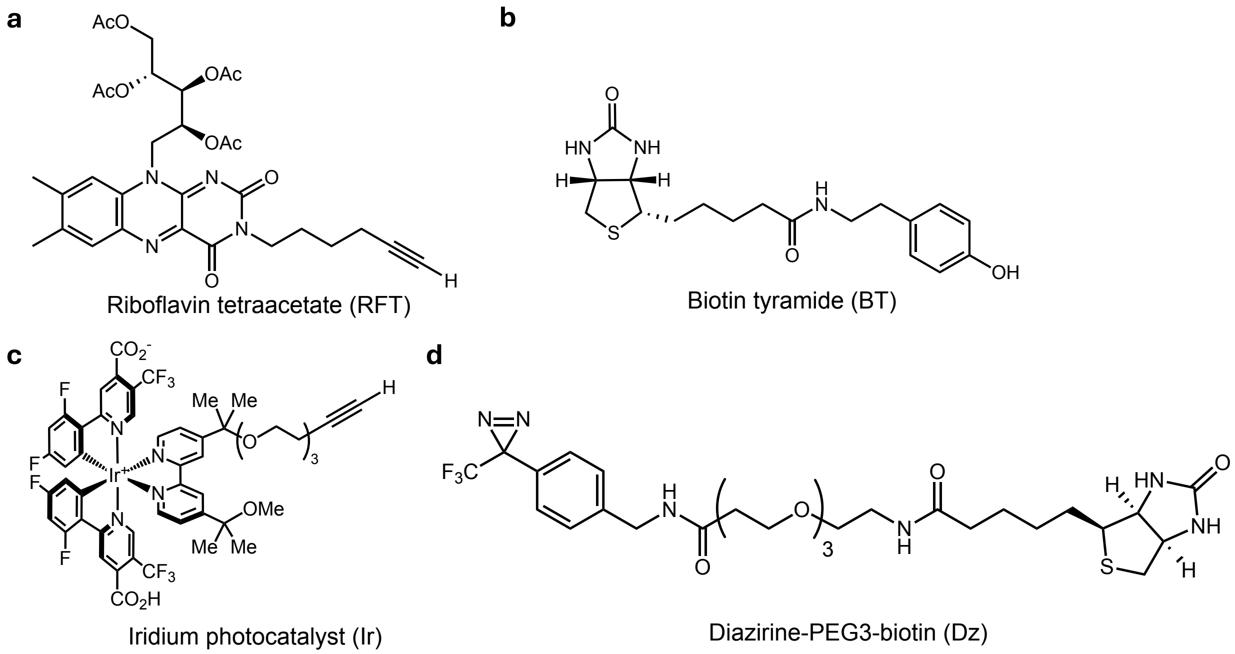

**Supplementary Figure 5. Chemical structures of photocatalysts and small-molecule probes. a)** Riboflavin tetraacetate (RFT) photocatalyst and **b)** Biotin tyramide (BT), a biotin-containing phenol probe used for RFT-mediated proximity labeling**. c)** Iridium photocatalyst (Ir) and **d)** Diazirine-PEG$_3$-Biotin (Dz), a biotin-containing aryl diazirine probe used for IR-mediated proximity labeling.

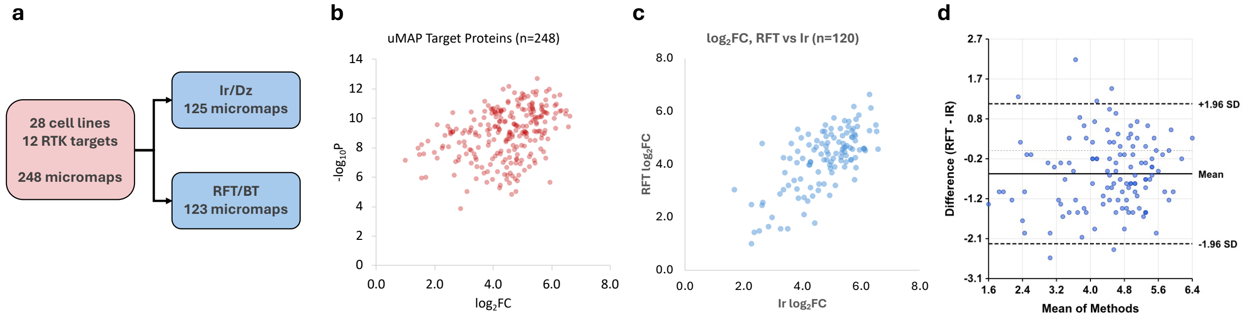

**Supplementary Figure 6. Target protein enrichment across all RTK micromaps. a)** Schematic of total microenvironment mapping experiments and workflow breakdown by Ir/Dz and RFT/BT photocatalyst probe pair labeling approaches. **b)** Volcano plot showing enrichment of targets across 248 micromap experiments. Each point represents the target from an individual micromap experiment, plotted as $\log_2$ fold change over an isotype matched non-specific IgG antibody (x-axis) versus the $-\log_{10}$ p-value (y-axis). Robust enrichment of the intended target was observed in all experiments, highlighting successful targeting and consistent enrichment across receptor tyrosine kinase (RTK) micromaps. **c)** The $\log_2$ fold change values shown in b are plotted pair-wise for the 120 matched target and cell line experiments that were performed with both photochemical systems (three Ir targets did not have a cell-line matched micromap for the RFT photochemistry and one RFT target lacked a corresponding Ir micromap). Matched Ir and RFT experiments showed only a modest quantitative correlation (Pearson r = 0.691, $R^2$ = 0.477, n = 120). **d**) However, Bland-Altman analysis revealed excellent agreement: mean bias = -0.563 (95% CI of differences: -2.26 to 1.13), with 5.0% of points outside the limits of agreement.

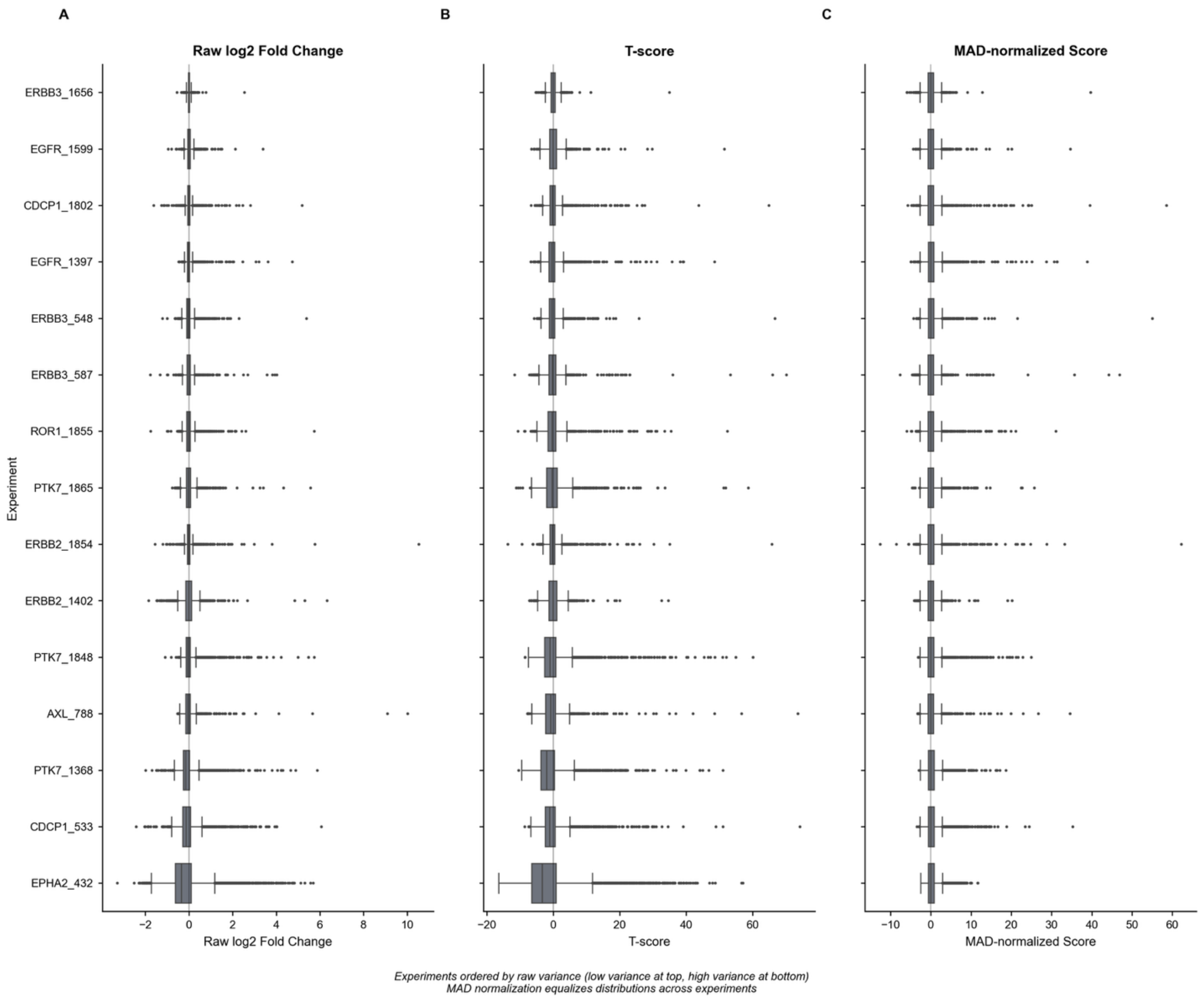

**Supplementary Figure 7. Micromap cross-experiment normalization**. Box plots showing the distribution of protein enrichment scores for 15 representative micromapping experiments at three stages of the normalization pipeline: (A) raw log2 fold change values, (B) T-scores from replicate comparisons, and (C) MAD-normalized scores. Experiments are ordered vertically by increasing variance in raw log2 fold change values (lowest variance at top, highest at bottom). While raw fold change distributions (A) and T-scores (B) show substantial heterogeneity in spread—reflecting intrinsic differences between "quiet" protein targets with few strong interactors and "loud" targets with many enriched proteins—MAD normalization (C) produces comparable distributions across all experiments. This transformation enables direct comparison of interaction confidence scores across the dataset regardless of individual experiment characteristics, facilitating integrated analysis and threshold-based hit calling.

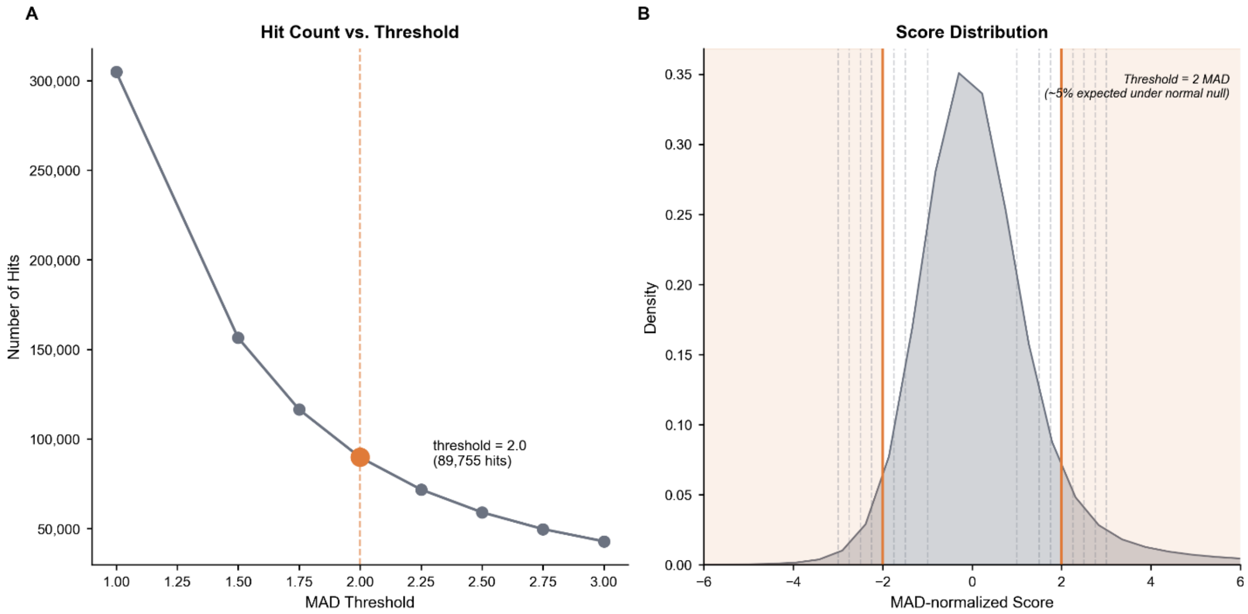

**Supplementary Figure 8. Threshold sensitivity analysis for MAD-normalized hit calling.** (A) Number of interactions exceeding the MAD-normalized score threshold as a function of threshold stringency. The relationship follows an inverse curve typical of score-based filtering, with 89,755 bait-prey pairs meeting the threshold of 2.0 MAD (orange). (B) Kernel density estimate of the MAD-normalized score distribution across all target protein – enriched protein pairs. Vertical lines indicate candidate thresholds ranging from 1.0 to 3.0 MAD, with the selected threshold of 2.0 MAD highlighted (orange solid lines). Shaded regions denote scores exceeding ±2.0 MAD, corresponding to approximately 5% of observations expected by chance under a normal null distribution. This threshold balances sensitivity for detecting true interactions against specificity for excluding background noise, analogous to a two-tailed α = 0.05 significance criterion.

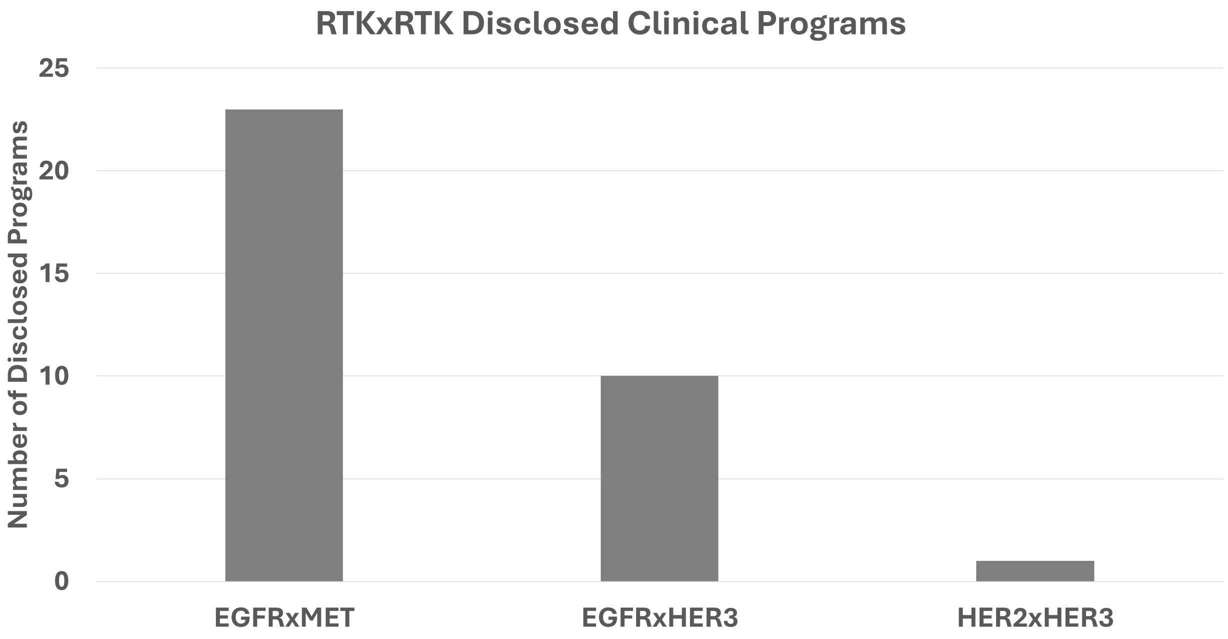

**Supplementary Figure 9. Limited clinical exploration of heterotypic RTK–RTK bispecifics.** Bar plot showing the number of publicly disclosed clinical-stage bispecific antibodies, antibody–drug conjugates (ADCs), or T-cell engagers targeting heterotypic RTK–RTK pairs. Counts were curated using Synapse by Patsnap and exclude discovery and preclinical programs. Clinical activity was limited to three RTK–RTK pairs (EGFR×MET, EGFR×HER3, and HER2×HER3) indicating that most RTK–RTK interactions identified by proximity mapping remain clinically unexplored.

| RTK_a | RTK_b | jaccard |
|---|---|---|
| AXL | DDR1 | 0.176 |
| AXL | EGFR | 0.324 |
| AXL | EPHA2 | 0.292 |
| AXL | ERBB2 | 0.260 |
| AXL | ERBB3 | 0.298 |
| AXL | FGFR2 | 0.004 |
| AXL | IGF1R | 0.269 |
| AXL | MET | 0.300 |
| AXL | PDGFRB | 0.011 |
| AXL | PTK7 | 0.307 |
| AXL | ROR1 | 0.269 |
| DDR1 | EGFR | 0.145 |
| DDR1 | EPHA2 | 0.146 |
| DDR1 | ERBB2 | 0.100 |
| DDR1 | ERBB3 | 0.178 |
| DDR1 | FGFR2 | 0.012 |
| DDR1 | IGF1R | 0.349 |
| DDR1 | MET | 0.111 |
| DDR1 | PDGFRB | 0.012 |
| DDR1 | PTK7 | 0.129 |
| DDR1 | ROR1 | 0.320 |
| EGFR | EPHA2 | 0.427 |
| EGFR | ERBB2 | 0.369 |
| EGFR | ERBB3 | 0.282 |
| EGFR | FGFR2 | 0.005 |
| EGFR | IGF1R | 0.235 |
| EGFR | MET | 0.490 |
| EGFR | PDGFRB | 0.005 |
| EGFR | PTK7 | 0.471 |
| EGFR | ROR1 | 0.265 |
| EPHA2 | ERBB2 | 0.392 |
| EPHA2 | ERBB3 | 0.270 |
| EPHA2 | FGFR2 | 0.003 |
| EPHA2 | IGF1R | 0.246 |
| EPHA2 | MET | 0.436 |
| EPHA2 | PDGFRB | 0.003 |
| EPHA2 | PTK7 | 0.440 |
| EPHA2 | ROR1 | 0.272 |
| ERBB2 | ERBB3 | 0.256 |
| ERBB2 | FGFR2 | 0.004 |
| ERBB2 | IGF1R | 0.168 |
| ERBB2 | MET | 0.406 |
| ERBB2 | PDGFRB | 0.004 |
| ERBB2 | PTK7 | 0.444 |
| ERBB2 | ROR1 | 0.233 |
| ERBB3 | FGFR2 | 0.006 |
| ERBB3 | IGF1R | 0.252 |
| ERBB3 | MET | 0.329 |
| ERBB3 | PDGFRB | 0.006 |
| ERBB3 | PTK7 | 0.303 |
| ERBB3 | ROR1 | 0.272 |
| FGFR2 | IGF1R | 0.006 |
| FGFR2 | MET | 0.003 |
| FGFR2 | PDGFRB | 0.083 |
| FGFR2 | PTK7 | 0.004 |
| FGFR2 | ROR1 | 0.006 |
| IGF1R | MET | 0.197 |
| IGF1R | PDGFRB | 0.010 |
| IGF1R | PTK7 | 0.215 |
| IGF1R | ROR1 | 0.352 |
| MET | PDGFRB | 0.004 |
| MET | PTK7 | 0.504 |
| MET | ROR1 | 0.220 |
| PDGFRB | PTK7 | 0.003 |
| PDGFRB | ROR1 | 0.014 |
| PTK7 | ROR1 | 0.251 |

**Supplementary Figure 10. Jaccard similarity indices for protein target pairs**. This table lists the Jaccard values calculated for every pair of target proteins. The Jaccard index quantifies the overlap in the enriched protein microenvironment between each target pair. These similarity values were used to determine the hierarchical ordering and arrangement of the protein targets in the Venn diagram shown in Figure 2d.

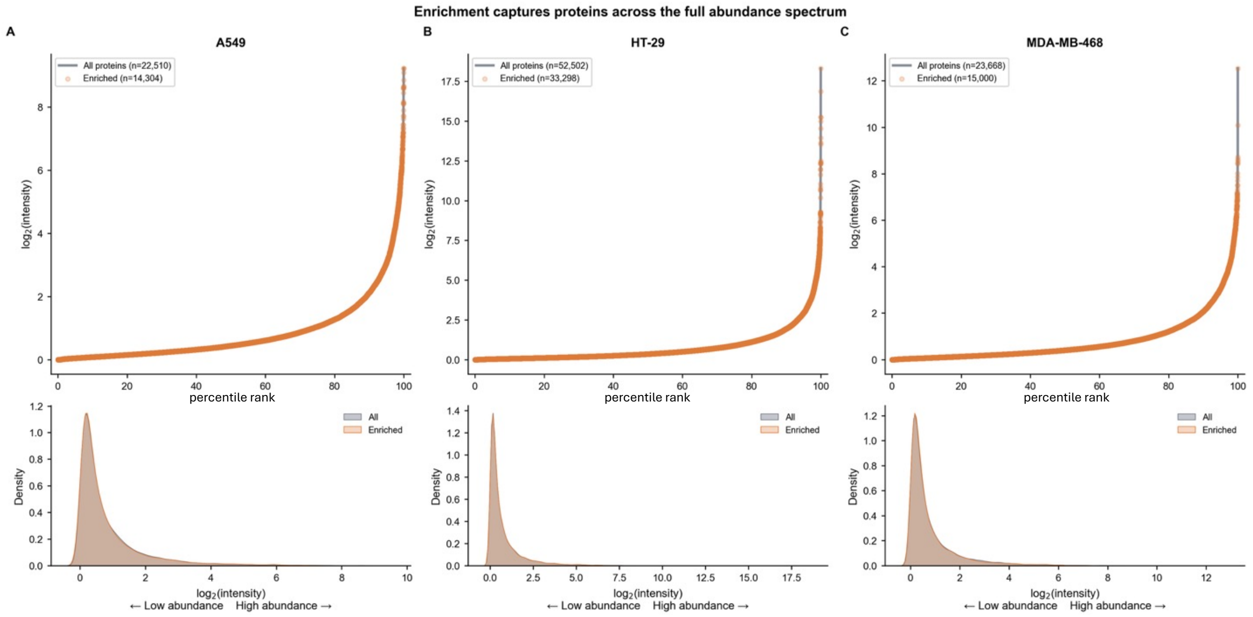

**Supplementary Figure 11. Enrichment captures proteins across the full proteomic abundance spectrum.** Comparison of protein abundance distributions between all detected proteins and micromap-enriched proteins across three representative cell lines (A549, lung adenocarcinoma; HT-29, colorectal adenocarcinoma; MDA-MB-468, triple-negative breast cancer). Top panels (A-C): Ranked abundance plots showing all proteins as a gray curve with micromap enriched proteins overlaid as orange points. Proteins are ranked by DIA-measured normalized intensity (x-axis: 0th to 100th percentile). Enriched proteins are distributed across the entire abundance range, from low-abundance (left) to high-abundance (right) proteins. Bottom panels: Kernel density estimation (KDE) plots of $log_2$-transformed normalized intensity for all proteins (gray) versus enriched proteins (orange). The near-complete overlap of distributions demonstrates that enrichment is not biased toward highly abundant proteins. Kolmogorov-Smirnov tests confirmed negligible effect sizes across all cell lines ($D < 0.02$), indicating that micromap enrichment samples proteins representatively across the full dynamic range of cellular protein abundance.

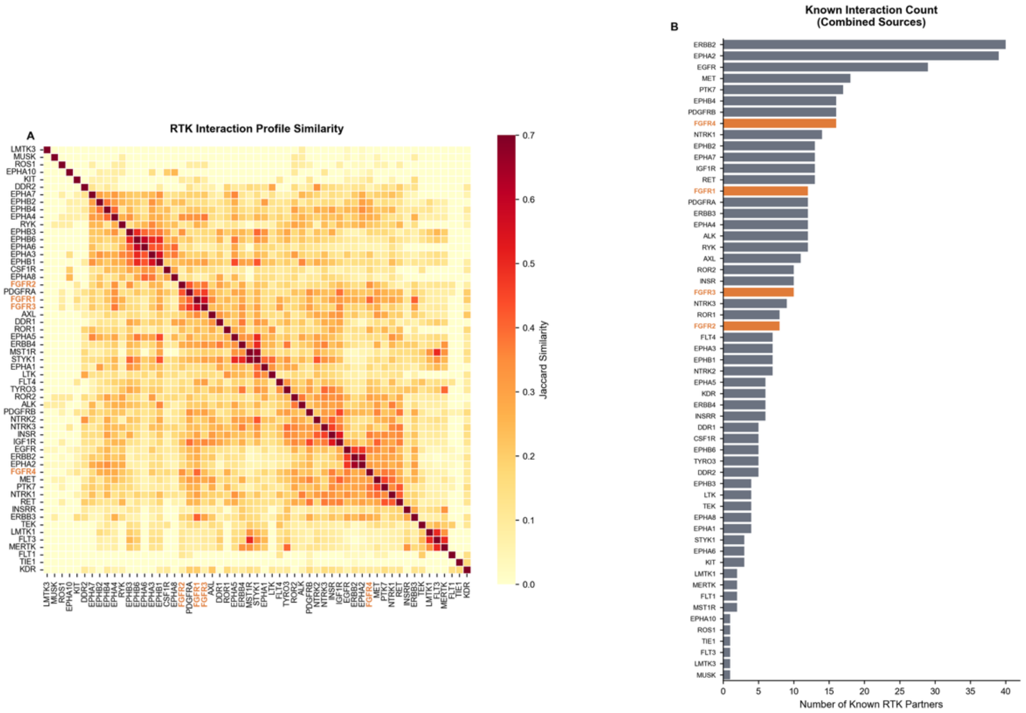

**Supplementary Figure 12. FGFR2 has fewer documented RTK-RTK interactions compared to other receptor tyrosine kinases. A)** Jaccard similarity heatmap comparing known RTK-RTK interaction profiles across 57 receptor tyrosine kinases. Interactions were compiled from curated literature, BioGRID, STRING (combined score ≥700), and IntAct databases. Hierarchical clustering (average linkage) groups RTKs with similar interaction partners. FGFR family members (FGFR1–4) are highlighted in orange. Higher Jaccard similarity (darker red) indicates greater overlap in known interaction partners between RTK pairs. **B)** Number of known RTK-RTK interaction partners per receptor tyrosine kinase from combined reference sources. FGFR2 has 8 documented partners (EPHA2, EPHA4, ERBB2, ERBB3, FGFR1, FGFR3, FLT4, RYK), the fewest among FGFR family members (FGFR1: 12, FGFR3: 10, FGFR4: 16) and substantially fewer than well-characterized RTKs such as ERBB2 (40), EPHA2 (36), and EGFR (29). This sparse annotation in reference databases, combined with limited experimental coverage (single cell line), may contribute to FGFR2's distinct microenvironment profile observed in MetaMap analyses.

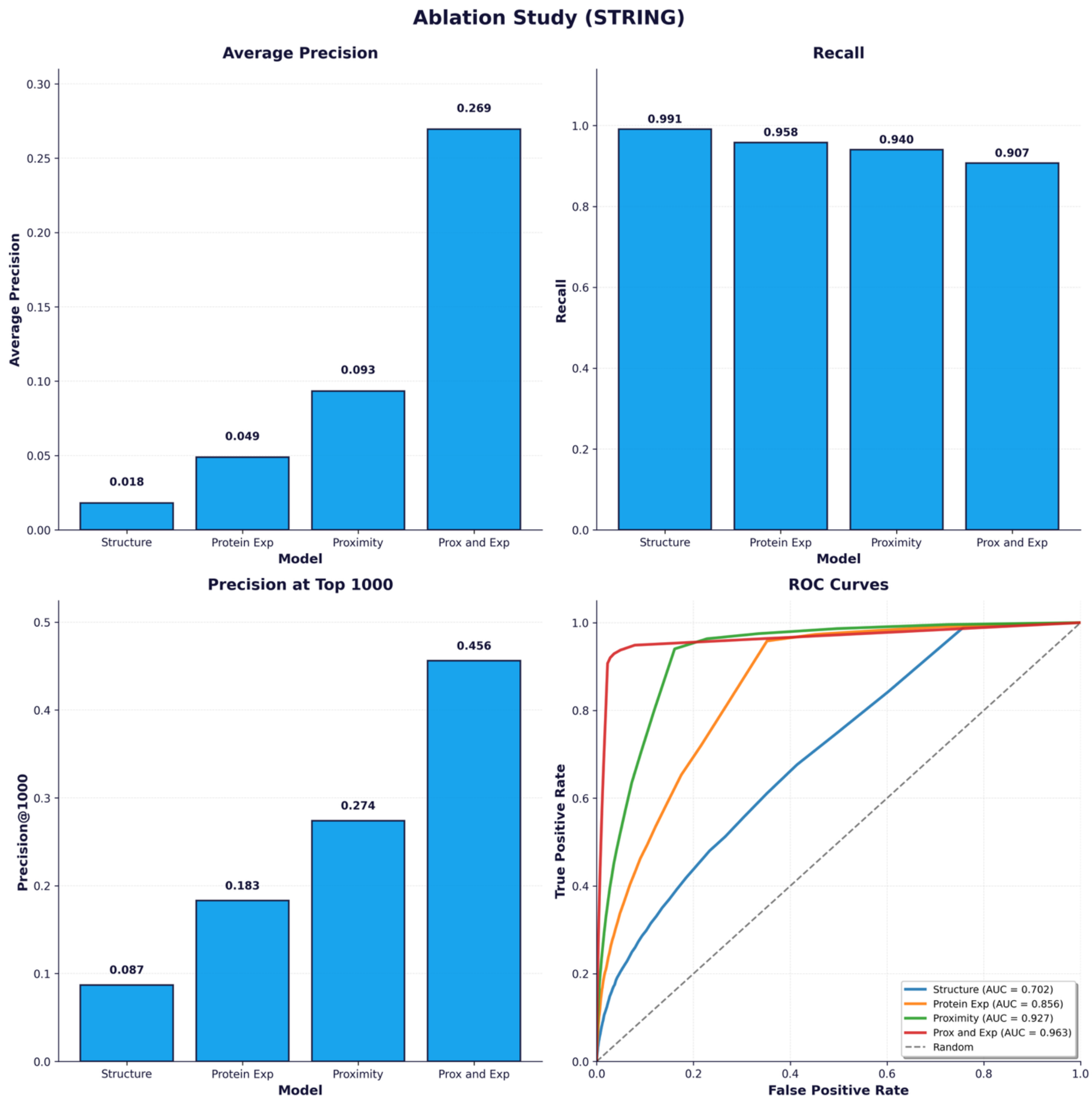

**Supplementary Figure 13. Ablation study of GAT model performance across key evaluation metrics**. Feature ablation analysis as in Figure 4c, showing GAT model average precision, recall, precision@1000, and AUC ROC across training conditions: graph structure alone (Structure), graph structure with protein expression features (Protein exp), graph structure with proximity features (Proximity), or full feature set (Prox and exp). Upper left panel reproduced from Figure 4c.

**a**

| Cell line | Lineage | Subtype info |
|---|---|---|
| A549 | Lung | Lung (NSCLC) |
| AGS | Gastric | Gastric |
| BT549 | Breast | Breast (TNBC) |
| BXPC3 | Pancreas | Pancreas |
| CACO2 | Bowel | Bowel (MSS) |
| CALU1 | Lung | Lung (NSCLC) |
| CAPAN2 | Pancreas | Pancreas |
| EBC1 | Lung | Lung (NSCLC) |
| HCC827 | Lung | Lung (NSCLC) |
| HPAC | Pancreas | Pancreas |
| HT29 | Bowel | Bowel (MSS) |
| MDAMB468 | Breast | Breast (TNBC) |
| NCIH1650 | Lung | Lung (NSCLC) |
| NCIH1975 | Lung | Lung (NSCLC) |
| NCIH358 | Lung | Lung (NSCLC) |
| NCIN87 | Gastric | Gastric |
| SKMES1 | Lung | Lung (NSCLC) |
| SW48 | Bowel | Bowel (MSI) |
| SW900 | Lung | Lung (NSCLC) |

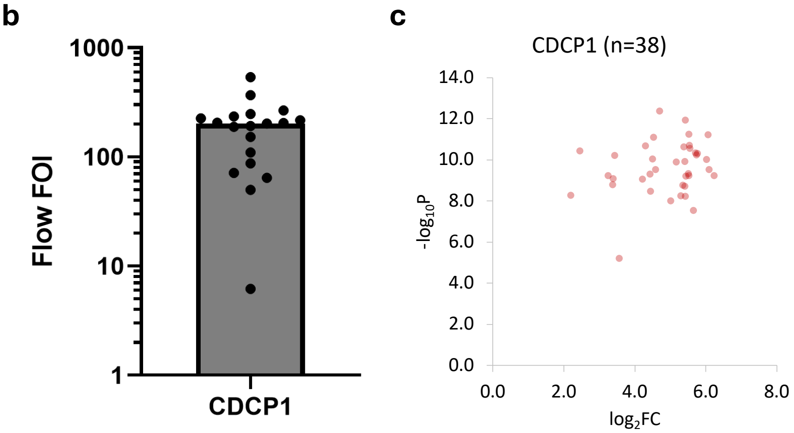

**Supplementary Figure 14. Flow cytometry and micromap enrichment analysis of CDCP1. a)** Table of the 19 cell systems used for mapping CDCP1 with Ir and RFT chemistries. **b)** Fold-over-isotype signal for the cell line–target pairs used in CDCP1 micromapping. Each symbol represents an individual binder–cell line combination; bars indicate the geometric mean fold change. The dashed line indicates a threshold fold-over-isotype value of 2. These measurements confirm target-specific binding and expression levels across the profiled cell systems. **c)** Plot of $-log_{10}$ p-value ($-log_{10}P$) vs $log_2$ fold change ($log_2FC$) of CDCP1 protein enrichment over isotype matched non-specific IgG antibody across 38 micromap experiments (each point represents an individual CDCP1-targeted micromap). Robust enrichment of CDCP1 was observed across all experiments, confirming successful targeting and consistent recovery of the target protein across all CDCP1-directed micromaps.

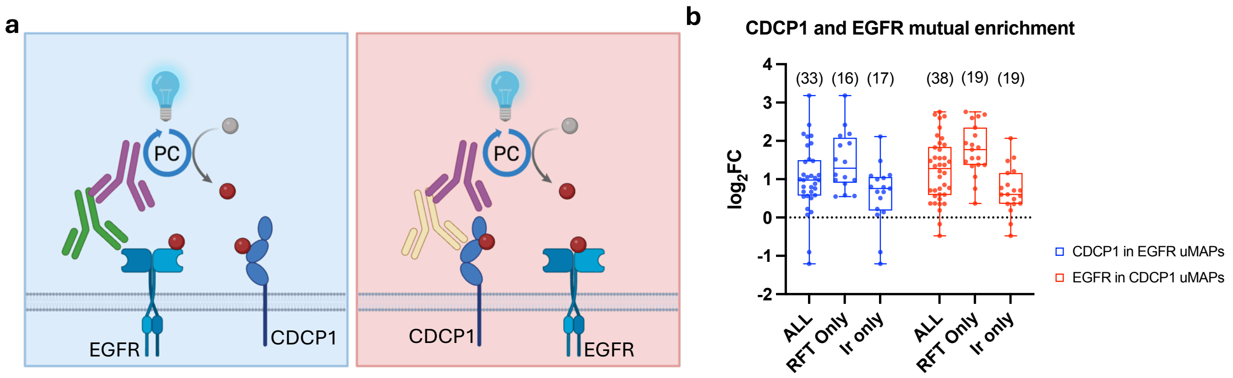

**Supplementary Figure 15. EGFR and CDCP1 enrich each other through targeted micromapping. a)** Schematic showing targeted photocatalytic labeling of either CDCP1 or EGFR and the resulting capture of proximal EGFR and CDCP1. Created in BioRender. May, C. (2026) https://BioRender.com/hs5gs92. **b)** Box plots summarize the distribution of enrichment values ($\log_2$ fold-change of targeted protein over isotype matched non-specific IgG antibody) for CDCP1 from EGFR-targeted microenvironment mapping and EGFR from CDCP1-targeted microenvironment mapping. Each point represents an individual microenvironment mapping measurement for either EGFR (blue) or CDCP1 (red) across RFT only mapping, Ir only mapping, or combined (ALL). Boxes indicate the interquartile range, the horizontal line denotes the median, and whiskers represent 1.5× the interquartile range.

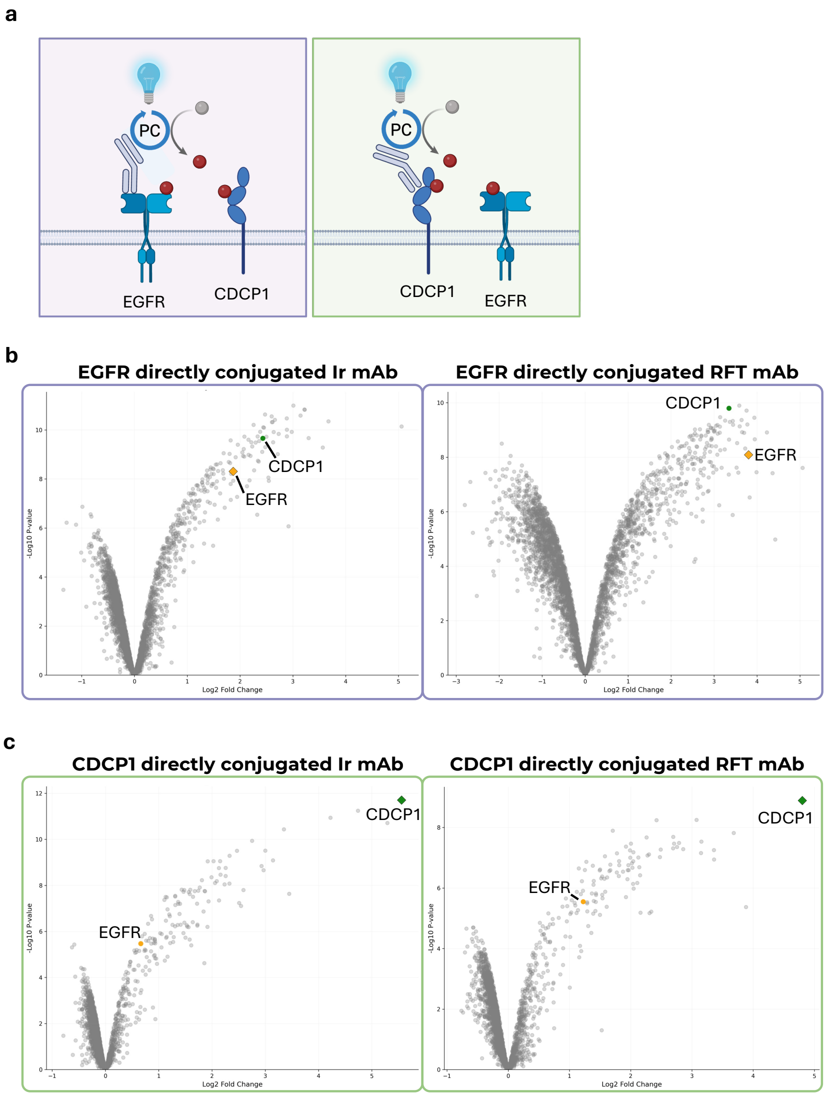

**Supplementary Figure 16. CDCP1 and EGFR targeted micromapping using directly conjugated antibody-photocatalyst conjugates. a)** Schematic showing targeted photocatalytic labeling of either CDCP1 or EGFR using a directly conjugated monovalent antibody photocatalyst and the resulting capture of proximal EGFR and CDCP1. Created in BioRender. May, C. (2026) https://BioRender.com/smg7oq0. **b)** Volcano plots from two independent micromap experiments on HCC827 lung adenocarcinoma cells. In both datasets, EGFR (orange diamond) and CDCP1 (green dot) show significant enrichment together (p-value < 0.05 and $log_2FC > 1.5$) (n = 3 experiments). Notably, the antibody in this experiment (cetuximab) binds a different epitope compared to the antibody used in Figure 2a and Supplementary Figures 4 and 6. **c)** Volcano plots from two independent micromap experiments on NCIH1650 lung adenocarcinoma cells using a directly conjugated antibody photocatalyst. In both datasets, CDCP1 (green diamond) and EGFR (orange circle) show significant co-enrichment (p-value < 0.05 and $log_2FC > 1.5$) (n = 3 experiments). The antibody in this experiment (CDCP1 41A9) binds a different epitope compared to the antibody used in Supplementary Figure 14. These results demonstrate that the observed EGFR-CDCP1 proximity is independent of binder epitope and direct antibody–photocatalyst conjugation.

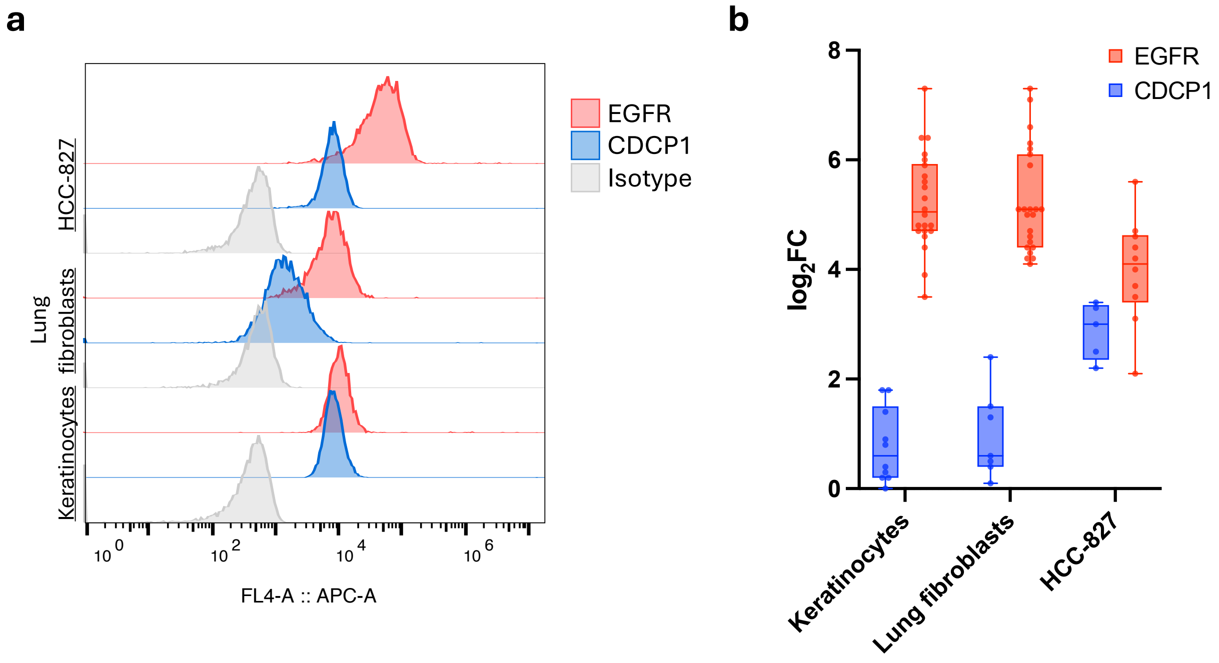

**Supplementary Figure 17. EGFR micromapping reveals tumor-selective CDCP1 enrichment. a)** Flow cytometry data comparison of EGFR and CDCP1 surface expression in two normal cell lines (keratinocytes and normal lung fibroblasts) and HCC-827 cancer cells. **b)** EGFR targeted micromaps recovered higher levels of CDCP1 from HCC-827 tumor cells than from the two normal cell populations. For each cell model, individual $\log_2$ peptide ratios of EGFR-targeted vs isotype-control samples are plotted for peptides corresponding to EGFR (red) and CDCP1 (blue). The box midpoints represent the sample medians, box boundaries indicate the first and third quartiles, and whiskers extend from the minimum to the maximum values.

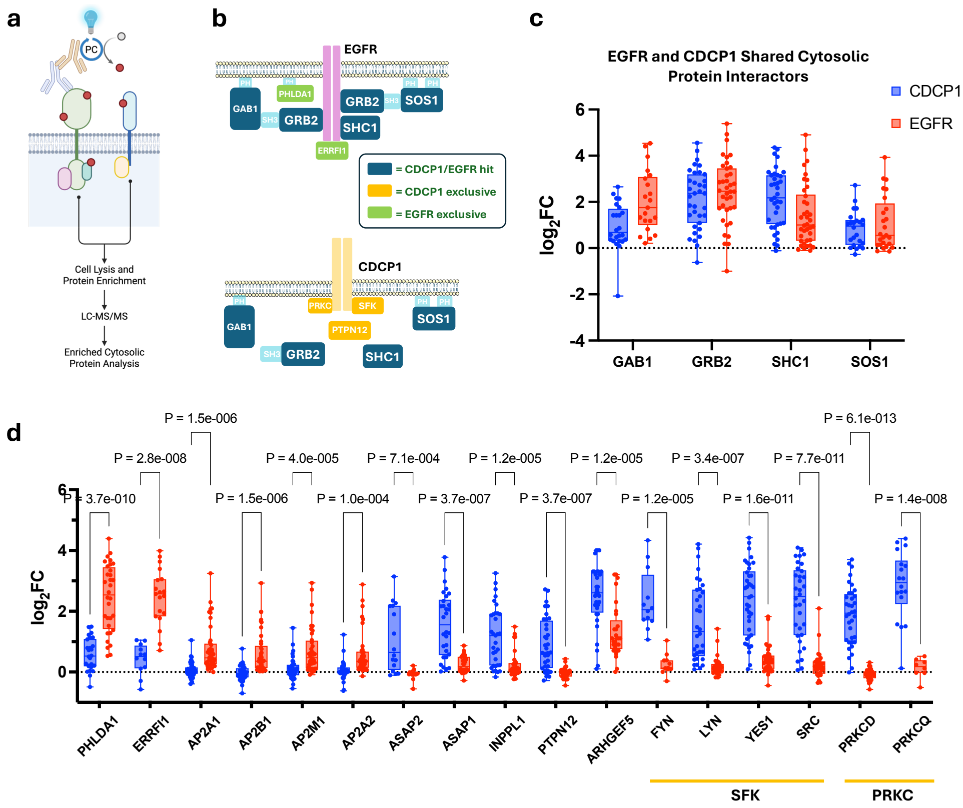

**Supplementary Figure 18. Micromapping of EGFR and CDCP1 reveals convergent and divergent cytosolic protein interactors. a)** Schematic depicting the workflow to enrich and quantify cytosolic proteins in proximity to EGFR and CDCP1. Created in BioRender. May, C. (2026) https://BioRender.com/czujvdc **b)** Schematic of EGFR and CDCP1 scaffolding and adaptor proteins[1-3] identified via microenvironment mapping. Proteins in dark blue indicate shared enrichment by both EGFR and CDCP1, green indicates EGFR-preferential interactors, and yellow indicates CDCP1-preferential interactors. **c)** Box plots show that both EGFR (red) and CDCP1 (blue) enrich signaling proteins GAB1, GRB2, SHC1, and SOS1. **d)** Box plots showing proteins preferentially enriched by EGFR (red) or CDCP1 (blue). CDCP1 has strong preference for SRC family kinases (SFKs), Protein Kinase C delta and theta proteins (PRKCs), PTPN12, INPPL1, and ASAP1/2. EGFR interacts with PH domain containing protein PHLDA1 and the negative feedback inhibitor protein ERRFI1. EGFR also shows a more modest enrichment of components of the AP-2 adaptor complex. Box plots summarize the distribution of enrichment values ($log_2$ fold-change of targeted protein over isotype matched non-specific IgG antibody) for each cytosolic interactor across replicate micromap experiments. Each point represents an individual micromap measurement for either EGFR (red) or CDCP1 (blue). Boxes indicate the interquartile range, horizontal lines denote the median, and whiskers indicate the full range of values. Note that enriched cytosolic proteins may result from either receptor-mediated proximity labeling and/or through direct recruitment as physically associated complexes. P-values were calculated using Welch's t-test with unequal variance. Multiple hypothesis testing was done using the False Discovery Rate (FDR) with the two-stage step-up method developed by Benjamini, Krieger, and Yekutieli[4], and a desired FDR (Q) set at 1%. The number of measurements for each protein with each target ranged from n = 8 to n = 41, with a median = 32.

**a** CDCP1 Plasma Membrane Abundance: Normal vs Malignant Tissue (Non Small Cell Lung Cancer)

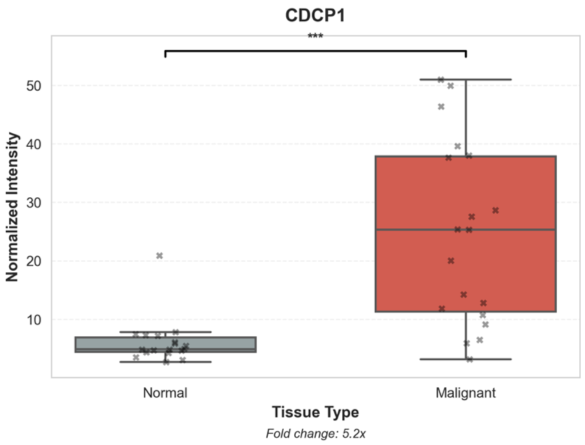

**b** CDCP1 Plasma Membrane Abundance: Normal vs Malignant Tissue (Colon Cancer)

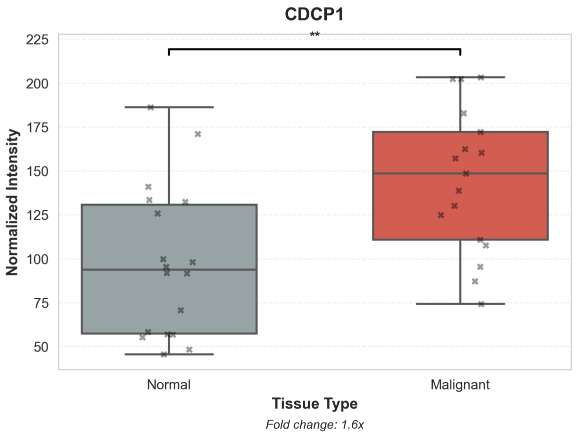

**Supplementary Figure 19. In house proteomic analysis of NSCLC and CRC tumor vs matched normal tissues for CDCP1 protein level quantification**. Box-and-whisker plots show normalized plasma membrane abundance of CDCP1 measured across normal and malignant human tissues. **a)** non-small cell lung cancer (NSCLC), where CDCP1 exhibits a marked increase in malignant tissue samples relative to matched normal tissue (median fold change = 5.2×). **b)** Colon cancer (CRC), where CDCP1 is also significantly elevated in malignant tissue samples (median fold change = 1.6×). Individual data points represent independent tissue samples. Statistical significance was assessed using a two-sided test; *P*-values are indicated above comparisons (**, $P < 0.01$; ***, $P < 0.001$). These data match the observation of increased CDCP1 protein levels in tumor vs matched normal tissues in Figure 4.

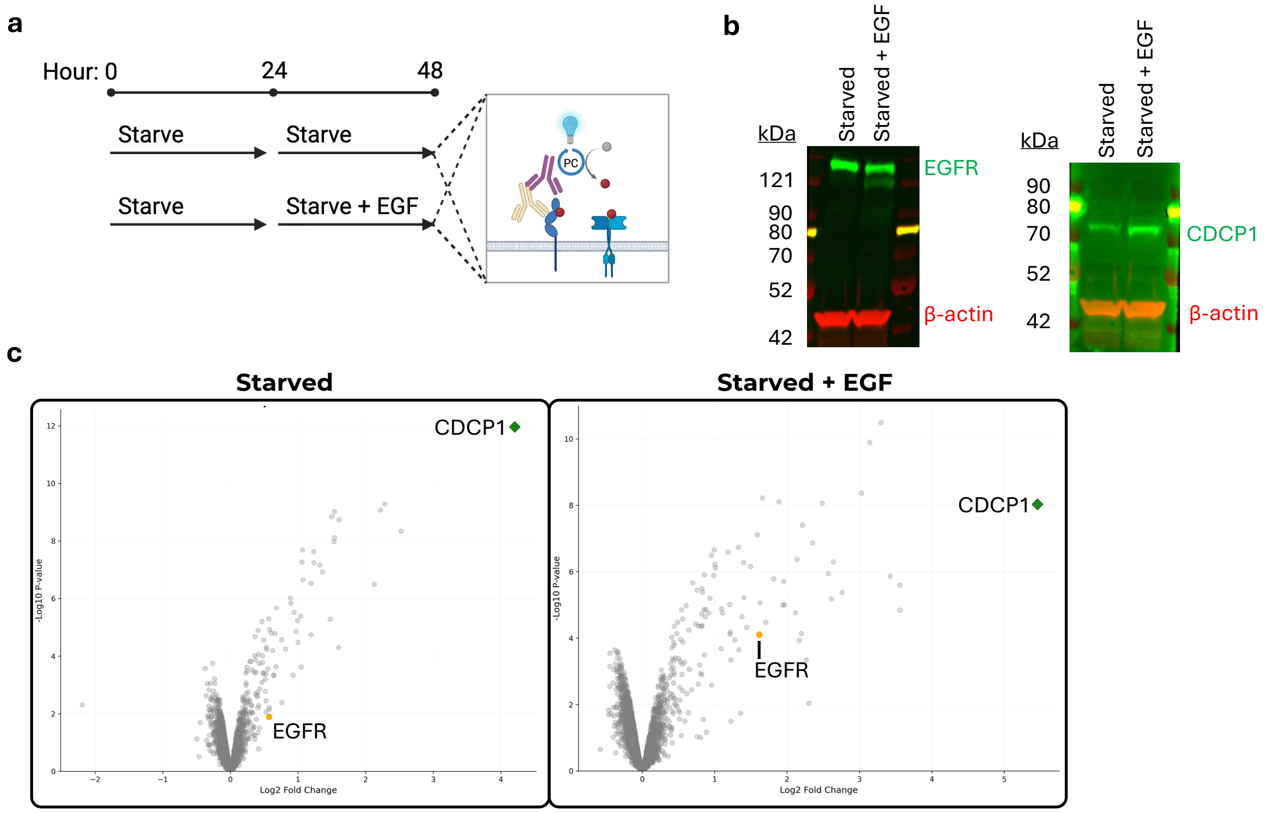

**Supplementary Figure 20. Micromapping of CDCP1 in EGF treated CAOV3 cells. a)** Schematic showing CAOV3 starvation and EGF stimulation. Cells were starved for 24 hours. Starvation media was replaced for an additional 24 hours with one set receiving EGF. Created in BioRender. May, C. (2026) https://BioRender.com/564j6jf. **b)** Western blots of EGFR and CDCP1 show upregulation of CDCP1 upon EGF stimulation. β-actin was included as a loading control. **c)** Volcano plots of CDCP1-targeted micromaps with Iridium/Diazirine photocatalyst probe pair detect EGFR in both conditions with increased EGFR enrichment observed upon EGF stimulation. Volcano plots are plotted as -$\log_{10}$ p-value vs $\log_2$FC with CDCP1 highlighted as a green diamond and EGFR highlighted as an orange dot (n = 3 experiments).

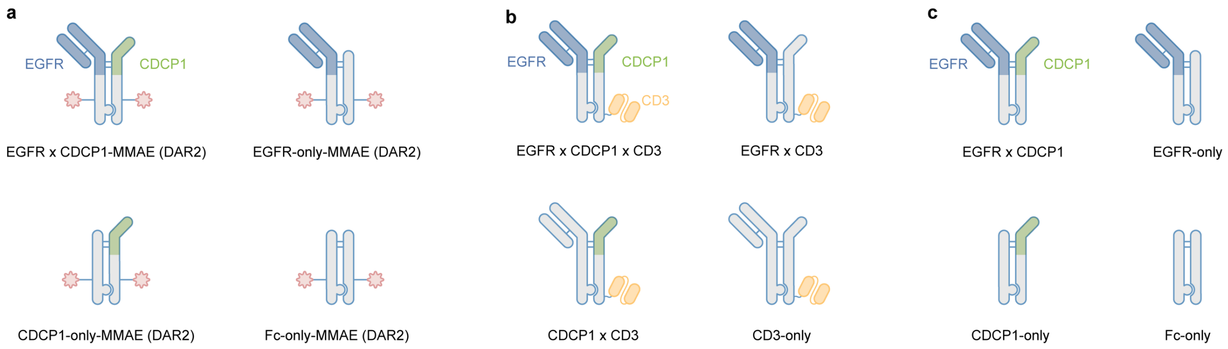

**Supplementary Figure 21. EGFR x CDCP1 targeting modalities. a)** Diagrams of site-specific ADC molecules. Top left: EGFR Fab x CDCP1 VHH bispecific. Top right: EGFR-only ADC with matched DAR. Bottom left: CDCP1-only control with matched DAR. Bottom right: non-targeting Fc-only. A version of these molecules without engineered cysteines where the linker-payload was instead conjugated to interchain disulfides with a DAR ~3 was used in some experiments as indicated. **b)** Diagrams of TCE molecules. Top left: a trispecific EGFR x CDCP1 x CD3 molecule. Top right: EGFR x CD3 control, Bottom left: CDCP1 x CD3 control, Bottom right: non-targeting CD3-only control. **c)** Diagrams of unconjugated bispecific molecules used in internalization assays. Top left: EGFR Fab x CDCP1 VHH bispecific. Top right: EGFR-only. Bottom left: CDCP1-only control. Bottom right: non-targeting Fc-only.

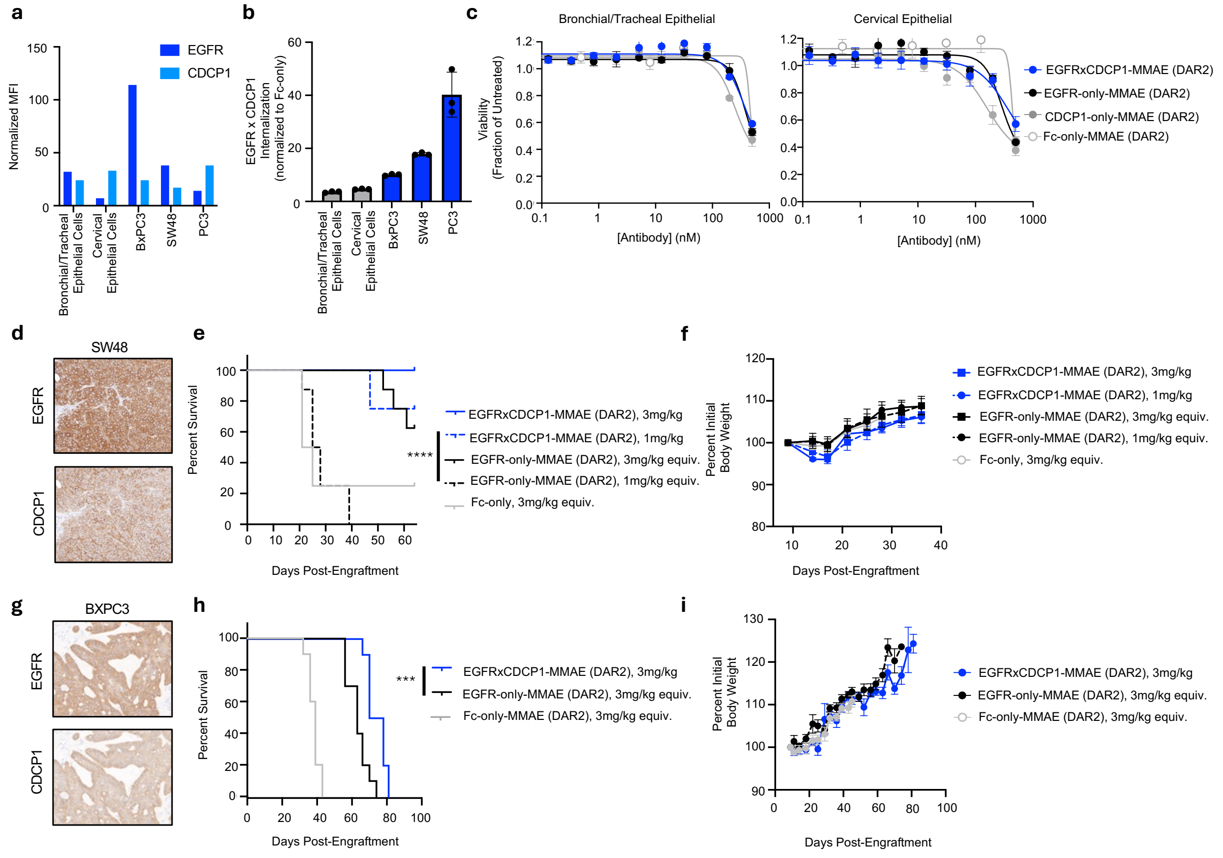

**Supplementary Figure 22. EGFR x CDCP1 bispecific has enhanced activity in cancer cells and shows activity in vivo. a)** EGFR and CDCP1 expression in indicated primary cell and cancer cell systems as assessed by flow cytometry. **b)** Relative internalization of unconjugated EGFR x CDCP1 bispecific in primary cells (grey) and cancer cell lines (blue) at 24 hours. Individual data points are shown with the box at the mean plus and minus standard deviation. **c)** Relative viability of indicated primary cells after EGFR x CDCP1 ADC or matched DAR control treatment (antibody structures depicted in Supplementary Figure 21). Viability was assessed by CellTiter-Glo and normalized to untreated cells. Data points are mean plus and minus standard deviation. **d)** Representative EGFR and CDCP1 expression by IHC in SW48 xenograft tumors. **e)** Kaplan-Meier survival curves for mice in Figure 5E. Tumors that reached 800 $mm^3$ were considered a survival endpoint. Logrank (Mantel-Cox) test, ****, p<0.0001. **f)** Mean body weight plus and minus standard error of the mean (SEM) for mice depicted in Figure 5E. **g)** Representative EGFR and CDCP1 expression by IHC in BxPC3 xenograft tumors. **h)** Kaplan-Meier survival curves for mice in Figure 5G. Tumors that reached 800 $mm^3$ were considered a survival endpoint. Logrank (Mantel-Cox) test, ***, p<0.001. **i)** Mean body weight plus and minus standard error of the mean plus and minus SEM for mice in Figure 5G.

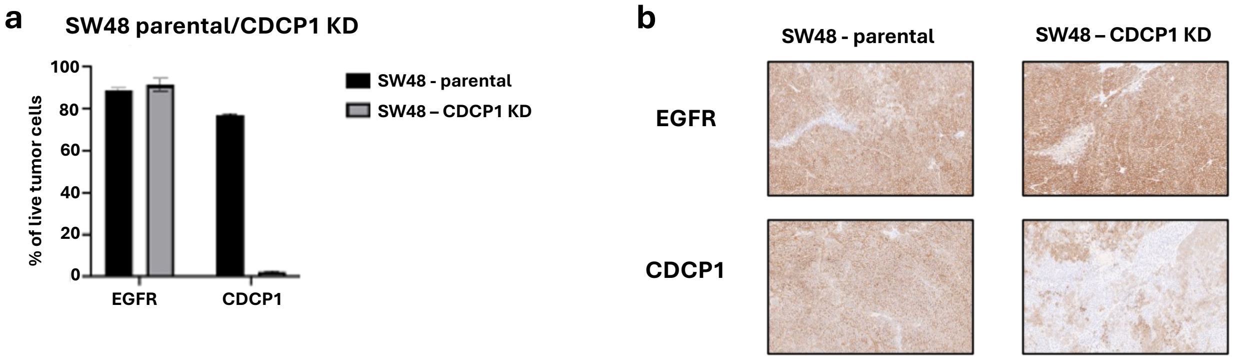

**Supplementary Figure 23. Characterization of SW48 CDCP1-knockdown (KD) tumor xenograft model. a)** EGFR and CDCP1 expression in SW48 parental and CDCP1-KD tumor xenografts by flow cytometry. Tumor cells were implanted subcutaneously in female NCr nude mice and collected once tumor volume reached ~500 mm3 (Day 22). **b)** Representative EGFR and CDCP1 expression by IHC in SW48 parental and CDCP1-KD xenograft tumors.

### Materials and Methods

#### General Cell Culture

The following cancer cell lines were purchased from ATCC: A549 (Cat: CRM-CCL-185), AGS (Cat: CRL-1739), BT549 (Cat: HTB-122), BXPC3 (Cat: CRL-1687), CACO2 (Cat: HTB-37), CAPAN2 (Cat: HTB-80), CAOV3 (Cat: HTB-75), HCC1954 (Cat: CRL-2338), HCC4006 (Cat: CRL-2871), HCC827 (Cat: CRL-2868), HCT116 (Cat: CCL-247), HPAC (Cat: CRL-2119), HS578T (Cat: HTB-126), HT29 (Cat: HTB-38), KatoIII (Cat: HTB-103), LS123 (Cat: CCL-255), MDAMB468 (Cat: HTB-132), NCIH1563 (Cat: CRL-5875), NCIH1650 (Cat: CRL-5883 ), NCIH1975 (Cat: CRL-5908), NCIH358 (Cat: CRL-5807), NCIH441 (Cat: HTB-174), NCIN87 (Cat: CRL-5822), PC3 (Cat: CRL-1435), SKBR3 (Cat: HTB-30), SKMES1 (Cat: HTB-58), SW48 (Cat: CCL-231) and SW900 (Cat: HTB-59). The following cells were purchased from Accegen: CALU1 (Cat: ABC-TC0110), EBC1 (Cat: ABC-TC0170) and MKN 7 (Cat: ABC-TC0688).

The media for all cancer cells was purchased from ATCC and include RPMI 1640 (Cat: 30-2001), DMEM (Cat: 30-2002), EMEM (Cat: 30-2003), F-12K (Cat: 30-2004), IMDM (Cat: 30-2005) and McCoy's 5A (Cat: 30-2007). All media was supplemented with FBS (10% or 20% final concentration as according to cell line supplier's recommendation, Thermo Scientific, Cat: 10082147) and Penicillin/Streptomycin (1% final concentration from a 100X stock, Thermo Scientific, Cat: 15140163). BT549 cells were supplemented with insulin (Thermo Scientific, Cat: 12585-014) at 0.023 U/ml. HS578T was supplemented with insulin at 0.01 mg/ml. Cell culture media was filter-sterilized via 0.2 µm Nalgene™ Rapid-Flow™ Sterile Disposable Filter Units with PES Membrane, 500 mL capacity (Thermo Scientific, Cat: 566-0020) or 1000 mL capacity (Thermo Scientific, Cat: 567-0020). Cells were grown in manufacturer's recommended media with the following exceptions: MDA-MB-468 cells were grown in DMEM, HPAC in IMDM, SW48 in McCoy's 5A and SW900 in RPMI containing media.

Primary bronchial/tracheal epithelial cells (Cat: PCS-300-010), epidermal keratinocytes (Cat: PCS-200-011) and cervical epithelial cells (Cat: PCS-480-011) were purchased from ATCC. Primary bronchial tracheal epithelial cells were grown in airway epithelial cell basal media (ATCC, Cat: PCS-300-030) supplemented with a bronchial epithelial cell growth kit (ATCC, Cat: PCS-300-040). Primary epidermal keratinocytes were grown in dermal cell basal media (ATCC, Cat: PCS-200-030) supplemented with keratinocyte growth kit (ATCC, Cat: PCS-200-040). Primary cervical epithelial cells were grown in cervical epithelial cell basal medium (ATCC, Cat: PCS-480-032) supplemented with cervical epithelial growth kit (ATCC, Cat: PCS-480-042).

All cells were grown at 37$^{o}$C with 5% $CO_2$ in 75 $cm^2$ (Corning, Cat: 430641U) or 150 $cm^2$ (Corning, Cat: 430825) vented cap sterile cell culture flasks or 15 cm plates (Thermo Scientific, Cat: 150468)

#### Method for generating the Osimertinib resistant HCC827 cells

HCC827 cells were cultured in 500 nM osimertinib. Media and osimertinib was replaced every 3-4 days. Once the cells began to double in 500nM osimertinib (~day 35) the concentration was increased to 1 µM. Cells were allowed to expand and then frozen back. Resistance to osimertinib was confirmed in a dose response assay.

### General Synthetic Information

Riboflavin tetraacetate (RFT) photocatalyst utilized in these studies (Supplemental Figure 5) was synthesized as previously described in Oslund, R.C. et al.[5] The iridium photocatalyst and biotin-containing diazirine probe (Ar-PEG3-Biotin) utilized in these studies (Supplemental Figure 5) was synthesized as previously described in Geri, J.B. et al.[6]

### Preparation of secondary antibody-photocatalyst conjugate

350 µL of polyclonal goat α-mouse IgG (Millipore, Cat: AP124) or polyclonal goat α -rabbit IgG (Millipore, Cat: AP132) was combined with 50 µl of 1M sodium bicarbonate buffer (pH 8.5, Thermo Scientific, Cat: J60408.AK) in a Low Protein Binding tube. 4 µL of 100 mM azidobutyric acid NHS ester (prepared in DMSO, Broadpharm, Cat: BP-22526) was added and the reaction mixture was incubated for 1.5 hours at room temperature (RT) in the dark. After 1.5 hours, an additional 4 µL of 100 mM azidobutyric acid NHS ester linker was added, and the sample was incubated for 1.5 hours at RT in the dark. In the meantime, a Zeba Spin desalting column (2 mL column, 40,000 MWCO, Thermo Scientific, Cat: 87769 or A57762) was prepared by first removing the storage solution (centrifuged 2,000xg for 3 min 4°C). The column was then primed by three washes of 1 ml 50 mM Tris pH 7.5 and spun at 2,000xg 5 min 4°C after each addition except for the last spin which was extended to 10 min. After the second incubation of antibody and azidobutyric acid NHS ester, the sample was buffer exchanged into 50 mM Tris pH 7.5 using the Tris primed Zeba Spin desalting column and spun at 2,000xg 2 min 4°C. The antibody-azide conjugate was transferred to a new tube for click chemistry azide-alkyne cycloaddition of the photocatalyst using the Click-iT™ Protein Reaction Buffer Kit (Thermo Scientific, Cat: C10276). 15 µL photocatalyst (from 5 mM stock in DMSO) was added to the antibody-azide conjugate and mixed. 15 µL of copper sulfate and 15 µL of additive 1 was added and mixed and let incubate for 1.5 min at RT. Following incubation, 30 µL of additive 2 was added to the reaction mixture and incubated in the dark for 30 min at RT. After, the sample was buffer exchanged into DPBS (Thermo Scientific, Cat: 14190144) using a Zeba Spin desalting column primed in the same manner as above except with DPBS. In some instances, the conjugated antibodies were spun at 16,000xg 10 min 4°C to remove precipitates. The final protein concentration of the secondary antibody-photocatalyst conjugate was determined using the Pierce™ BCA Protein Assay kit (Thermo Scientific, Cat: 23227) according to the manufacturer's instructions. The photocatalyst concentration was determined by measuring the absorbance (350 nm for Ir and 450 nm for RFT) and compared to a standard curve consisting of known free photocatalyst concentrations. The micromolar concentration of photocatalyst was divided by the micromolar concentration of antibody to determine the antibody:photocatalyst ratio. A ratio of 1:8 was routinely obtained.

### Preparation of direct antibody-photocatalyst conjugates

**Azide conjugation**

CDCP1 and EGFR targeted micromapping using directly conjugated antibody-photocatalyst conjugates was performed using EGFR×Fc and CDCP1×Fc binders as well as a non-binding anti-RSV×Fc control. All antibodies were produced in ExpiCHO-S cells (Thermo Scientific, Cat: A29127) in ExpiCHO expression medium (Thermo Scientific, Cat: A2910001). These were purified using an AKTA pure. All samples were first purified using PrismA HiTrap affinity capture (Cytiva, Cat: 17549854), neutralized with 1M NaPi pH 7.0 and followed by SEC purification over a Superdex 200 increase (Cytiva, Cat: 28990944) in PBS (Fisher Scientific, Cat: SH3025602). Four antibodies were produced in-house; an anti-EGFR (cetuximab, sequence obtained from US patent 7,060,808[7]), anti-CDCP1 clone 41A9 (sequence obtained from US patent 11,702,481[8]), anti-RSV (sequence obtained from https://go.drugbank.com/drugs/DB00110) and an antibody fragment referred to as "Fc" which is comprised of a human IgG1 antibody sequence from just above the hinge (E215) to the C-terminus (G446). Each of these carried mutations in the CH3 domain specific for conducting Fab-arm exchange[9]. The Fc-antibody was directly conjugated with azide by buffer exchanging antibody into PBS using Zeba desalting columns. 10% v/v of 7.5% $NaHCO_3$ was added to adjust the pH (Hyclone, Cat: sh30033.01). 3-Azidopropanoic acid NHS ester (Broadpharm, Cat: BP-23800) was prepared to 50 mM stock solution in DMSO. 20 molar equivalents was used as a challenge ratio and added to the solution for 90 minutes on a shaker platform protected from light. Sample was buffer exchanged into PBS using Zeba desalting columns to remove excess azide. Fab arm exchange was then performed on the azide conjugated Fc with each of the targeting arms (EGFR, CDCP1 and RSV). Equimolar amounts of each antibody were added to the reaction and adjusted to a 25 mM concentration of 2-mercaptoethylamine-HCl (2-MEA, Sigma-Aldrich, Cat: 30078). The samples were incubated at 37°C for 2 hours, then buffer exchanged with PBS to remove 2-MEA. This azide labeling of the Fc approach allows for high throughput production of azide conjugated bispecifics while ensuring there is no disruption to the binding sites on the targeting arm.

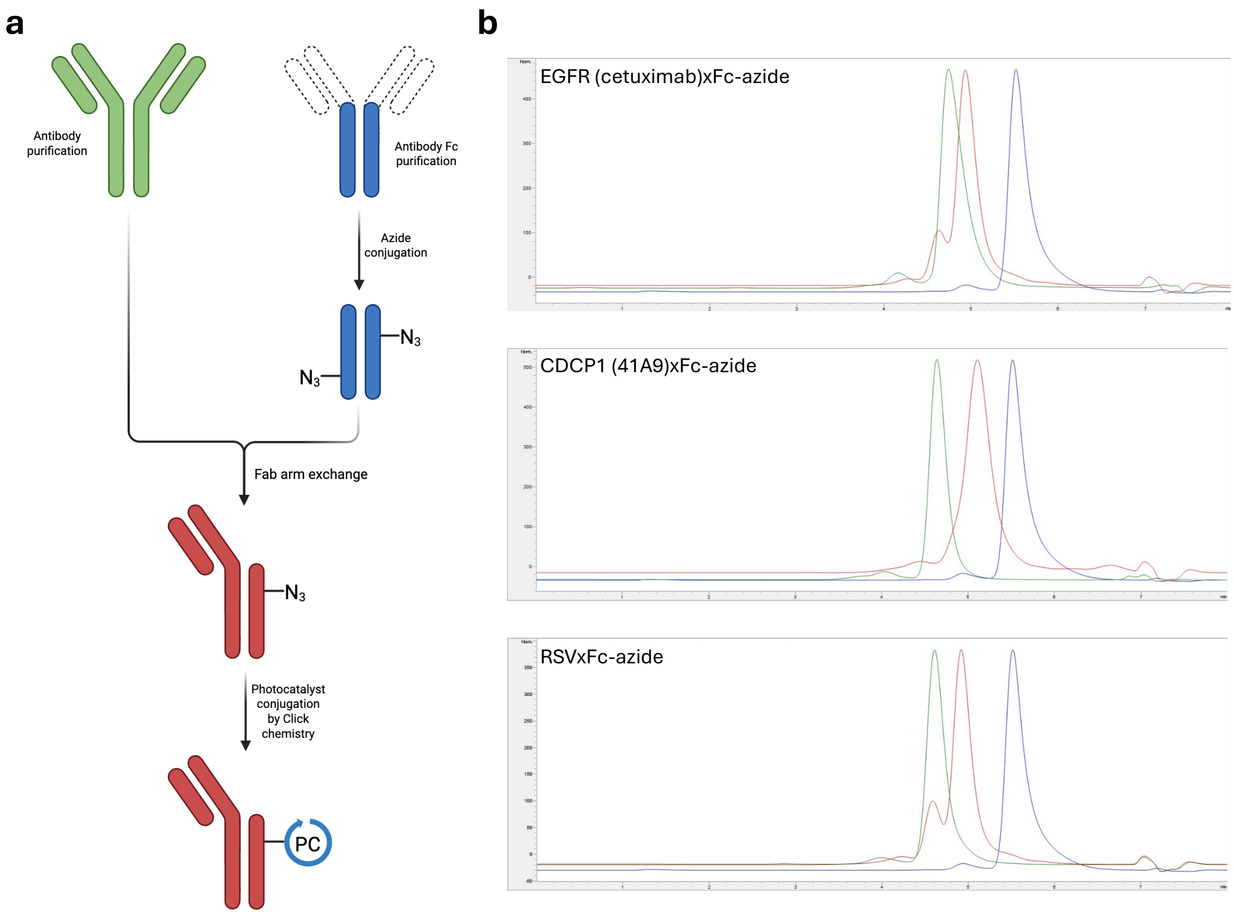

Above figure shows **a)** the general workflow for azide conjugation of bispecifics. Created in BioRender. May, C. (2026) https://BioRender.com/s9x2nl8. **b)** Analytical SEC traces of the three azide conjugated bispecifics showing EGFR×Fc-azide (top), CDCP1×Fc-azide (middle) and RSV×Fc-azide (bottom). The parental targeting arm in green, the bispecific in red, and the azide labeled parental Fc arm in blue.

**Photocatalyst click labeling**

200 µL of 10 µM targeting arm×Fc-azide antibody was covalently conjugated to the photocatalyst using the Click-iT™ Protein Reaction Buffer Kit and purified using a Zeba Spin desalting column as described above. An isotype control (anti-RSV) was prepared in parallel using the same protocol. The final protein concentration of the direct antibody-photocatalyst conjugate was determined using the Pierce™ BCA Protein Assay Kit - Reducing Agent Compatible (Thermo Scientific, Cat: 23250) according to the manufacturer's instructions. The photocatalyst concentration was determined as described above. Ratios of 1:9 and 1:5 were routinely observed for Ir and RFT, respectively.

### Flow cytometry analysis of target proteins

Cells were harvested with Accutase (BioLegend, Cat: 423201), centrifuged at 800xg 5 min 4°C, resuspended in DPBS, counted using a Beckman Coulter Vi-CELL XR Cell Viability Analyzer and aliquoted at 200,000 cells per well in a 96-well U-bottom plate (Fisher Cat: 351177) in 100 µL DPBS. 100 µL DPBS containing LIVE/DEAD™ Fixable Near-IR Dead Cell Stain (Thermo Scientific, Cat: L10119) at a 1:1000 dilution was added to the cells and then incubated for 30 min at RT in the dark. The cells were centrifuged at 800xg 5 min 4°C and washed once with 200 µL FACS buffer (DPBS containing 2% FBS and 2 mM EDTA). Cells were resuspended in 100 µL containing 25nM primary antibody (see "List of primary antibodies used for microenvironment mapping" below) and incubated for 30 min at 4°C. 100 µL FACS buffer was added and the cells were centrifuged to remove the primary. The cells were then washed once in 200 µL FACS buffer before resuspension in 100 µL of FACS buffer containing a 1:250 dilution of PE secondary (goat anti-mouse BioLegend Cat: 405307 or donkey anti-rabbit BioLegend Cat: 406421) for 30min at 4°C. 100 µL FACS buffer was added and the cells were centrifuged to remove the secondary. The cells were then washed once in 200 µL FACS buffer before resuspension in 50 µL fixation buffer (BioLegend, Cat: 420801) for 30 min at RT in the dark. 150 µL FACS buffer was added and the cells were centrifuged. The cells were then washed once in 200 µL FACS buffer before resuspension in 150 µL FACS buffer. Samples were then run on a BD LSRFortessa X-20 with BD FACSDiva software or kept 4°C overnight for next day analysis. Data was analyzed using FlowJo, v10.

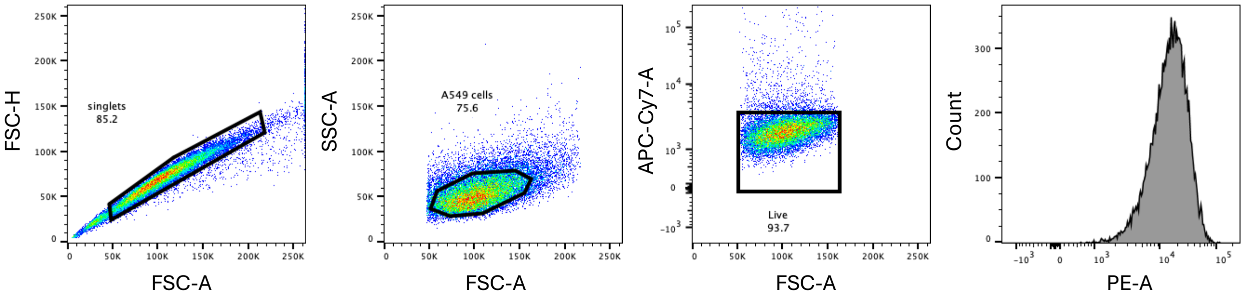

Above figure shows representative flow cytometry gating strategy. Single cells were selected by gating on FSC-H versus FSC-A to remove doublets. The target cells were selected by SSC-A versus FSC-A. Lastly, the viable cells were selected by exclusion of positive staining from the LIVE/DEAD™ Fixable Near-IR Dead Cell Stain using APC-Cy7-A. Either a histogram or geometric mean of the PE-A channel was used for further analysis.

### Flow cytometry competitive binding assay

NCI-H1975 cells were harvested with Accutase, centrifuged at 800xg 5 min 4°C, resuspended in DPBS, counted using a Vi-CELL XR Cell Viability Analyzer and aliquoted at 2 million cells each to Low Protein Binding tubes. Cells were spun down and resuspended in 500 µL cold DPBS. 5 µg primary antibody was next added: for EGFR, cetuximab antibody (prepared as described above, obtained from US patent 7,060,808[7]) or human isotype antibody (BioLegend, Cat: 403502) was used; for CDCP1, CDCP1 antibody (BioLegend, Cat: 324002) or mouse isotype antibody (BD, Cat: 556648) was used. For the competitive sample, 25 µg competing antibody was added immediately before the primary. In the case of EGFR competition, EGFR (BD, Cat: 556648) was used; for CDCP1 competition, CDCP1 41A9 (prepared as described above, obtained from US patent 11,702,481[8]) was used. Samples were incubated 1.5 hours 4°C on a rotisserie. Cells were spun down at 800xg 5 min 4°C and washed once with 1 mL cold DPBS before resuspension in 500 µL cold DPBS containing Zombie Green Viability dye (1:500 dilution from stock prepared according to manufacturer's instructions, BioLegend, Cat: 423111). Secondary antibody specific for the primary, but not the competing antibody was added – for EGFR competition, goat anti-Human APC (1:100 dilution, R&D Systems, Cat: F0135); for CDCP1 competition, goat anti-Mouse APC (1:500 dilution, Thermo Scientific, Cat: A865). Samples were incubated for 30 min at 4°C on a rotisserie. Cells were spun down at 800xg 5 min 4°C and washed once with 1 mL cold DPBS before resuspension in 500 µL cold DPBS and transfer to FACS tubes. Samples were then run on a BD Accuri C6 Plus with CSampler Plus. Data was acquired with the BD Accuri CSampler Plus software and analyzed using FlowJo, v10 using the above gating strategy except the Viability gate utilized the FITC-A channel and the primary antibody was detected with the APC-A channel.

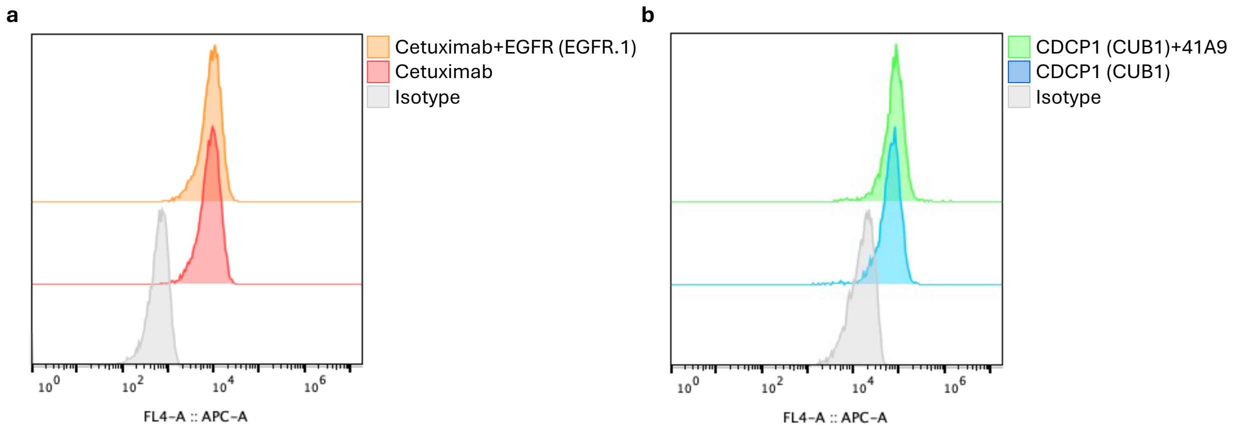

Above figure shows flow cytometry competitive binding analysis of the binders used for EGFR and CDCP1 micromapping and the binders used for direct conjugate mapping in Supplementary Figure 16 to confirm that different epitopes are engaged. **a)** EGFR competition experiment where cells were incubated with cetuximab alone (red histogram) or cetuximab with an excess of anti-EGFR antibody (clone EGFR.1, orange histogram) that was used for the micromapping studies. No reduction in binding was observed indicating that each antibody recognizes separate epitopes. **b)** CDCP1 competition experiment where cells were incubated with anti-CDCP1 antibody (clone CUB1, blue histogram) that was used for micromapping studies or anti-CDCP1 (clone CUB1) with an excess of CDCP1 41A9 (green histogram). No reduction in binding was observed indicating that each antibody recognizes separate epitopes.

### Micromapping of surface proteins in live cells for LC-MS/MS Analysis

For two antibody-based micromapping, cells were harvested using Accutase and centrifuged at 800xg 5 min 4°C before being resuspended in DPBS and counted using a Vi-CELL XR Cell Viability Analyzer. 10-20 million cells were aliquoted to each Low Protein Binding tube with each condition done in triplicate (n=3). The cells were spun down and resuspended in 1 mL cold DPBS containing a targeting primary antibody or an appropriate isotype control (see "List of primary antibodies used for micronenvironment mapping" below) at 1 µg antibody for every million cells and incubated at 4°C for 30 min on a rotisserie. After the primary incubation, cells were spun down and washed twice with 1 mL cold DPBS. The cells were then resuspended in 1 mL cold DPBS containing the secondary antibody conjugated to photocatalyst at 1 µg antibody for every million cells and incubated at 4°C for 30 min on a rotisserie. Cells were then spun down and washed twice with 1 mL cold DPBS. The cells were resuspended in 1 mL cold DPBS containing 250 µM biotin probe (biotin tyramide from ApexBio, Cat: A80111000). The samples were put in a bio-photoreactor[10] for 2 min (for RFT photocatalyst samples) or 3 min (for Ir photocatalyst samples) and irradiated at full intensity. The samples were then spun down and washed twice with 1 mL cold DPBS. The cells were then lysed in one of the two following methods: 1) 1 mL membrane permeabilization buffer (MEM-PER Plus Membrane Fractionation Kit, Thermo Scientific: 89842) with 1X protease inhibitor tablet (Sigma-Aldrich, Cat: 4693159001). The samples were then allowed to rotate for 20 min at 4°C before centrifugation at 16000xg for 15 min 4°C. The supernatant was discarded and the membrane pellet was resuspended in 300 µL RIPA (Thermo Scientific, Cat: 89901) with 1% sodium dodecyl sulfate (SDS, prepared from a 20% stock, Quality Biological, Cat: 351-066-101) and 1X protease inhibitor tablet, sonicated and boiled at 95°C for 5 min. 1 mL RIPA buffer was added and the lysate was sonicated again to homogenize. 2) Alternately, the cells were lysed in 1 mL RIPA buffer containing 1X protease inhibitor tablet and 1:1000 dilution of benzonase (Sigma-Aldrich, Cat: 70664-3) and incubated for 15 min 4°C on a rotisserie. After lysis, the protein concentrations were measured by BCA assay stored at -80°C until the bead enrichment.

**List of primary antibodies used for microenvironment mapping**

| Antibody | Clone | Source | Cat: |
|---|---|---|---|
| Mouse Isotype | MOPC-21 | BD Biosciences | 556648 |
| Rabbit Isotype | 60024B | R&D Systems | MAB1050-500 |
| AXL | 108724 | R&D Systems | MAB154-100 |
| CDCP1 | CUB1 | Biolegend | 324002 |
| DDR1 | EPR22316-508 | Abcam | ab255810 |
| EGFR | EGFR.1 | BD Biosciences | 555996 |
| EPHA2 | 371805 | R&D Systems | MAB3035 |
| FGFR2A | 1057909 | R&D Systems | MAB11119-100 |
| HER2 | 24D2 | Biolegend | 324402 |
| HER3 | 66223 | R&D Systems | MAB3481 |
| IGF1R | 33255 | R&D Systems | MAB391-100 |
| MET | 95106 | R&D Systems | MAB3582 |
| PDGFRB | 28D4 | BD Biosciences | 558820 |
| PTK7 | OTI2E7 | Thermo Scientific | MA5-25774 |
| ROR1 | 4A5 | BD Biosciences | 564464 |

The directly conjugated micromapping was carried out as described above with the following changes. Instead of primary and secondary antibody, a directly conjugated antibody was added at equimolar amounts equivalent to 1µg of full-length antibody for every 1 million cells. In addition, a control (anti-RSV) directly conjugated to photocatalyst was used for the isotype samples. Lysis was carried out only using the RIPA buffer containing 1X protease inhibitor tablet and 1:1000 dilution of benzonase.

Following labeling, biotinylated proteins were enriched using streptavidin magnetic beads using either a manual or automated (KingFisher Apex, Thermo Scientific) workflow. In all cases, enrichment followed a common binding and wash procedure, with samples processed either as bead-retained material or by competitive biotin elution, as specified below.

For bead enrichment, 100 or 250 µL of streptavidin magnetic beads (Thermo Scientific, Cat: 88817) were washed twice with 1 mL RIPA. Equal protein amounts were added to the beads and the differences in volume were made up with RIPA buffer. The tubes were allowed to rotate 3 hours at RT. The beads were collected on a magnetic rack and washed three times with 1 mL DPBS containing 1% SDS. Next, the beads were washed three times with 1 mL DPBS containing 1 M NaCl (prepared from a 5M stock, Research Products International [RPI], Cat: S24600-500.0) followed by three washes with 1 mL DPBS containing 10% ethanol (200 proof stock, Fisher, Cat: BP2818-500) and then one wash with RIPA buffer.

After washing, enriched proteins were processed by one of two endpoints. In bead-retained workflows, beads were carried forward directly for downstream proteomic processing. In elution workflows, proteins were released from the beads by boiling at 95 °C for 10 min in 4× Laemmli buffer (Bio-Rad, Cat: 1610747) supplemented with 20 mM DTT (RPI, Cat: D11000-10.0) and 25 mM biotin (Sigma-Aldrich, Cat: 14400-1G).

Automated bead enrichment was performed using a KingFisher system following the same binding and wash sequence, with wash volumes reduced to 900 µL. In this format, the final RIPA wash was replaced by release of beads into 180 µL of 0.2 M HEPES buffer (pH 8.5; prepared from 0.5 M stock, Thermo Scientific, Cat: J63218.AK). Plates were sealed and samples stored at −80 °C prior to proteomic analysis.

#### Protein Extraction and Digestion for LC-MS/MS Analysis

In early experiments, proteins eluted from the beads were TCA precipitated and washed with ice-cold acetone. Dried protein pellets were reduced and alkylated by resuspending in 4 M urea with 5 mM tris(2-carboxyethyl) phosphine (TCEP) and 20 mM chloroacetamide (CAA) and incubating at 20°C for 20 min. Proteins were digested by adding an equal volume of 100 mM Tris-Cl, pH = 8.5 with 200 ng lysyl endopeptidase (lysC, FUJIFILM Wako, Cat: 125-05061) and shaking overnight at 30°C then adding an equal volume of 50 mM Tris-Cl, pH=8.5, with 200 ng trypsin (Promega, Cat: V5111) and shaking for 6 hr at 37°C. For on-bead digestion experiments, bead-enriched lysates were reduced and alkylated on the beads with 5 mM TCEP and 20 mM CAA for 20 min. The beads were then washed with 0.2 M HEPES, pH=8.5 buffer before a 4-hour digestion with lysC and overnight digestion with trypsin as above. Digested peptides were pipetted off the beads and 1x 0.2M HEPES wash of the beads was performed and added to the digested peptides to recover any remaining peptides from the beads. This protocol originally was conducted in microcentrifuge tubes with a magnet rack and manual pipetting and bead mixing. It was later modified to enable 96-well format automation using the KingFisher. The separate digestions with lysC and trypsin were combined into a single four-hour step performed on the KingFisher with mixing at 37°C. There was no noticeable impact on digestion. Following digestion, peptides were labeled with 250 µg tandem mass tag (TMTpro; Thermo Scientific, Cat: A52045) isobaric reagents for 2 hours at room temperature. Labeling efficiency was checked by pooling 5µl from each sample within a single plex. For the full experiment, all samples were quenched with hydroxylamine (0.5%) and pooled. Mixes were acidified with TFA (2%) and desalted with SPE-C18 columns (Waters Sep-Pak). Desalted TMT mixes were dried down in a SpeedVac concentrator (Thermo Scientific) and fractionated using the high pH reverse-phase peptide fractionation kit (Pierce) into 13 fractions (5%-50% acetonitrile in 0.1% triethylamine). Every fourth fraction was combined to make 4 pools (1-5-9-13, 2-6-10, 3-7-11, 4-8-12) which were then dried down in a SpeedVac and resuspended in 5% formic acid for LC-MS/MS analysis.

### LC-MS/MS-based Proteomic Analysis of Micromapping Experiments

**LC-MS Data Collection**

All mass spectra were acquired on an Thermo Scientific Orbitrap Eclipse Tribrid Mass Spectrometer coupled to a Vanquish Neo UPLC (Thermo Fisher). Peptides were separated on a 25 cm Aurora Ultimate C18 capillary column (Ionopticks, Cat: AUR4-25075C18) using a 76 min linear gradient from 8% to 28% buffer B (80% Acetonitrile [ACN], 0.1% formic acid [FA]), followed by 5 min from 28-38% B and then 3 min from 38-55% B. The column was held at 60°C with a PRSO-V2 column oven (Sonation). The mass spectrometer was operated in a data dependent mode using the real-time search (RTS) feature to trigger synchronous precursor selection (SPS) of positively identified peptides for $MS^3$ analysis of TMT reporter ions[11]. The scan sequence began with a full survey scan in the Orbitrap (resolution = 120,000; mass range = 400-1600 *m/z*; max injection time = 246 ms; automatic gain control [AGC] target = 4e5; dynamic exclusion for 40 seconds with a +/- 10 ppm window excluding all isotope peaks and collecting only one charge state peptide per precursor). Using a cycle time = 3 s, the most intense precursor ions were selected for $MS^2$ analysis in the linear ion trap via collisional-induced dissociation (CID) (normalized collision energy [NCE] = 32%; activation time = 20 ms; activation Q = 0.25; scan rate = 'rapid'; max injection time = 50 ms; quadrupole isolation window = 1.0 m/z; AGC target = 2e4). $MS^2$ spectra were searched in real-time to trigger MS3 collection[12]. The real-time search used a database of 20,420 canonical human protein sequences (downloaded from uniprotkb on 8/19/2024: www.uniprot.org/proteomes/UP000005640). Search parameters allowed for 1 missed cleavage; fixed modifications of +304.207 (TMTpro) on lysines and peptide amino termini and +57.021 on cysteines; a dynamic modification of +15.999 on methionine. Confidently identified peptides were selected for synchronous-precursor-selection (SPS) $MS^3$ scans to capture TMT reporter ions. Up to eight $MS^2$ product ions were selected for high energy collisional-induced dissociation (HCD) and analysis in the Orbitrap (NCE = 45%; resolution = 50,000; max injection time = 1000 ms; AGC target = 5e5; scan range = 110 – 140).

**LC-MS Data Analysis**

All data processing steps were performed with Thermo Proteome Discoverer 3.0. Searching was done using SEQUEST HT with Inferys rescoring and a database of canonical human protein sequences (downloaded from uniprotkb either on 5/23/23 or 8/19/2024: www.uniprot.org/proteomes/UP000005640) containing common contaminants and an equal size decoy database. Searches required fully tryptic ends with one or fewer missed cleavages; a precursor mass tolerance of 20 ppm and a fragment ion tolerance of 1.005 Da; oxidation of methionine (15.9949 Da) and TMTpro on lysines and N-termini of peptides (304.207) were set as dynamic modifications; and a static modification of carbamidomethylation of cysteines (57.021).

Peptide-spectrum matches were filtered using linear discriminant analysis[13] and adjusted to a 1% (strict) and 2% (relaxed) peptide false discovery rate (FDR)[14].

### LC-MS/MS-based Proteomic Analysis of Tumor Cells and Patient Derived Tissue Samples

**Cell Extract and Peptide Preparation**

Whole-cell extract (WCE) generation was performed all on cell systems listed in Supplementary Figure 3a. Three technical replicates of 5e5 cells were lysed in parallel in 96-plates. Before aliquoting, washed cell-pellets were re-suspended in ice-cold PBS at 1.1e7 cells/ml and 45 µl was transferred to the lysis tube or plate. Cells were pre-treated for 10 min on ice with Pierce Universal Nuclease (125 U total, Cat: 88701) and lysed by the addition of 50 µl of 2x lysis buffer (100 mM HEPES pH = 8.5, 1% SDS, 2% sodium deoxycholic acid, 2 mM MgCl2, 100 mM NaCl, 10 mM TCEP, 40 mM CAA, 1x protease inhibitor). The cells were incubated with mixing at 37°C for 10 min. 10 µl of 20% SDS was added to bring the final concentration to >2% and the samples were mixed for an additional 10 min at 70°C. Protein concentrations were measured using the Pierce reducing agent compatible BCA kit (Thermo Scientific, Cat: 23250).

Frozen tumor and matched normal tissue samples analyzed in Supplementary Figure 19 were obtained from Audubon Bioscience. Samples were cut on ice with a clean scalpel, blotted dry to remove excess liquid, weighed, and stored at -80°C. For WCE analysis, aliquots of ~10 mg were lysed in microcentrifuge tubes using the procedure described above for cultured cells with the following alterations. Prior to lysis, tissue samples were washed 4x with ice-cold PBS to remove blood and preincubated with 20 µl of nuclease solution before adding 100 µl of 1x lysis buffer and grinding with a plastic SpiralPestle (RPI, #2999017) driven by a rotary pestle motor (Cole-Parmer, Cat: EW-44468-25) for 30 s. The pestle was washed with an additional 100 µl of lysis buffer to remove residual lysate and the samples were mixed at 37°C for 10 min before the addition of 1/10 volume 20% SDS and a final mixing incubation at 70°C for 10 min. Samples were cleared by centrifugation and the supernatant was transferred to fresh tubes. Protein concentrations were measured by BCA analysis as previously described. Larger aliquots (~30 mg) of tissue samples were also processed to isolate plasma membrane fractions using the Minute Plasma Membrane fractionation kit (Invent, Cat: SM-005) per the manufacturer's protocol. For all cultured cell and tissue samples, we digested 10 µg of total protein using the SP3 method[15] with lysC and trypsin. The resulting peptides were desalted using Supel Swift HLB dispersive tips (DPX Technologies, Cat: DPX170501), eluted with 70% ACN, dried in a SpeedVac, and resuspended in 5% FA.

**LC-MS Analysis**

200-500 ng of each sample was analyzed using a 90 min data independent acquisition (DIA) method on the same column and LC-MS system described above. The separation gradient included a 73 min segment ranging from 5-28% B followed by 7 min from 28-36% and then 4 min from 36-50%. Precursor scans were collected from 380-980 *m/z* in the orbitrap with a resolution = 60,000 and AGC = 4e5. Sixty, 10 *m/z* wide windows, with 1 *m/z* overlap were selected for HCD fragmentation $MS^2$ scans in the orbitrap at resolution = 15,000 with AGC target = 1e5 and a 40 ms maximum ion accumulation time.

**DIA LC-MS Data Analysis**

DIA raw files were analyzed using Spectronaut (v. 20.1; Biognosys AG). Data were searched in directDIA+ mode against a human protein database (UniProtKB UP000005640, release 2024_08; 20,420 entries). For peptide identification, the Pulsar search engine was used with the following parameters: [Trypsin/P] digestion, [two] maximum missed cleavages. Fixed modification was set to Carbamidomethyl (C), and variable modifications included Oxidation (M) and Acetylation (Protein N-term). Identification stringency was controlled using a target-decoy approach with a 1% False Discovery Rate (FDR) at both the precursor and protein group levels. Quantification was performed at the MS2 level, utilizing the maxLFQ algorithm for protein group intensity calculation. Data were normalized using the Global Normalization strategy. Automated non-linear iRT and mass calibration were applied to each run.

Protein abundance data derived from LC–MS/MS–based proteomic profiling of tumor cells were used as input features for downstream computational analyses (see Graph ML Protein Proximity Prediction section).

Proteomic data of tumor vs matched normal tissue samples were used for measuring CDCP1 levels (Supplementary Figure 19).

### Bioinformatic Analysis of Micromapping Experiments

**Differential Enrichment Analysis**

Primary bioinformatic analysis of LC-MS/MS data was performed in the R statistical computing environment. Peptide-level abundance data was used to identify the number of peptides corresponding to each protein. To normalize for loading differences, peptide abundance was normalized to the summed total abundance for each sample; these totals were averaged, and individual normalized values were rescaled by this average. Peptide-level data was then merged to protein-level data by calculating the median of all peptides assigned to a given protein.

Differential protein enrichment was subsequently assessed using the limma R package. Log2-transformed summed intensity values were used to calculate statistical significance, reporting t-statistics and p-values. To account for multiple hypothesis testing, p-values were adjusted using the Benjamini-Hochberg (BH) procedure. Results were visualized using Seaborn, generating volcano plots that display log2 fold change against the negative log10 transformed adjusted p-values.

### Statistical Methods and Network Analysis

**Proximity Interaction Networks**

Global RTK proximity network was constructed using the NetworkX Python library, where edges connected experimental target proteins to enriched proteins. We applied permissive enrichment thresholds ($\log2FC > 0.1$, $q\text{-value} < 0.2$) to define edges, ensuring the inclusion of lower-affinity or transient interactions. The resulting network topology was exported in GEXF format and spatialized in Gephi using the ForceAtlas2 layout algorithm (LinLog mode).

The known RTK-RTK interaction network in Figure 2c was constructed in NetworkX from previously reported[16], literature-curated RTK heterointeractions, augmented with high-confidence intra-family interactions from STRING (experimental score > 0.7). In contrast, analyses in Figures 2e–f were evaluated against a broader, more inclusive reference set integrating multiple interaction databases (STRING, CORUM, BioGRID, IntAct), enabling assessment against less stringent but more comprehensive interaction annotations. Metrics were calculated as described in Evaluation Metrics. The computational graph was exported to Gephi for final formatting.

**Overlap and Similarity Analysis**

To systematically compare RTK interactomes across targets, we generated a 2-dimensional array of pairwise comparisons. Overlaps were visualized as area-proportional Venn diagrams using matplotlib-venn, with circle sizes weighted by the total number of enriched proteins. The Jaccard index was computed to quantify set similarity ($J = |A \cap B| / |A \cup B|$). To facilitate interpretation, conditions were ordered by Euclidean distance, placing phenotypes with the greatest overlap in proximity.

**Score Normalization**

Raw t-statistics were normalized using robust z-score transformation applied independently within each experiment. The robust z-score was computed as:

$$z_i = \frac{t_i - \mathrm{median}(t)}{\mathrm{MAD}(t)}$$

where MAD denotes the Median Absolute Deviation, scaled by factor 1.4826 for consistency with the standard deviation of a normal distribution [scipy.stats.median_abs_deviation with scale='normal']. This per-experiment normalization equalizes distributional differences across experiments (loud vs. quiet targeted protein effects), enabling fair cross-experiment comparisons.

**Hit Calling**

Proteins were flagged as enriched ("hits") if their normalized score exceeded a threshold of 2.0 MAD units:

$$\mathrm{hit}_i = 1[z_i > 2.0]$$

An alternative statistics-based criterion required both log2 fold-change > 1.0 and Benjamini-Hochberg adjusted p-value ≤ 0.05.

**Pairwise Correlation Analysis**

Spearman rank correlations were computed between all protein pairs across experiments using pairwise-complete observations. Correlations were computed only for pairs with at least 10 co-observations (experiments where both proteins had non-missing scores). The correlation coefficient ρ was computed as the Pearson correlation of ranked values:

$$\rho = \frac{\sum_i (R_{x,i} - \bar{R}_x)(R_{y,i} - \bar{R}_y)}{\sqrt{\sum_i (R_{x,i} - \bar{R}_x)^2 \sum_i (R_{y,i} - \bar{R}_y)^2}}$$

Where $R_{x,i}$ denotes the rank of protein *x* in experiment *i* among valid observations. Ties were resolved using average ranking. P-values were computed analytically using the t-distribution approximation:

$$t = \rho\sqrt{\frac{n-2}{1-\rho^2}}, \quad p = 2 \cdot P(T > |t|), \quad T \sim t_{n-2}$$

Multiple testing correction was performed using the Benjamini-Hochberg procedure at FDR $\alpha$ = 0.05 with monotonicity enforcement.

**Network Clustering**

Protein modules were identified using the Louvain community detection algorithm [NetworkX] on weighted undirected graphs constructed from significant correlations. Edge weights corresponded to Spearman $\rho$ values. Parameters: correlation threshold ≥ 0.3 for edge inclusion, resolution parameter = 1.0, minimum cluster size = 5 proteins, random seed = 42 for reproducibility.

Community clusters, such as the one depicted in Figure 3, were constructed by drawing edges between core community proteins with high correlation ($\rho > 0.4$). Additional edges connect highly correlated proteins of interest to core members. Edges are annotated as documented in reference databases or as potentially novel.

**Evaluation Metrics**

Predicted protein associations were evaluated against reference databases using:

- **Precision**: TP / (TP + FP), proportion of predicted pairs in reference
- **Recall**: TP / (TP + FN), proportion of reference pairs recovered
- **Recall@k**: Recall among the top k predicted pairs by correlation strength

Evaluations were stratified by pair type: RTK-RTK pairs, RTK-target pairs (involving experimental targets), RTK-non-target pairs, and all detected pairs.

To evaluate whether correlated protein pairs are enriched for known interactions, we compared predicted pairs against a random baseline. For each reference database, we calculated precision as defined above. We then generated 100 random samples of protein pairs, matched in size to the predicted set, drawn from the same protein universe. Fold enrichment was computed as the ratio of predicted precision to mean background precision. Statistical significance was assessed using an empirical p-value: the proportion of random samples achieving precision equal to or greater than the predicted precision, with a +1 correction applied to both numerator and denominator to avoid zero p-values ($p = (k + 1) / (n + 1)$, where k is the count of random samples ≥ predicted and n = 100 iterations).

**Reference Interaction Databases**

Protein-protein interaction references were obtained from STRING (v12), BioGRID (human physical interactions), IntAct (human molecular interactions), and CORUM (manually curated protein complexes, v5.1). For STRING interactions, only high-confidence associations (combined score ≥ 0.7) were retained. CORUM complexes were expanded to all pairwise protein combinations within each complex (minimum complex size: 2 proteins). Interactions were normalized such that protein_a ≤ protein_b alphabetically to ensure consistent pair representation.

**Visualization**

Publication figures were generated at 300 DPI in both PNG and SVG formats. Hierarchical clustering used Ward linkage with Euclidean distance on correlation matrices. Kernel density estimates used bandwidth adjustment factor 4.5 for score distributions.

### Graph ML Protein Proximity Prediction

To identify candidate proximity-associated pairs supported by spatial proximity in micromap experiments, we developed graph-based machine learning models that integrate proteomic features with known interaction networks. Model outputs were used to prioritize protein pairs exhibiting consistent proximity behavior across experimental conditions for further biological interpretation. Proximity predictions were used solely to prioritize candidates and do not imply direct physical interaction.

**Computational Graph Construction**

We constructed a multi-graph dataset where each graph represents a single micromap experiment, with nodes representing proteins and edges representing known high-confidence interactions from STRING. Feature vectors comprised of two components: our protein expression data (z-score normalized per cell line after log transformation) and our spatial proximity data (Inter-Quartile Range normalized). To prevent information leakage, known interaction edges used as prediction targets were excluded from the graph structure during feature computation and training.

Spatial proximity data were compiled from our complete set of 248 micromap experiments conducted across distinct experimental conditions. For each experiment proteins were filtered using a proximity $\log_2$ fold-change threshold of ≥ 0.3.

Protein expression data were compiled from our protein abundance measurements captured on all cell systems listed in Supplementary Figure 3a. See LC-MS/MS-based Proteomic Analysis of Tumor Cells and Patient Derived Tissue Samples  above.

### Model Architecture and Training

Analysis was restricted to protein pairs co-occurring in at least 5 experiments, yielding 890,776 evaluable protein pairs. Edges were randomly split into training (75%) and test (25%) sets. The dataset exhibits extreme class imbalance characteristic of positive unlabeled (PU) learning: STRING contained 5,144 known interactions among 890,776 possible pairs (0.58%, 173:1 unlabeled-to-known ratio). The unlabeled set contains both true negatives and hidden positives (interacting pairs lacking robust experimental validation), necessitating specialized loss functions rather than treating all unlabeled pairs as negatives. Known positive edges from test and training sets were excluded from negative sampling to prevent information leakage.

We trained three model architectures, a naïve structure only model (Node2Vec), a variational graph autoencoder (VGAE), and a graph attention network (GAT). All models were trained on identical data splits and evaluated on the same held-out test edges. All models normalize predictions in the range [0,1]. For reference, model specific parameters are presented in the table below: Core Model Hyperparameters.

**Core Model Hyperparameters**

| Parameter | Node2Vec | VGAE | GAT |
|---|---|---|---|
| Embedding/Latent dim | 128 | 64 | 64 |
| Hidden channels | - | 64 | 64 |
| Attention heads | - | - | 4 |
| Training epochs | 20 | 50 | 30 |
| Learning rate | 0.025 | 0.001 | 0.01 |
| Dropout | - | 0.2 | 0.2 |
| Walk length | 20 | - | - |
| Context window | 10 | - | - |
| Neg samples per positive | 5 | - | - |
| Test Neg:Pos sampling ratio | - | 25:1 | 25:1 |
| KL weight | - | 0.00001 | - |
| Reconstruction weight | - | 1.0 | - |
| Loss function | Skip-gram | PU-BCE + KL | nnPU |
| Early stopping patience | 10 | 10 | 10 |

Node2Vec learns 128-dimensional node embeddings through biased random walks on graph structure without using node features. A single shared embedding matrix is maintained for all proteins and trained sequentially on each experiment graph, enabling cross-graph learning through shared parameters. Link prediction uses cosine similarity between node embeddings.

VGAE uses a two-layer graph convolutional encoder (64 hidden channels, 64 latent dimensions) with variational inference to learn latent node representations from both graph structure and node features. The model implements a hierarchical embedding architecture that separates stable protein identity from experiment-specific modulation: base embeddings capture protein-intrinsic properties shared across experiments, while experiment-specific embeddings capture condition-dependent variation. A learned attention mechanism determines how much each protein relies on its base identity versus experiment-specific context, enabling per-graph predictions. Training uses PU-adjusted weighted binary cross-entropy loss with a 25:1 unlabeled-to-positive negative sampling ratio.

GAT uses two-layer multi-head attention (4 heads, 64 hidden channels, 64 latent dimensions) to learn node representations with learned neighbor weighting. Like VGAE, GAT employs the hierarchical embedding architecture enabling per-graph predictions. Training uses non-negative PU (nnPU) loss[17]:

$$L_n nPU = \pi \cdot E_{x \in P}[\ell(f(x), 1)] + \max(0, E_{x \in U}[\ell(f(x), 0)] - \pi \cdot E_{x \in P}[\ell(f(x), 0)])$$

Where π is the class prior (estimated positive rate in the unlabeled set), P denotes the set of known positive pairs, U denotes the unlabeled set, f(x) is the model's predicted score for pair x, and ℓ is the binary cross-entropy loss. The first term penalizes incorrect predictions on known positives. The second term estimates the negative class risk by subtracting the expected loss on positives (weighted by the class prior) from the loss on unlabeled samples. The non-negative constraint (clamping values below zero to zero) prevents the negative risk from becoming negative, addressing a known failure mode of standard PU learning where overconfident predictions on unlabeled data can produce unbounded negative loss. Both models use curriculum learning that gradually increases the negative sampling ratio from 1:1 to the target 25:1 over the first epochs, allowing the model to first learn clear positive patterns before confronting the full class imbalance.

**Reference Interaction Databases**

To establish a reference interaction set for model evaluation, annotated protein interactions were compiled from four databases: STRING (comprehensive interactions integrating experimental and computational evidence), CORUM (manually curated protein complexes), IntAct (experimentally verified molecular interactions), and BioGRID (physical interactions only). All databases were filtered to retain protein pairs co-occurring at least 5 times across experiments. Dataset-specific filtering criteria are detailed in the table below.

Protein Interaction Datasets, Versions, and Filtering Criteria Used for Model Training and Evaluation

| Dataset | Version | Additional Filtering |
|---|---|---|
| String | 12.0 | High confidence interactions (score ≥ 0.7) |
| Corum | 5.1 | Not filtered |
| IntAct | Release 250 | Protein Pairs with miscore ≥ 0.4 |
| BioGrid | 4.4.245 | Physical interactions only |

**Model Evaluation**

Performance was evaluated using Recall, Average Precision (AP), ROC-AUC, and Precision@K (P@1000) on held-out test edges against each ground truth dataset. Given extreme class imbalance, Average Precision and Precision@K were prioritized over ROC-AUC. Evaluation used a 25:1 unlabeled-to-positive ratio, excluding both training and test positive edges from the negative sample pool. Performance is reported separately for each ground truth dataset to assess generalization across interaction types.

VGAE and GAT generate experiment-specific predictions by encoding each graph's unique node features and structural context through the hierarchical embedding system. For Node2Vec, per-graph metrics were computed by applying global predictions to each graph's test edges separately, then averaging across graphs. For cross-dataset evaluation, models trained exclusively on STRING interactions were evaluated on held-out interactions from CORUM, IntAct, and BioGRID (excluding any interactions present in STRING) to assess generalization. Note that each dataset has different inclusion criteria and different numbers of identified interactions.

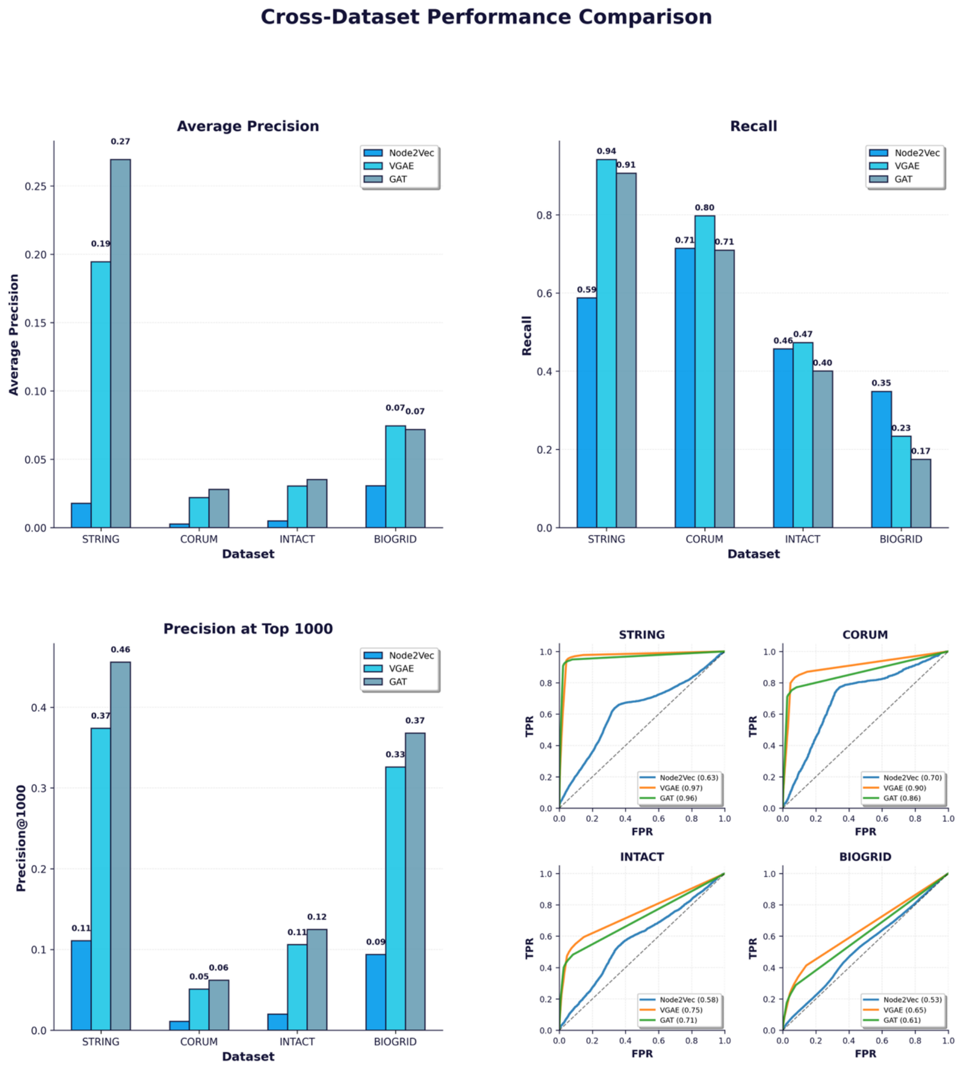

Above figure shows multi dataset model evaluation across key metrics**.** Cross dataset performance for our key metrics (Average precision, Recall, Precision@1000, AUC ROC) using the GAT model.

**Ablation Studies**
To isolate feature contributions, we conducted ablation studies across four configurations: structure only, proximity only, protein expression only, and combined features. Each configuration was trained independently with identical hyperparameters and evaluated on the same held-out test set (see Supplementary Figure 13).

**Thresholding Model Outputs**
For each experiment graph, we evaluated protein pairs where both proteins exhibited log2 fold change ≥ 0.3. A protein pair was considered a positive prediction if it received a score ≥ 0.75 from GAT or ≥ 0.5 from VGAE. Thresholds were selected based on the distribution of prediction scores, choosing values that distinguish high-confidence predictions from background signal. A protein pair was classified as a valid prediction if predicted in at least 5 different experiments. Prediction strength was quantified as the frequency with which a protein pair exhibited protein interaction-like behavior across experiments.

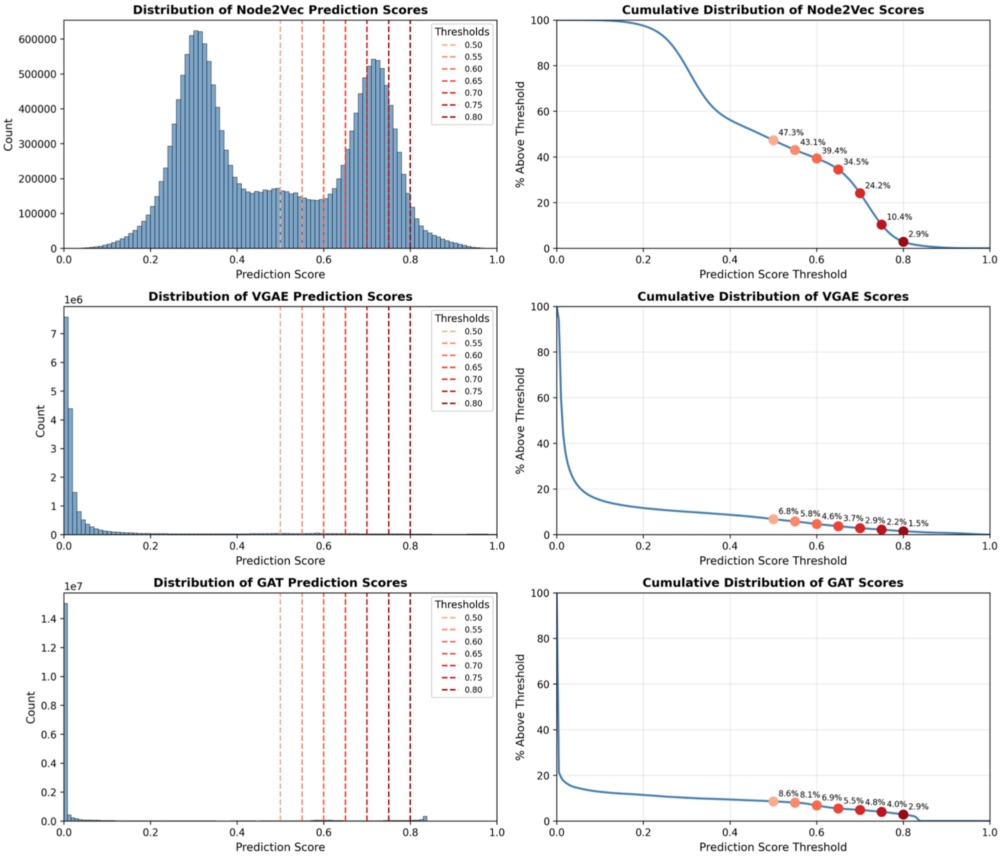

Above figure shows multi model prediction score distribution demonstrating prediction distribution of each ML model and how score thresholds were determined.

Because protein interaction annotations are incomplete, unlabeled protein pairs may include true but unobserved interactions, and model performance metrics represent lower bounds on true predictive accuracy. Models were trained on cell lines and conditions in the micromap dataset; generalization to other tissues or organisms has not been validated. High prediction scores indicate statistical association with annotated protein interaction patterns but do not establish direct physical interaction or biological mechanism.

**Contextualizing Model Predicted EGFR Interactions**

For GAT and VGAE models, we selected the top predicted EGFR proximal protein pairs by ranking candidates based on prediction frequency across micromap experiments, prioritizing pairs that were consistently predicted across multiple experimental contexts as indicators of robust interactions. For Node2Vec, which generates a single global embedding per protein rather than experiment-specific hierarchical embeddings, we ranked pairs by their prediction score and selected the top EGFR interactions.

When evaluating predicted protein pairs, we considered multiple complementary factors to assess biological relevance and therapeutic potential. We considered if the protein pair has a high confidence STRING interaction (score $\geq 0.7$). Therapeutic relevance is evaluated by determining whether each protein has an approved drug or is currently in clinical trials, as druggable targets offer immediate translational potential for combination therapy development. Expression patterns are examined across both tumor and normal tissues: tumor-normal differential expression from TCGA[18]/GTEx[19] analysis identifies proteins that are overexpressed in cancer (positive log2 fold change) and may represent tumor-selective targets, while baseline expression in normal tissues from GTEx provides safety context by revealing potential toxicity concerns in healthy organs.

**Differential Gene Expression Analysis**

Differential gene expression analysis was performed for each protein-indication pairing using DESeq2[20] implemented via pyDESeq2 (v0.4). Gene expression data were obtained from the UCSC Toil Recompute harmonized dataset[21], which reprocesses TCGA tumor samples and GTEx normal tissue samples through a unified RNA-seq pipeline to eliminate batch effects arising from different sequencing protocols.

For each indication (primary site), expression profiles from primary tumor samples (TCGA) were compared against normal tissue samples (GTEx). Statistical significance was assessed using the Wald test. P-values were adjusted for multiple testing using the Benjamini-Hochberg procedure to control the false discovery rate (FDR) at $\alpha = 0.05$. A minimum of 3 samples per group was required for analysis.

Results are reported as $\log_2$ fold change (tumor versus normal) with corresponding adjusted p-values.

**Reactome Pathway Analysis**

Additionally, we use Reactome[22] pathway analysis functionality to identify biological pathway overlap between predicted interacting proteins. For each protein pair, we submit the overlapping predicted partners to Reactome's enrichment analysis to identify shared pathway memberships, retaining only pathways with Reactome Entities FDR < 0.05. When evaluating the proportion of pathway coverage, we consider only proteins from the possible pathway entities.

**Proteomic Data Commons**

Clinical proteomics data were retrieved from the Proteomics Data Commons (PDC)[23] and from Liu et al.[24] by ingesting all studies that contained a proteomics file in their Protein Assembly. From these studies, we selected relevant cancer indications for analysis. Gene symbols from each proteomics file were mapped to UniProt[25] accessions by first attempting to match against UniProt primary symbols, when no primary symbol match was found, we fell back to matching gene symbols to protein synonyms to maximize coverage.

**Software Environment**

Subsequent analyses were performed using Python 3.10+ with the following core packages: NumPy (≥1.26), pandas (≥2.1), SciPy (≥1.11), NetworkX (≥3.2), Numba (≥0.60), and umap-learn (≥0.5.9). Visualizations were generated using Matplotlib (≥3.8) and Seaborn (≥0.13.2). Data were stored in Apache Parquet format via PyArrow (≥15.0).

Graph ML Models were implemented in Python 3.12 using PyTorch (v2.7.0), PyTorch Geometric (v2.6.1), NumPy (v2.2.6), and scikit-learn (v1.6.1). All machine learning models were trained on an AWS g4dn.xlarge instance equipped with a single NVIDIA T4 GPU, 4 vCPUs, and 16 GB of RAM.

### Cell starvation and EGF stimulation for microenvironment mapping

CAOV3 cells were grown in manufacturer's recommended medium. Two sets of cells were washed twice with DPBS and starved in medium lacking FBS for 24 hours. The media was replaced in both sets with one of the sets supplemented with 30 ng/mL EGF (Thermo Scientific, Cat: PHG0311L) for an additional 24 hours. Cells were then harvested for western blot analysis and micromapping.

For western blot analysis, equal numbers of cells were lysed in RIPA with 1X protease inhibitor tablet and 1:1000 dilution of benzonase (Sigma-Aldrich, Cat: 70664-3). Protein concentrations were measured by BCA assay and normalized. Samples were mixed with 4X Laemmli buffer (Biorad, Cat: 1610747) with ß-mercaptoethanol (Thermo Scientific, Cat: AC125472500) and stored at -80$^{0}$C. Before loading, samples were boiled at 95°C 5 min and run 1 hour at 180V on a 12% TGX Criterion gel (Biorad, Cat: 5671045). The iBright Prestained Protein Ladder (Thermo Scientific, Cat: LC5615) was included as a protein ladder. Gel was washed briefly in water before transfer using an iBlot2 Gel Transfer device and PVDF stacks (Thermo Scientific, Cat: IB24001). PVDF blots were blocked in 3% bovine serum albumin (Sigma, Cat: A7906-100G) in 1X TBST (diluted from 20X stock, Boston BioProducts, Cat: IBB-181X-4L) for at least 1 hour. Blocking buffer was replaced and primary antibodies for EGFR (1:1000 dilution, Cell Signaling Technology, Cat: 4267S), CDCP1 (1:1000 dilution, Cell Signaling Technology, Cat: 4115S), and ß-actin (1:5000 dilution, Thermo Scientific, Cat: MA5-15739) were added. The next day, blots were washed three times 5 min each with 1X TBST before addition of goat anti-mouse IRDye 680RD (1:5000 dilution, LICORbio, Cat: 926-68070) and goat anti-rabbit IRDye 800CW (1:5000 dilution, LICORbio, Cat: 926-32211) in blocking buffer for 1 hour. Blots were washed three times 5 min each with 1X TBST and two times briefly with water before imaging on a Licor Odyssey CLx.

### Internalization, ADC cytotoxicity, and TCE cytotoxicity assays

For internalization assays, cells (SW48, BXPC3, PC3, bronchial/tracheal epithelial cells and cervical epithelial cells) were grown in manufacturer's recommended media and were harvested and plated at 20,000 cells per well in a tissue culture treated 96 well plate. Cells were allowed to adhere at 37°C overnight. The following day, test and control antibodies were prepared in a 3:1 molar ratio of Incucyte Human Fabfluor-pH Antibody Labeling Dye (Sartorius, Cat. #4722). Labeling reactions were incubated in the dark for 15 min at 37 °C. Immediately following incubation, 50 µL of 2X labeled antibody solution was added to each well containing 50 µL of culture medium to achieve the final concentration (2.5nM for SW48, 10nM for all other cell lines). Plates were transferred to an Incucyte S3/SX1 live-cell imaging system (Sartorius) and imaged every 45 minutes for 24 h in both phase contrast and red fluorescence channels (10× objective, 400 ms exposure). Imaging began 30 min after placement in the incubator to allow condensation to dissipate. Images were analyzed using Incucyte software (v2023A). Phase segmentation was performed using the AI Confluence default settings, while red fluorescence was analyzed using Top-Hat segmentation (radius = 30 µm; threshold = 0.2 RCU). Total red integrated intensity per well was normalized to phase confluence and exported for visualization in GraphPad Prism (v10.4.1).

For cytotoxicity assays, cells (SW48, BXPC3, PC3, bronchial/tracheal epithelial cells and cervical epithelial cells) were harvested and plated in their supplier's recommended media at 300 cells/well in 45 µL/well in tissue culture treated white 384-well plates (Corning 3570). Cells were allowed to adhere at 37°C overnight, and then an ADC dilution plate was prepared by making serial dilutions of 10x concentrated ADCs in PBS in a 96-well plate. An automated liquid handler was used to move 5 8µL of these ADCs into each well of the 384 well assay plates to get n=4 of each condition. Plates were returned to the incubator for five days. To read out cell viability, 10 µL of CellTiter-Glo (Promega) was added to each well, plates were incubated for 30 minutes at room temperature, and endpoint luminescence was read out using a plate reader. To calculate viability, all luminescence values were normalized to the average of untreated wells (antibody concentration of 0) for a given plate and cell type.

To measure T cell dependent cellular cytotoxicity, selected cell lines (SW48, HCC827 and PC3) were stably infected with a lentiviral construct expressing a codon optimized firefly luciferase driven by EF1α and selected with blasticidin to generate cell lines that could be monitored for viability. For the assay, luciferized cells were harvested and plated in their supplier's recommended media at 20,000 cells/well in 96-well tissue culture treated plates. That same day, PBMCs were thawed and put in culture in RPMI media with 10 ng/mL IL-12 (Gibco 200-12H). Both types of cells were incubated at 37°C overnight. The following day, PBMCs were harvested, counted, and plated in fresh media with no IL-12 at 200,000 cells/well in the assay plate with the tumor cells. TCE antibodies were added in triplicate and assay plates were returned to the incubator for two days, then read out by adding an equal volume of Steady-Glo (Promega) and reading out on a luminescence plate reader. Luminescence signal was normalized to wells with PBMCs but no antibodies added to determine tumor cell cytotoxicity.

### Generation of SW48 CDCP1 Knockdown Cell Line

SW48 CDCP1 knockdown cells were generated using CRISPR–Cas9–mediated gene disruption. A synthetic dual–nuclear localization signal (sNLS)–SpCas9 nuclease (Aldevron) was combined with a pooled set of CDCP1-targeting CRISPR guide RNAs obtained from Synthego. Guide selection followed the manufacturer's design criteria for targeting early coding exons to promote functional knockdown. Ribonucleoprotein (RNP) complexes were prepared following general guidelines provided by Integrated DNA Technologies (IDT) for Cas9–gRNA complex assembly. After treatment of SW48 parental cells with the assembled complexes, cells were expanded under standard culture conditions to allow outgrowth of edited populations. Following recovery, bulk-edited cultures were screened for loss of CDCP1 surface expression by flow cytometry. CDCP1 knockdown cells were enriched by fluorescence activated cell sorting (BD FACSAria), ensuring a stable CDCP1-knockdown cell line suitable for downstream analysis.

### In vivo cell line-derived xenograft (CDX) studies

Female CrTac:NCr-Foxn1<nu> (NCr nude) or NOD.Cg-Prkdcscid Il2rgtm1Sug/JicTac (NOG) mice, age 5 to 8 weeks, were obtained from Taconic Biosciences. All mice were housed up to 5 mice per cage in individually ventilated cages in a pathogen-free animal facility. Mice were maintained under artificial lighting (12 hours) in a controlled ambient temperature of 68 to 79°F, and relative humidity between 30 and 70%. Mice were acclimated for at least 5 days before the experiments. All animal procedures were conducted in accordance with, and with approval of, the policy of the Bloodworks NW Research Institutional Animal Care and Use Committee.

Tumor cells (2 to $10\times10^6$ / 100 µL) were implanted subcutaneously into the right hind flank of female mice in a 1:1 mixture of base media and matrigel (Corning, Cat: 356234). In the dual-flank model, parental tumor cells were implanted in the right hind flank and CDCP1 knockdown tumor cells were implanted in the contralateral flank.

Tumors were measured by digital calipers, and tumor volume (TV) was calculated as $TV = (L\times W\times W)/2$, where L (length) is the longest measurement (in mm) and W (width) is perpendicular to L. When tumors reached ~100-150 $mm^3$, mice were randomized by TV into study groups (n=8-10 mice per group).

The ADCs were administered at the indicated dose in 100 µL formulation buffer (20 mM histidine, 8% sucrose, pH 5.5) via tail vein injection. Tumors and body weight (BW) were measured twice weekly until control-treated tumors reached 2000 $mm^3$, at which point tumor growth inhibition (TGI) and mean BW change were evaluated. TGI was calculated as (1 – [TVfinal, treated – TVinitial, treated / TVfinal, control – TVinitial, control]) x 100. TGI statistics were performed on GraphPad Prism using a two-way ANOVA with Tukey's all groups comparison. Animals were euthanized early if they lost >20% of their initial BW, had severe tumor ulceration, or became moribund.

Treated mice continued to be monitored for long-term responses until Day 80, or when tumors reached 800 $mm^3$ (survival endpoint). A Kaplan-Meier analysis was performed in which a survival event was considered a tumor volume exceeding 800 $mm^3$. Survival statistics were performed on

GraphPad Prism using logrank (Mantel-Cox) test. P values of less than 0.05 were considered significant (*, $p<0.05$; **, $p<0.01$; ***, $p<0.001$; ****, $p<0.0001$).

### CDX tumor dissociation for photocatalytic proximity labeling

Female NCr nude mice were implanted with SW48 tumor cells as described above. When tumors reached ~500 $mm^3$, tumors were excised with scissors and forceps, placed in RPMI-1640 media (Gibco, Cat: A10491-01) and kept on ice. Single cell suspensions of SW48 tumor cells were prepared by chopping tumor tissue with a razor blade and digesting with an enzyme cocktail supplied in Miltenyi's Tumor Dissociation Kit, Human (Cat: 130-095-929). Approximately 500 mg of tumor tissue was incubated with enzyme cocktail in gentleMACS C tubes (Miltenyi, Cat: 130-096-334) on a gentleMACS™ Dissociator (Miltenyi) following the manufacturer's protocol. Cell suspensions were washed with RPMI-1640 media, passed through a 70 µm MACS® SmartStrainer (Miltenyi, Cat: 130-110-916), washed with PBS and kept on ice. Viable cell counts were obtained, and cells were frozen in CryoStor® CS10 media (Stem Cell Technologies, Cat: 07930) until use.

### Terminal Tissue Collection and Immunohistochemistry

Tumors from untreated mice were dissected with skin attached and fixed in 10% neutral buffered formalin (Sigma-Aldrich, Cat: HT501128) for 24 hours, then transferred to 70% ethanol (Decon Labs, Inc., Cat: 8601). Tumors were paraffin embedded, sectioned, and analyzed by immunohistochemistry (IHC) at Acepix Biosciences (Union City, CA) using validated IHC assays for CDCP1 and EGFR. Specimens were embedded tumor side down (excess skin was trimmed if needed). If the tumor was large, it was bisected from the center of the tumor and both pieces were embedded on the same block cut side down so sections were collected from the center of the tumor. Specimens were incubated with anti-CDCP1 (Cell Signaling Technologies, Cat: CST4115) or anti-EGFR (Abcam, Cat: ab227642) for 1 hour at 0.42 and 0.0667 mg/mL, respectively.

### Antibody and ADC production

EGFR x CDCP1 bispecifics, TCE trispecifics, and their monovalent controls were transfected in ExpiCHO-S cells (Thermo Scientific, Cat: A29127) in ExpiCHO expression medium (Thermo Scientific, Cat: A2910001). Proteins were purified on a Cytiva ÄKTA Pure. All samples were first purified using PrismA HiTrap affinity capture (Cytiva, Cat: 17549854), neutralized with 1M NaPi pH 7.0, and polished by SEC purification over a Superdex 200 Increase column (Cytiva, Cat: 28990944) in PBS (Fisher Scientific, Cat: SH3025602).

All constructs were generated using knob-in-hole approach[26]. These molecules contain CH3 mutations that preferentially drive heavy chain heterodimer formation, and light chain swapping is not a concern because the molecules are asymmetric, containing only one FAB arm.
Purity was determined by analytical SEC on an Agilent 1260 Bioinert system using an AdvanceBio SEC 300Å, 4.6 x 150mm, 2.7µm LC column (Agilent, Cat: PL1580-3301) with an AdvanceBio SEC 300A, 4.6 x 50mm, 2.7µm guard column (Agilent, Cat: PL1580-1301).

Interchain conjugate DAR was determined by analytical HIC on an Agilent 1260 Bioinert system using an AdvanceBio HIC, 4.6 X 100 mm (Agilent, Cat: 685975-908). A 20 minute gradient was used to elute the product from 1.2 M ammonium sulfate (Thermofisher, Cat: J64419.A3), 50 mM NaPi pH7.0 (Thermofisher, Cat: J63791.AP) to 50 mM NaPi, 20% Isopropanol (Thermofisher, Cat: 383910025).

Identity and drug antibody ratio, DAR, analysis were performed on a Waters Bioaccord system using a BioResolve RP mAb Polyphenyl Column (Waters, Cat: 186008945). Binding functionality was performed using a Sartorius Octet BLI (Sartorius, RH16) by capturing antibodies to AHC2 sensors (Sartorius, Cat: 18-5142) and incubating in target proteins of interest; EGFR, CDCP1 and CD3. (Acrobiosystems, Cat: EGRH5222, CD1H52H6, CDEH5223 respectively)

Conjugates utilizing engineered cysteines were generated following a protocol similar to that previously described for site-specific Thiomab™ conjugation[27]. Antibodies and bispecifics were first buffer exchanged into 100 mM Tris, 1 mM EDTA pH 8.0 at concentrations above 5mg/ml. 80 molar equivalents of DTT (Thermo Scientific, Cat: A39255) was added to fully reduce interchain disulfides and engineered cysteines, either 3 hours at room temp or overnight at 4°C. The product was buffer exchanged into 20 mM Tris pH 7.5 using Zeba desalting columns (Thermo Scientific, Cat: 89892) to remove DTT and release cysteine and glutathione.

15 molar equivalents of DHAA (Thermo Scientific, Cat: 250930050) was added and incubated at room temperature for one hour or until interchain disulfides reform. DHAA was removed by desalting into PBS using a Zeba column. 10% v/v DMSO (Sigma, Cat: D2653) was added to prepare for the linker-payload addition. Maleimide-VC-PAB-MMAE (Medchemexpress, Cat: HY-15575) was solubilized in DMSO at 5 mM. 4 molar equivalents of maleimide-VC-PAB-MMAE were added and incubated for at least one hour at room temperature or overnight at 4°C.

Material was run over a preparative SEC column, Superdex 200 Increase, to remove aggregates and free linker-payload. The final material was buffer exchanged and concentrated using Amicon Ultra concentrators (Millipore, Cat: UFC903024) to remove remaining free linker-payload, formulate it into the appropriate buffer, and reach the desired concentration.

For non-site-specific conjugates utilizing interchain disulfides, the antibodies and bispecifics were buffer exchanged into 100 mM Hepes pH 7.0 using Zeba columns. 3 molar equivalences of TCEP (Thermo Scientific, Cat: 77720) were added to the solution and mixed for 90 minutes at 25°C with mixing. 10% v/v DMSO was added to the solution and mixed. Maleimide-VC-PAB-MMAE was solubilized in DMSO at 5 mM. 6 molar equivalences of maleimide-VC-PAB-MMAE were added and mixed at room temperature for at least 60 minutes. Material was run over a preparative SEC column, Superdex 200 increase to remove aggregates and free linker-payload. The final material was buffer exchanged and concentrated using Amicon ultra to also remove free linker-payload, formulate it into the appropriate buffer, and reach the desired concentration.

### Characterization data of site-specific engineered cysteine bispecific ADCs

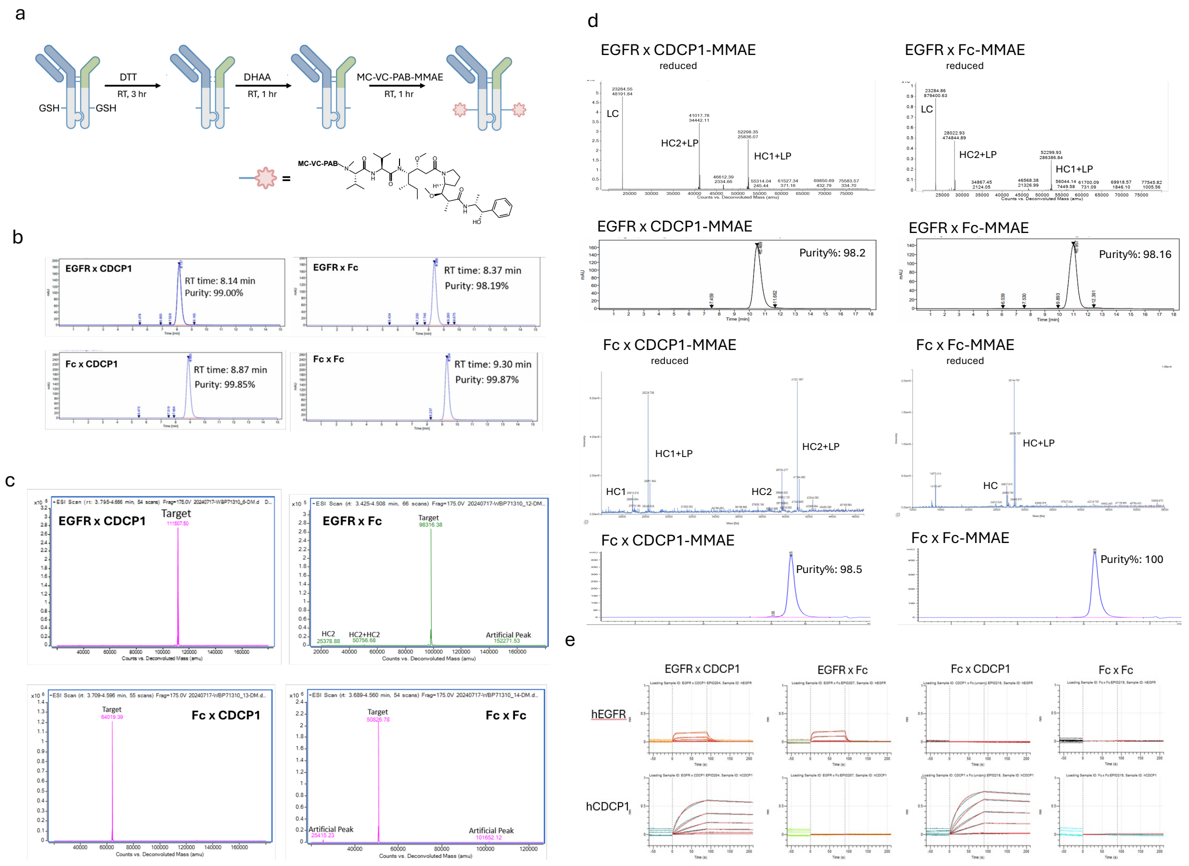

Figure above shows **a)** Outline of the general process flow for site specific conjugation of engineered cysteines. **b)** Analytical SEC chromatograms of the bispecifics used in-vitro. **c)** Deconvoluted, intact mass spectrometry traces of bispecifics used in-vitro. **d)** Analytical SEC chromatograms and deconvoluted, reduced traces of conjugated bispecifics used in vitro and in-vivo **e)** Sensorgrams of each binding arm of bispecifics used in-vivo.

### Characterization data of TCE trispecifics

a

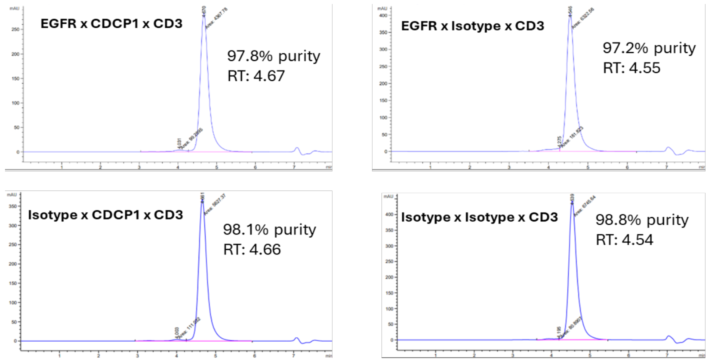

b

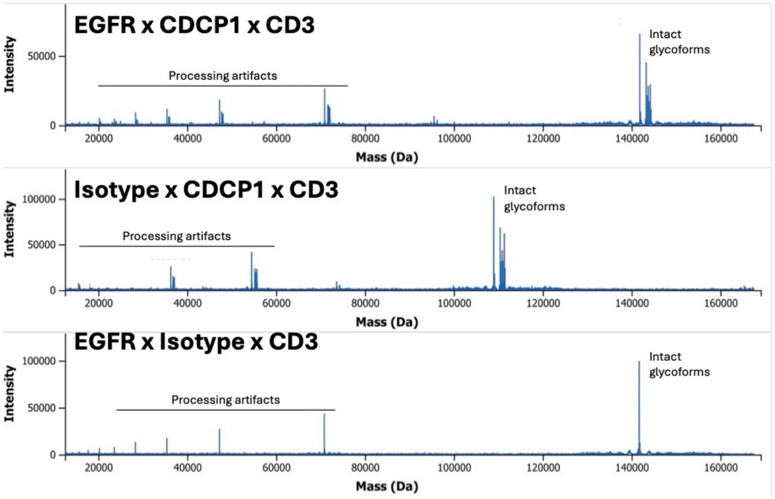

c

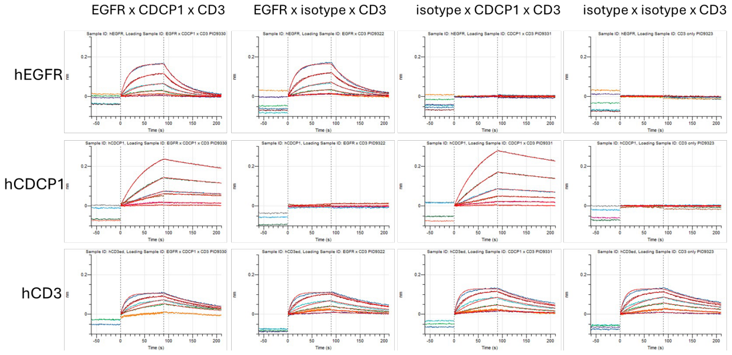

Figure above shows **a)** Analytical SEC chromatograms of TCE trispecifics used in-vitro. **b)** Deconvoluted, intact mass spectrometry traces of TCE trispecifics. **c)** Sensorgrams of each binding arm of TCE trispecifics.

**Characterization data of interchain disulfide bispecific ADCs**

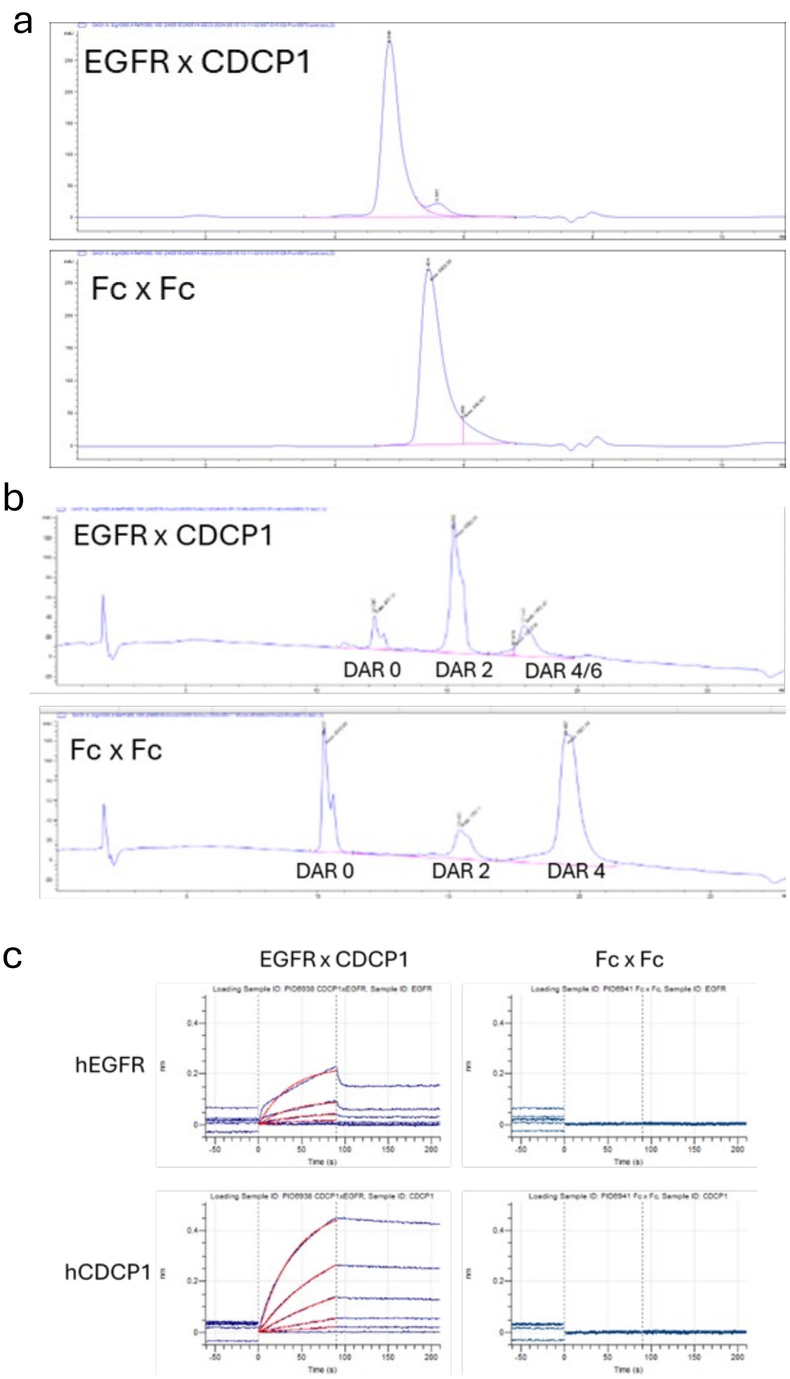

Figure above shows **a)** Analytical SEC profiles of ADCs **b)** HIC chromatograms of ADCs **c)** Binding profiles of the arms used in-vivo.
